# Challenges and lessons learned from preliminary modeling of with-in herd transmission of highly pathogenic avian influenza H_5_N_1_ in dairy cattle

**DOI:** 10.1101/2024.08.06.606397

**Authors:** Brinkley Raynor Bellotti, Michael E. DeWitt, Jennifer J. Wenner, Jason E. Lombard, Brian J. McCluskey, Nicholas Kortessis

## Abstract

The emergence of highly pathogenic avian influenza A H_5_N_1_ in dairy cattle raises many questions related to animal health and changes to the risk of an epidemic in humans. We synthesized information currently published to fit a compartment model of H_5_N_1_ transmission within a dairy herd. An accompanying web application allows users to run simulations for specific outbreak scenarios. We estimated *R_0_* near 1.2 with a short duration of infectiousness and fast time course of an epidemic within a farm, which we discuss in the context of possible on-farm control strategies. The web application allows users to simulate consequences of an epidemic using herd-specific information, a tool we propose will help inform stakeholders about potential consequences of uncontrolled H_5_N_1_ spread. Our modeling work has identified several key information gaps that would strengthen our understanding and control of this emerging infectious disease.

## Background

The emergence of highly pathogenic avian influenza (HPAI) A in cattle is a striking development in the history of this pathogen, which in the past two and a half years has spread to three new continents and affected many new hosts species (1–4). On March 25, 2024, HPAI A H_5_N_1_ Clade 2.3.4.4b (H_5_N_1_) was first diagnosed in dairy cattle by the United States Department of Agriculture (USDA) in samples from Texas (1). Analysis of viral genomes indicates that the first cattle infection likely occurred in early 2024 and that the current outbreak is the result of sustained cow-to-cow transmission (5,6). As of August 1, 2024, 175 dairy herds have been reported to be affected in 13 states (Figure 1). Cows infected with H_5_N_1_ have been reported to exhibit lower milk production, with some proportion never recovering to pre-infection production levels (7,8). Dairy cattle tend to be intensively managed with high human contact. Previous H_5_N_1_ outbreaks in humans have had high case fatality rates (9,10), but only mild cases have been observed in the United States (11–14). The pandemic potential of this emerging pathogen as it continues to expand and evolve is yet unknown. Recent H_5_N_1_ spillover into cattle has the potential to be highly disruptive to the dairy industry, the economic livelihood of farmers, and human health.

**Figure 1:**
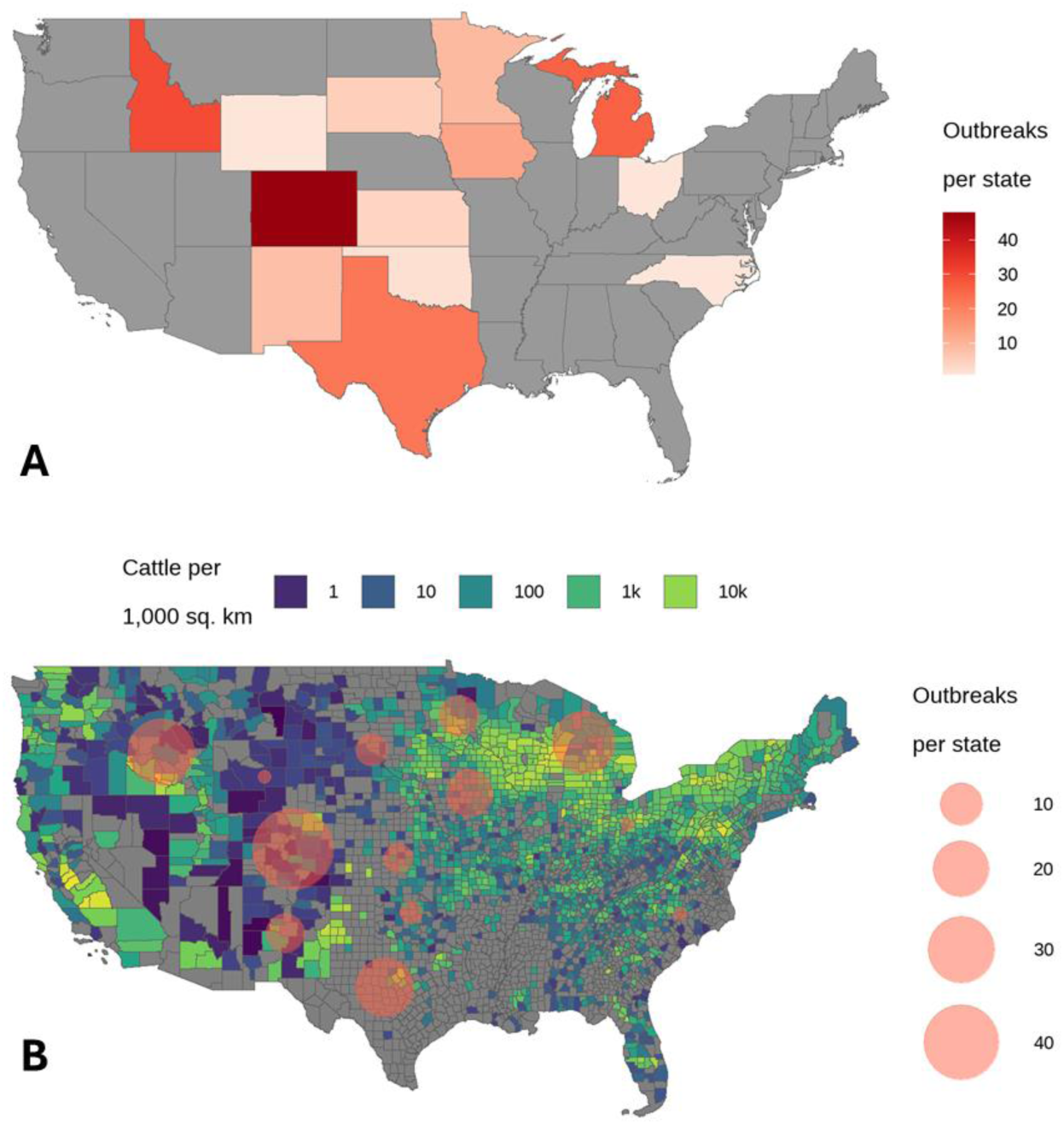
Map of reported H_5_N_1_outbreaks in dairy herds. Panel A represents the outbreak size (number of infected farms) per state that have been reported to APHIS. Panel B displays the outbreak size per state overlaid against the underlying dairy cattle density. The shapefiles used for mapping are the 2022 state files at a 1:500,000 resolution and the 2022 county files at a 1:500,000 resolution (15) and agricultural data on dairy cattle operations are available through the National Agricultural Statistics Service (NASS) 2017 agricultural census accessed through the NASS Quick Stats tool using the data item “cattle, cows, milk – inventory” at the county level (16). Maps last updated with cases on August 1, 2024.

Emerging infectious diseases present challenges to animal and public health officials, medical professionals, veterinarians, producers, and industry stakeholders working with limited information to mitigate onward transmission. Many interventions rely on disrupting transmission chains; understanding the way a pathogen spreads through a population is critical to formulating effective control measures. Mathematical models of pathogen transmission are powerful tools to infer important factors related to transmission given available data, to evaluate alternate intervention scenarios, and to identify key gaps in knowledge related to the determinants of pathogen spread. In this report, we incorporate currently available data about the ongoing H_5_N_1_ outbreaks in dairy cattle into a model of within-farm transmission to establish a framework for characterizing H_5_N_1_ outbreaks.

## Methods

### Review of published data and outbreak settings

Dairy cattle in modern operations are housed in large groups with the ability to directly contact and interact with other individuals in the group and between groups. Some spaces are utilized by all groups, such as the milking parlor. Milking equipment has been implicated in transmission between cows (17). Other transmission pathways have been implicated including aerosolization (7,8,18). Studies have found large viral loads shed through milk (17–20). Though initial investigations have detected viral shedding in milk rapidly after exposure (18,21), some clinical signs (e.g., drop in milk production, decreased rumination) may take several days to manifest (18,22). Clinical pictures vary between infected individuals, ranging from no clinical signs to death/euthanasia (7). Decreased milk production has been reported to persist past other clinical signs, with some cows never fully recovering to previous production levels (7,18,22). A preliminary study indicates milk production decreasing to a nadir of 15-30% during the infection period and improving to 71-77% of pre-infection baseline at the end of the study period; on histopathology, fibrosis of mammary gland tissue suggested permanent changes in milk production capacity after infection (18). Economic considerations have led to cows with long-term decreased milk production being sent to market and replaced (7,22). Ongoing investigations found outbreaks within a herd to last approximately a month with the peak infection being approximately two weeks (7,22). Responses have varied by state, and different clinical manifestations have hindered development of a single, clear clinical case definition leaving open the possibility for subclinical and undetected infections in cattle (7).

### Model structure

The model describes short term dynamics within a farm during an outbreak (typically no longer than two months) based on published data and case reports (Figure 2). The model is similar to an Susceptible-Infectious-Recovered (**SIR**) model, but includes three additional compartments: a transitional compartment **B** of duration 1/*α* where cows display clinical signs, such as decreased milk production (7,18), but are not shedding sufficient virus to be considered infectious, a compartment **D** that includes cows that die or are euthanized due to disease, and a compartment **C** that represents cows that do not sufficiently recover from disease and are elected to be sent to market (culled). The model assumes density-dependent transmission between susceptible, **S,** and infectious, **I**, individuals such that susceptible cows are infected at per-capita rate *β***I**, and that that infectious individuals stay infectious for a duration 1/*γ*. We assume a fraction *ϕ* of infectious individuals die due to disease, and a fraction *ρ* of surviving clinically affected individuals fail to recover from disease and are sent to market. We also assume recovery to compartment **R** following infection leads to persistent immunity. While information about immunity is not yet available, the duration of epidemics is sufficiently short that we assume immunity persists over this time period. With this **SIBR** model structure, we can estimate milk production at each compartment to account for the economic impact of the reported decreased milk production. We estimate loss of milk production with a 50% drop during clinical illness (**I**, **B** compartments) and 25% drop once recovered (**R** compartment) (18). We considered an alternate model structure that included an exposed compartment **E;** however, the joint estimates of incubation duration with other parameters were sufficiently short (approximately an hour) to support a model framework more consistent with a classic SIR structure (Appendix Figure 1). The model is flexible to allow background population turnover, *μ* (cows replaced at end of productive lifespan, often between 2.5-4 years (23)), but for purposes of fitting the model, we have assumed background turnover to be zero during the outbreak period. The corresponding set of ordinary differential equations (1.1 – 1.6) is in the technical appendix, with state variables representing disease state compartments (Figure 2) and parameters summarized in Table 1.

**Figure 2:**
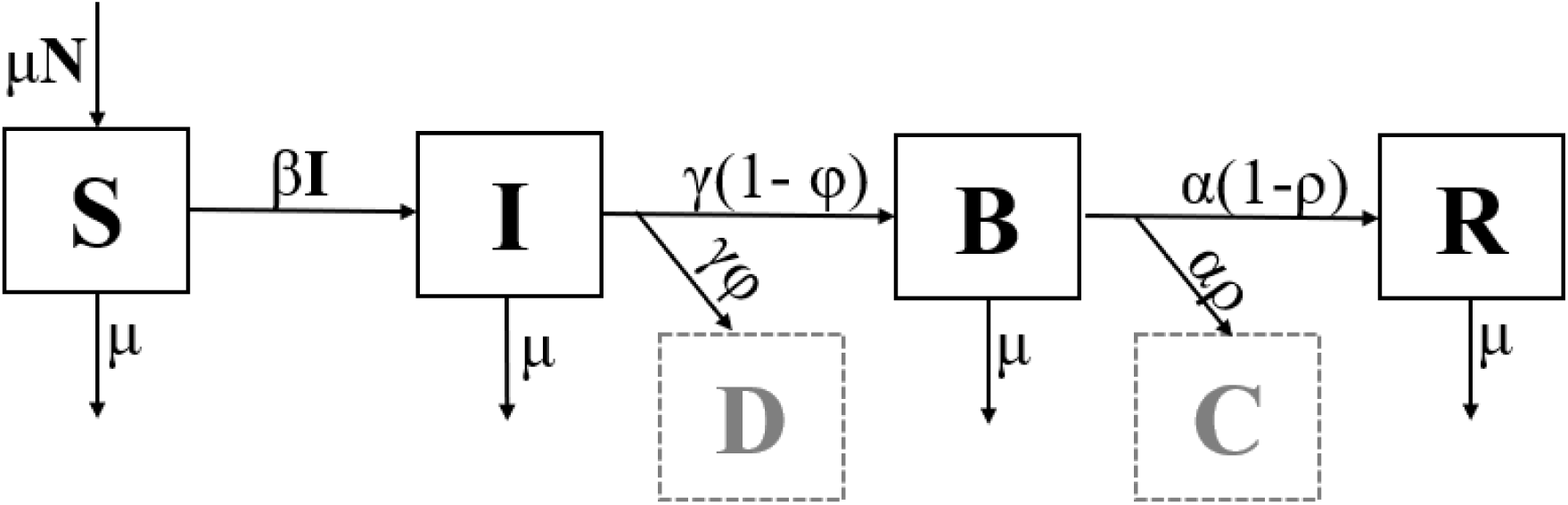
A flow diagram of the within herd HPAI model structure. The compartments represent disease states: **S** – susceptible, **I** – infectious, **B** – transitional state where no longer infectious but still clinically ill, **R** – recovered with immunity, **D** – died due to disease, and **C** – never fully recovered and sent to market. The arrows going in and out of the boxes represent the rates at which individuals move in and out of that disease state. **N** is the total numbers of living cows in the herd.

**Table 1.**
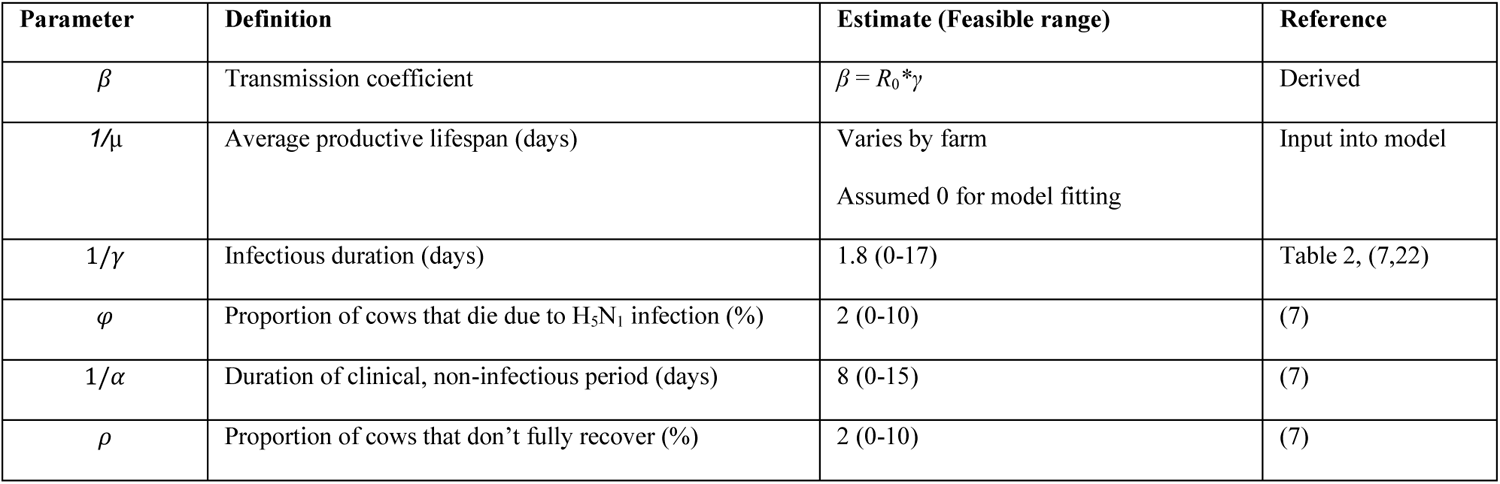
Parameters for within-herd H_5_N_1_ outbreak model.

| Parameter | Definition | Estimate (Feasible range) | Reference |
| --- | --- | --- | --- |
| $\beta$ | Transmission coefficient | $\beta = R_0 * \gamma$ | Derived |
| $1/\mu$ | Average productive lifespan (days) | Varies by farm<br>Assumed 0 for model fitting | Input into model |
| $1/\gamma$ | Infectious duration (days) | 1.8 (0-17) | Table 2, (7,22) |
| $\varphi$ | Proportion of cows that die due to H5N1 infection (%) | 2 (0-10) | (7) |
| $1/\alpha$ | Duration of clinical, non-infectious period (days) | 8 (0-15) | (7) |
| $\rho$ | Proportion of cows that don't fully recover (%) | 2 (0-10) | (7) |

### Parameterization

An important determinant of disease dynamics is the basic reproductive number, *R*_0_, defined as the average number of secondary cases a single index infectious individual will cause in a completely susceptible population. For this disease system, *R*_0_ = *β*/*γ*. In this model, *R*_0_ determines the number of individuals infected during an epidemic. Given *R*_0_, the peak of the infection occurs when the susceptible population is at the herd immunity threshold, *p_c_* = 1 – 1/*R*_0_, i.e., the critical proportion of herd immunity that is required to disrupt transmission (24). Moreover, *R*_0_ is related to the final infected fraction as 1 – **S**(∞) = 1 – exp{–*R*_0_(1 – **S**(∞))}. The time course of an epidemic is mainly determined by the infectious duration, 1/*γ.* We use data on the time to peak infection and the duration of the epidemic to estimate *γ*. Because the transmission coefficient, *β*, is not directly observed, we calculate the transmission coefficient given estimates of *R*_0_ and *γ* as *β* = *R*_0_*\*γ*.

Estimation was done using a stochastic version of the model. Based on biologically feasible ranges of parameters reported in the literature (Table 1), we ran model simulations over the entire parameter space for 500 iterations for each parameter combination for a total of 5 million stochastic realizations. The resulting simulations were subset to those iterations and parameter combinations which reproduced outbreaks durations and attack rates consistent with publicly reported data (7,8,22). This included those outbreaks with a peak incidence of clinical disease between days 7 and 21 with a final fraction infected between 30% and 80% with the duration of outbreak between 20 and 60 days (7,8,22,25). We employed a two-dimensional kernel density function with an axis-aligned bivariate normal kernel to estimate the joint values of *R*_0_ and infectious duration, *γ*, using those iterations which fit the outbreak phenomena.

### Uncertainty

We used a resampling approach to indicate a range of parameter values consistent with the minimal available data and the model structure. The scenarios that met the inclusion criteria for the outbreak were resampled with replacement. The two-dimensional kernel density function was estimated with the highest density parameter estimates extracted. We repeated this resampling procedure 500 times and calculated the 2.5% and 97.5% quantiles of these estimates.

### Data and computation

Data used in analyses come from publicly available sources. Case data at the farm level are available on a public dashboard hosted through the Animal and Plant Health Inspection Service (APHIS) (26). Spatial data in the form of cartographic boundary shapefiles are from the United States Census Bureau (15). All other parameter estimates come from the scientific literature published to date (Table 1). All analyses were performed using R version 4.3.1 (27). Code used to perform analyses and model transmission are available in the Appendix (Appendix Code 1, Appendix Code 2). This study was approved by the institutional review board of Wake Forest University School of Medicine (#IRB00117110).

## Results

### Outbreak simulation

Based on our resampling estimates for the joint values of *R*_0_ and infectious duration, *R*_0_ values between 1.15-1.25 are consistent with the data, with a most likely value of 1.2, and infectious duration parameters between 1.1-1.3 days are consistent with the data, with a most likely value of 1.15 (Table 2). This value of *R*_0_ predicts that up to 31% of the herd will be infected during an epidemic (Table 2, Figure 3) and that 17% of the herd must gain immunity to reach the herd immunity (Table 2). For our simulations, we did not include population turnover during the duration of the outbreak; however, population turnover would replace recovered individuals with susceptible individuals in a herd previously at a threshold herd immunity level leading to recurrent outbreaks depending on the rate of population turnover (Appendix Figure 2).

**Figure 3:**
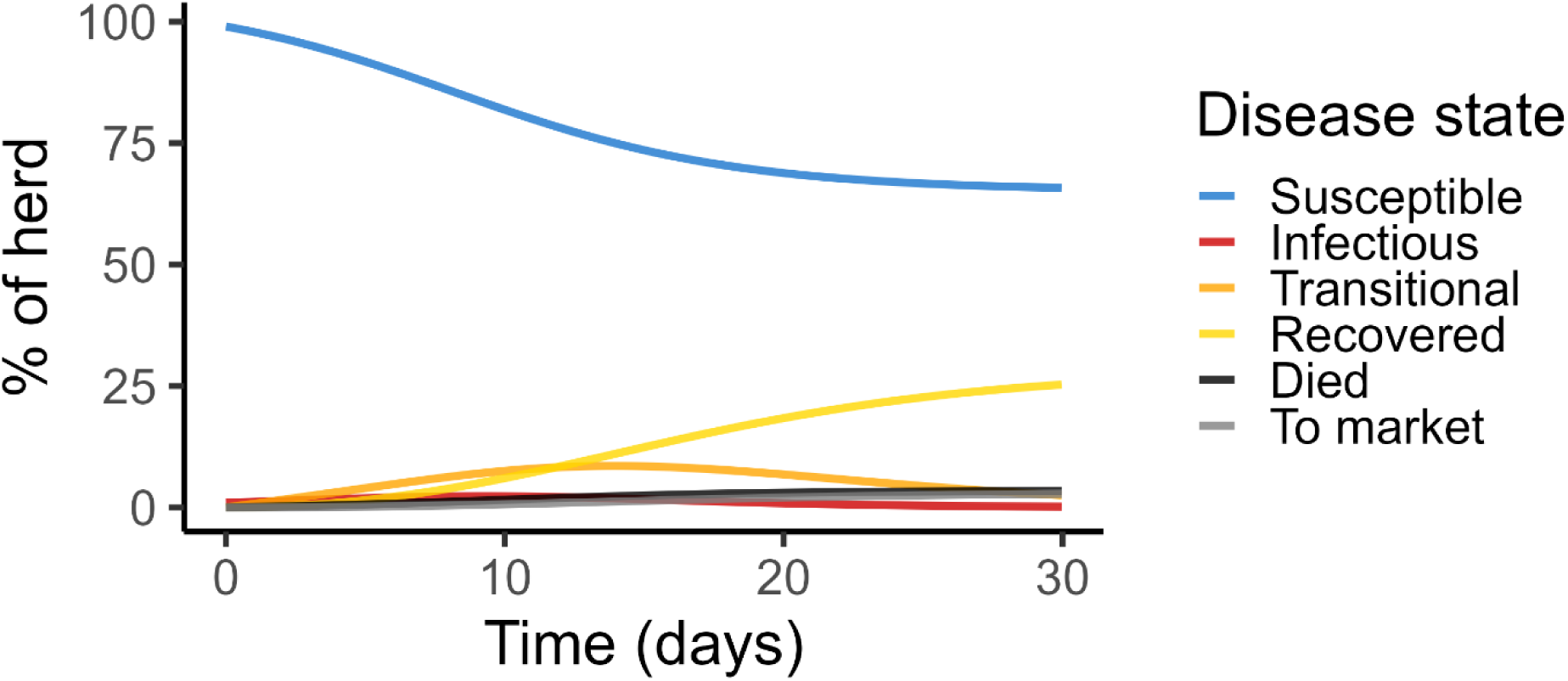
Disease dynamics with best fit parameterization (Table 2). Consistent with case reports and early epidemiologic data (5,7,8,18,22), the infection peak occurs within 2 weeks of clinical signs, and the outbreak course is rapid with the epidemic being effectively over within 4 weeks.

**Table 2.** Derived parameter estimates.

| Parameter | Estimate (95% Sampling Interval) |
| --- | --- |
| Basic reproduction number, $R_0$ | 1.2 (1.15 - 1.25) |
| Infectious duration (days) | 1.15 (1.09 - 1.28) |
| Final fraction infected | 0.31 (0.25-0.37) |
| Herd immunity threshold, $p_c$ | 0.17 (0.13, 0.20) |

A major implication of H_5_N_1_ outbreaks among dairy cattle is decreased milk production. Based on our simulations among lactating cows disregarding cow mortality, milk production decreases to 90% of baseline around day 20 before recovering to 91% of baseline. However, when completely discarding milk from clinical cattle as mandated by the Pasteurized Milk Ordinance (28), the usable milk decrease peaks around day 16 at 85% of baseline milk production.

### Web application

The model has been published online as a web application (https://kortessis-lab.shinyapps.io/h5n1/) to allow individual users to explore simulations of H_5_N_1_within dairy herds. The application runs the within-farm model, allowing users to input and modify both herd characteristics and model parameters. The model estimates milk loss in the herd with the option to discard all milk from affected cows as mandated by the Pasteurized Milk Ordinance (28) or to calculate milk loss due to decreased output (7,22). The application is a tool that can be used by dairy farmers to estimate economic impact of an outbreak in terms of both loss of cows and loss of milk production (Figure 4) using local farm characteristics and is flexible enough to allow for constantly updating information about transmission processes and clinical signs that will inevitably occur as new data become available.

**Figure 4.**
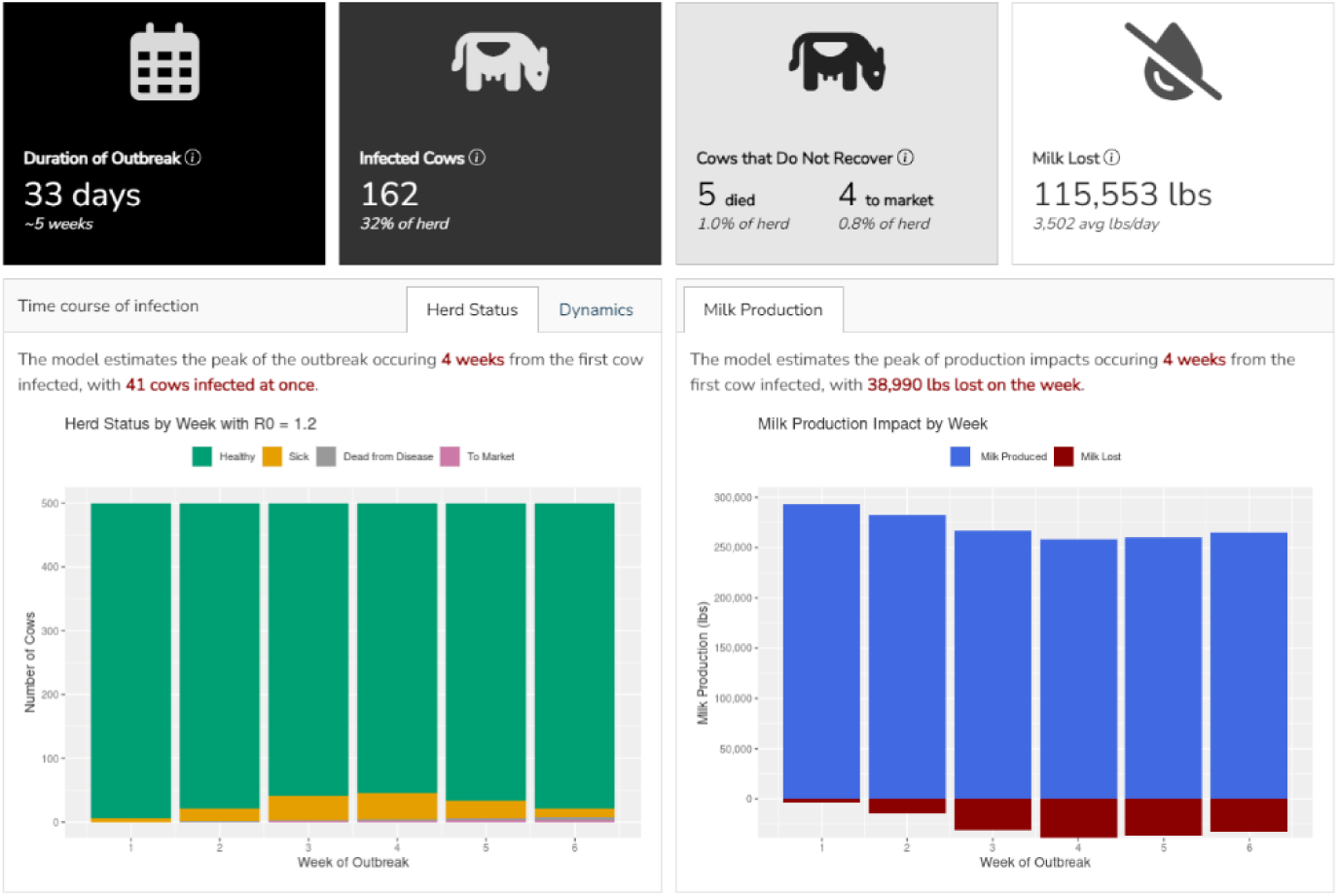
Sample output of H5N1 outbreak model web application. Included are a panel of key performance indicators to highlight important outcomes for a user considering a herd-level outbreak. Additionally, histograms display expected numbers of cows displaying clinical signs and amount of milk production expected throughout the outbreak.

## Discussion

### Model implications

The widespread emergence of H_5_N_1_ in a novel mammalian host demands action. Mathematical models of infectious disease transmission provide a structural framework to synthesize known data into a form conducive to considering control strategies even with limited information from epidemiologic data, laboratory findings, and clinical case reports. By parameterizing the model in a way consistent with scarce data currently available, our major findings are that R_0_ is around 1.2 suggesting that on average, one infectious cow infects around 1.2 other cows over an average of 1.2 days after initial exposure, with a possible lag in overt clinical signs. This fundamental epidemiologic indicator can provide guidance on control strategies. Strategies that could stop spread would need to reduce secondary cases by at least 17%. This could be accomplished either by isolating infectious individuals prior to onward transmission or by reaching a threshold herd immunity. Though seemingly simple, preventing secondary contacts is challenging in the face of the speed at which the virus is spreading. Though milk output of dairy cattle tends to be well monitored, the peak decrease in milk production due to H_5_N_1_ is reported to occur around day 4 post infection (18), well after the time frame we estimate most transmission occurs. Without a sensitive, rapid indicator of infection, isolating infectious cows will be extremely challenging based on existing surveillance systems within dairy operations. Currently no H_5_N_1_ livestock vaccine is available in the United States. Without a method of rapid detection or a vaccine, the virus is continuing to spread among the US dairy industry.

The web-based application accompanying this perspective report can be deployed for individual farms to inform outbreak preparation efforts by giving estimated epidemic size and a rough timeline. This information could be used to order supportive care supplies and hire increased labor to manage sick cows in advance. Users can estimate the economic impact of an outbreak in terms of milk and cow loss under farm specific thresholds for action (e.g., proportion of cows sent to market).

### Current control strategies in context

Continued spread and spillover into other species highlights the importance of control measures. Currently, APHIS requires a negative H_5_N_1_ diagnostic test for interstate movement of lactating cattle (29). The effectiveness of this measure depends on policy compliance, sensitivity of the diagnostic, and transmission mechanism. Other transmission routes, such as transport of other animals or fomites, could allow for continued long-distance viral spread. Though not mandated, APHIS recommends limiting transport of all cattle to help mitigate this risk (29).

Guidance provided by APHIS includes quarantining cows brought onto a farm for 30 days to prevent viral introduction into the resident herd (29). In addition to being impractical for modern-day dairies, quarantine efficacy depends on on-farm biosecurity practices, such as the location of the quarantine facility and how the quarantined animals interact with elements that could enter and leave the quarantine facility such as farm cats, wild birds, staff, and fomites. With H_5_N_1_, our model simulations suggest that there may be cows that are infectious without displaying overt clinical signs. If the quarantine exit criteria is absence of clinical signs as opposed to a negative diagnostic test, there is potential for introduction despite the quarantine via subclinical cases. Heightened monitoring for disease is also recommended (29). Isolation efficacy depends on how quickly transmission occurs in relation to detection. Some surveillance approaches, such as detecting febrile cows or a drop in milk production, may not provide a sensitive enough or timely indicator for containment. Biometric data collected in many dairies has not yet been analyzed for early detection capabilities. Our findings suggest that surveillance must be able to detect outbreaks rapidly with interventions employed quickly to arrest outbreaks.

### Challenges and open questions

Our parameter estimates are highly contingent on early reports providing outbreak overviews and should be taken with caution (7,12,18,22,25). ***Publicly-available well-characterized line-list data from individual outbreaks would greatly enhance modeling certainty.*** Our parameter estimates represent those most consistent with published reports without complete characterization of the outbreaks and may not generalize to other farms. With much yet unknown about transmission of this emerging disease, we made several assumptions in our model. Error included in the model was solely from variation in the stochastic process of disease transmission; true uncertainty is likely much larger. Ideally, we could quantify uncertainty in parameter estimates more robustly, but limited data makes this impossible without strong assumptions. To account for this uncharacterized uncertainty, the web-based application allows users to adjust parameters within a wide range of biologically feasible values; integrate local, farm specific information; and update parameters based on evolving information on H_5_N_1_.

***More research is needed to understand the underlying mechanism of virus spread within a farm.*** We assumed homogenous mixing of cows within a farm. Dairy cattle are generally grouped by lactation stage; the effect of ignoring this spatial contact structure is unknown as the underlying mechanism(s) of transmission is still being investigated. Susceptibility may vary among cattle based on age, lactation stage, level of production, and other factors. Our estimated *R_0_* around 1.2, based on an estimated 30-40% of a herd infected by the end of the outbreak, is relatively low (7). These results suggest a probability of a major outbreak is only 17% and conversely that approximately 83% of introductions end as a single cases and die out. The expansion of H_5_N_1_ from farm to farm suggests a strong force of infection; however, surprisingly, spread within a farm results in a low infection rate. Possible explanations include large proportions of cases being undetected or that an unknown factor—such as the mechanism of viral transmission, bovine contact networks, or heterogeneity in immunity—allows for rapid viral dissemination both within a farm and across the United States, despite low rates of clinical disease. ***Seroprevalence studies can shed light on the number of subclinical or undetected cases*** that ultimately seroconvert without being designated as a case. Our estimated *R_0_* of 1.2 describes the *average* on the farm and results could be much higher in different subpopulations (i.e. lactation stage or cattle age). Heterogeneity among cattle could also explain how outbreaks are being seeded despite a relatively low value of *R_0_*. Understanding the underlying mechanisms of transmission is crucial for a complete understanding of disease spread.

Our model provides a structural perspective to consider disease progression through a population. However*, **additional veterinary clinical research is needed to understand the natural history of disease among cases.*** More studies are needed to understand the relationship between viral shedding, duration of infectiousness, and clinical signs. Our model gives estimates on economic impacts of an outbreak within a farm based on reported levels of decreased milk production; however, long term effects on milk production are still being characterized (18). Another long-term implication of infection is infection-acquired immunity. We simulated recurrent outbreaks based solely on population turnover; the duration of immunity in recovered individuals is unknown. ***The potential and timescale for recurrent outbreaks or endemicity into the cattle population is yet uncharacterized*.**

Though our model provides a framework for considering an outbreak within a dairy farm, ***a more critical framework for public health interventions is a model of between-farm transmission.*** In dairy operations, there are varying levels of movement of cows, equipment, and personnel between farms. Farms may also have other domestic livestock present. This variation in contact structure is recognized in other diseases of veterinary and human importance, such as Foot and Mouth Disease (FMD) (30–32). Major models used in decision-making for control of the 2001 FMD outbreaks in the United Kingdom shared basic elements: the spatial locations of all farms were utilized and distance-informed transmission rates between farms was modeled to simulate disease spread over space and time (30–32). Published agricultural census data gives an overview of dairy cattle operation density in the United States (Figure 1) (16); however, H_5_N_1_ cases are reported at an insufficient resolution (state-level) to replicate FMD models. ***Modeling transmission along this network requires more understanding of the mechanisms of transmission between farms.*** Early genomic studies suggest that the virus likely had one initial spillover event (5,6). Though one study found that interstate movement of cattle led to long-distance spread to several new states, more short-distance spread has been reported in farms without recent cattle introductions (8). Data from outbreak investigations in Michigan report that among 15 affected dairy herds, 9 of them had imported no new cattle, and of the remaining 6, only one outbreak was associated with introduction from imported cattle. The epidemiologic data reported local spread between farms located between 1 and 93 Euclidian miles apart (8). Epidemiologic data from both Michigan and a broader national investigation found many potential mechanisms of local spread including shared staff, equipment, and support service providers (7,8). As the virus transmission continues to expand, publishing data is urgent to inform these models and potential control strategies.

Troubling questions about the emerging H_5_N_1_ infection regard the risk of a spillover event leading to sustained human-to-human transmission with pandemic potential. Regulation from APHIS and the Food and Drug Administration requires milk from clinically affected cows to be discarded and not enter the food supply (28). Spillover events in other animal species have been reported (1,7,8,19), with ingestion of unpasteurized milk from infected cattle being implicated in several cases (1,19). Testing continues to confirm that pasteurized milk remains safe for human consumption (19,20). Currently, all reported cases of the disease in humans have been in people with close dairy cattle contact and presented as mild disease; symptoms have chiefly included mild ocular irritation (11–14). However, H_5_N_1_ has shown historically high case fatality rates in humans with pulmonary involvement (9). Emerging diseases such as H_5_N_1_ in dairy cattle, highlight the need for a comprehensive One Health approach. For some zoonoses, the most effective way to protect human life is to control the disease in animals (33–35). As H_5_N_1_ continues to loom as a pending threat, public health strategies need to consider livestock and poultry level interventions to protect the human population. The results from our model suggest a need to reframe our approach to H_5_N_1_ surveillance and control strategies to prevent continued outbreaks among dairies and onward transmission to humans.

