## Appendix for "Challenges and lessons learned from preliminary modeling of with-in herd transmission of highly pathogenic avian influenza H_5_N_1_ in dairy cattle": AppendixCode1.html

Bellotti et al.

### Background

This code is to generate figures used to support the brief publication accompanying the H5N1 dairy cattle model shiny app. The web application is available at kortessis-lab.shinyapps.io/h5n1/

### Data

#### Load libraries

```
#libraries
library(dplyr)
library(sf)
library(ggplot2)
library(here)
#library(idstyle)
library(tidyUSDA)
library(deSolve)
```

#### Load data

##### **H5N1-affected herd case counts**

- Source: Published by APHIS on their public tableau dashboard
- Notes: Code to scrape tableau dashboard adapted from published github rep: https://github.com/bertrandmartel/tableau-scraping

```
#Function to scrape data from APHIS' public tableau dashboard

scrapeH5N1 <- function(){
  host_url <- "https://publicdashboards.dl.usda.gov"
  path <- "/t/MRP_PUB/views/VS_Cattle_HPAIConfirmedDetections2024/HPAI2022ConfirmedDetections"
  body <- rvest::read_html(httr::modify_url(host_url, 
                               path = path, 
                               query = list(":embed" = "y",":showVizHome" = "no")
  ))
  data <- body %>% 
    rvest::html_nodes("textarea#tsConfigContainer") %>% 
    rvest::html_text()
  json <- rjson::fromJSON(data)
  url <- httr::modify_url(host_url, path = paste(json$vizql_root, "/bootstrapSession/sessions/", json$sessionid, sep =""))
  resp <- httr::POST(url, body = list(sheet_id = json$sheetId), encode = "form")
  data <- httr::content(resp, "text")
  extract <- stringr::str_match(data, "\\d+;(\\{.*\\})\\d+;(\\{.*\\})")
  data <- rjson::fromJSON(extract[1,3])
  worksheets = names(data$secondaryInfo$presModelMap$vizData$presModelHolder$genPresModelMapPresModel$presModelMap)
  selected <-  7
  worksheet <- worksheets[as.integer(selected)]
  columnsData <- data$secondaryInfo$presModelMap$vizData$presModelHolder$genPresModelMapPresModel$presModelMap[[worksheet]]$presModelHolder$genVizDataPresModel$paneColumnsData
  i <- 1
  result <- list();
  for(t in columnsData$vizDataColumns){
    if (is.null(t[["fieldCaption"]]) == FALSE) {
      paneIndex <- t$paneIndices
      columnIndex <- t$columnIndices
      if (length(t$paneIndices) > 1){
        paneIndex <- t$paneIndices[1]
      }
      if (length(t$columnIndices) > 1){
        columnIndex <- t$columnIndices[1]
      }
      result[[i]] <- list(
        fieldCaption = t[["fieldCaption"]], 
        valueIndices = columnsData$paneColumnsList[[paneIndex + 1]]$vizPaneColumns[[columnIndex + 1]]$valueIndices,
        aliasIndices = columnsData$paneColumnsList[[paneIndex + 1]]$vizPaneColumns[[columnIndex + 1]]$aliasIndices, 
        dataType = t[["dataType"]],
        stringsAsFactors = FALSE
      )
      i <- i + 1
    }
  }
  dataFull = data$secondaryInfo$presModelMap$dataDictionary$presModelHolder$genDataDictionaryPresModel$dataSegments[["0"]]$dataColumns
  
  cstring <- list();
  for(t in dataFull) {
    if(t$dataType == "cstring"){
      cstring <- t
      break
    }
  }
  data_index <- 1
  name_index <- 1
  frameData <-  list()
  frameNames <- c()
  for(t in dataFull) {
    for(index in result) {
      if (t$dataType == index["dataType"]){
        if (length(index$valueIndices) > 0) {
          j <- 1
          vector <- character(length(index$valueIndices))
          for (it in index$valueIndices){
            vector[j] <- t$dataValues[it+1]
            j <- j + 1
          }
          frameData[[data_index]] <- vector
          frameNames[[name_index]] <- paste(index$fieldCaption, "value", sep="-")
          data_index <- data_index + 1
          name_index <- name_index + 1
        }
        if (length(index$aliasIndices) > 0) {
          j <- 1
          vector <- character(length(index$aliasIndices))
          for (it in index$aliasIndices){
            if (it >= 0){
              vector[j] <- t$dataValues[it+1]
            } else {
              vector[j] <- cstring$dataValues[abs(it)]
            }
            j <- j + 1
          }
          frameData[[data_index]] <- vector
          frameNames[[name_index]] <- paste(index$fieldCaption, "alias", sep="-")
          data_index <- data_index + 1
          name_index <- name_index + 1
        }
      }
    }
  }
  
  df <- NULL
  lengthList <- c()
  for(i in 1:length(frameNames)){
    lengthList[i] <- length(frameData[[i]])
  }
  max <- max(lengthList)
  for(i in 1:length(frameNames)){
    if (length(frameData[[i]]) < max){
      len <- length(frameData[[i]])
      frameData[[i]][(len+1):max]<-""
    }
    df[frameNames[[i]]] <- frameData[i]
  }
  options(width = 1200)
  df <- as.data.frame(df, stringsAsFactors = FALSE)
  
  #rename
  df <- df%>%
    rename("Operation" = "Production.value",
           "State" = "State.value",
           "Species" = "Species.value",
           "Date" = "Confirmed.value")%>%
    select(Operation, State, Species, Date)
  return(df)
}

#scrape
df.cases<- scrapeH5N1()
```

##### **Shapefiles of US state boundaries**

- Source: US census bureau public repository
- Notes: reading in highest resolution (500k - 1:500,000). Lower resolutions available if desired

```
#function to read zipped shapefiles from web
read_shape_URL <- function(URL){
  cur_tempfile <- tempfile()
  download.file(url = URL, destfile = cur_tempfile)
  out_directory <- tempfile()
  unzip(cur_tempfile, exdir = out_directory)
  
  st_read(dsn = out_directory) #read_sf also works here
}

#read in state shapefiles from census: 
sf.usa <- read_shape_URL("https://www2.census.gov/geo/tiger/GENZ2018/shp/cb_2018_us_state_500k.zip")
sf.usa <- sf.usa%>%
  filter(GEOID < 60 & GEOID != "02" & GEOID != "15") #continental US
usa <-  left_join(sf.usa, df.cases %>% count(State) %>% rename(Cases = n, NAME = State)) #append case counts
```

##### **Dairy cattle density**

- source: **NASS agricultural census**
- note: **Input required -** Update with your API key required to access USDA NASS data

```
##### Need to input your personal USDA API Key here ####
#readRenviron(".env") # Key from USDA 

# Get count of operations with sales in 2017
ops.with.sales <- tidyUSDA::getQuickstat(
  sector = NULL,
  group = NULL,
  commodity = NULL,
  category = NULL,
  domain = NULL,
  county = NULL,
  key = Sys.getenv("KEY"),
  program = "CENSUS",
  data_item = "CATTLE, COWS, MILK - INVENTORY", #dairy operations specific
  geographic_level = "COUNTY",
  year = "2017",
  state = NULL,
  geometry = TRUE,
  lower48 = TRUE)

#clean and format cases
cases <- left_join(df.cases,
                st_drop_geometry(usa) %>% select(STUSPS, NAME) %>% rename('State' = 'NAME'))%>%
  mutate(Date=as.Date(Date))%>%
  group_by(Date) %>%
  mutate(Position = 0.2 + 0.15*(row_number()-1))

cases_summary <- cases%>%
  group_by(STUSPS)%>%
  summarise(Total = n(),
            Initial = min(Date))%>%
  arrange(desc(Total))

#join cases with map
z <- left_join(cases_summary, tigris::states()) |>
  rename("Outbreaks" = "Total") |>
  st_as_sf() |>
  group_by(NAME) |>
  st_centroid()

all_counties <- tigris::counties(state = state.abb[!state.abb %in% c("HI", "AK")])

#join cattle with map
p_cattle_density <- all_counties |>
  select(STATEFP, COUNTYFP, GEOID, ALAND) |>
  mutate(ALAND = ALAND/1e9) |> #unit conversion: square meters to 10k sq km
  left_join(ops.with.sales |>
              as.data.frame() |>
              select(GEOID, Value, domain_desc) |>
              filter(domain_desc == "TOTAL") , by = "GEOID") |>
  mutate(Value = ifelse(is.na(Value), 0, Value))
```

### Model

Code for modified SIR model described in manuscript

```
#input parameter of interest
init.inf = 1
herd.size = 100
R0est = 1.2
fever.duration = 1.15
milk.loss.dur = 7
cow.prod.lifespan = 365.25*2
include.prod.lifespan = FALSE
base.milk.prod = 100 # in units of lbs per day
sympt.milk.prod = 100-11.4
discard.sick.prod = TRUE
disease.induced.mortality = .1
prop.cull = 0.1
sim.length = 365
add_milk = FALSE

state <- c("S" = 99, # Symptomatic compartment
           "I" = 1, # Infectious compartment
           "B" = 0, # Symptomatic, but not infectious compartment
           "R" = 0, # Recovered compartment
           "D" = 0, # Death from disease
           "C" = 0, # Culled compartment
           "Z" = 0) # Dummy compartment to count cows that were infected but died due to productive lifespan

# Set time in units of days
init.time <- 0
end.time <- 365
t.step.dur <- 1 / 24 # time step of an hour
time <- seq(from = init.time, to = end.time, by = t.step.dur)

#Define parameters
parms <- c(beta = R0est / (fever.duration) / herd.size,
           gamma = 1 / fever.duration,
           alpha = 1 / (milk.loss.dur - fever.duration),
           mu = (1 / cow.prod.lifespan) * dplyr::if_else(include.prod.lifespan, 1, 0),
           phi = disease.induced.mortality,
           m = base.milk.prod,
           p = 1 - sympt.milk.prod / base.milk.prod,
           c = prop.cull)


# Write the SIR model in absolute time
SIRMmod <- function (Time, State, pars) {
  with(as.list(c(State, pars)), {
    N <- sum(State) - C - D - Z
    dS = -beta * S * I + N * mu - S * mu
    dI = beta * S * I - gamma * I - I * mu
    dB = (gamma * I) * (1 - phi) - alpha * B - mu * B
    dR =  (1 - c) * alpha * B - mu * R
    dC = alpha * c * B
    dD = gamma * I * phi
    dZ = mu*R + mu*I + mu*B
    return(list(c(dS, dI, dB, dR, dD, dC, dZ)))
  })
}
```

### Figures

##### Figure 1a

```
fig1a <- ggplot() +    
  theme_void(base_size = 24)+   
  geom_sf(data = usa, colour = "grey40", size=0.1, aes(fill = Cases))+
  scale_fill_distiller(direction = 1, palette = "Reds", na.value = "grey60")+
  labs(fill = "Outbreaks \nper state")
```

##### Figure 1b

**Input required:** Update with your API key required to access USDA NASS data

```
fig1b <- ggplot(p_cattle_density) +
  geom_sf(aes(fill = Value/ALAND),  lwd = 0) +
  scale_fill_viridis_c(trans = "log", 
                       labels= c("0", "1", "10", "100", "1k", "10k", "100k"),
                       breaks =c(0, 1, 10, 100, 1000, 10000, 100000)) +
  theme_void(base_size = 24) +
  geom_sf(data = z, aes(size = Outbreaks), 
          shape = 19, alpha=0.5, color = "tomato")+
  scale_size_continuous(range = c(3, 20))+
  guides(fill = guide_legend(position = "top", title.position="top"))+
  labs(fill = "Head of dairy cattle per 1,000 sq. km", size = "Outbreaks \nper state")
```

##### Figure 3

```
out <- ode(state, time, SIRMmod, parms, rtol = 1e-15, maxsteps = 500000)
sub <- as.data.frame(out)%>%
  filter(time < 30)
  
#visualize
Pal1 <- c('Susceptible' = 'dodgerblue3',
          'Infectious' = 'red3',
          'Clinical, not infectious' = 'orange1',
          'Recovered' = 'gold',
          'Died' = 'black',
          'To market'= 'gray50')
    
fig3 <- ggplot()+
  theme_classic(base_size = 36)+ #white background w/ out grid
  geom_line(data=sub, aes(x=time, y=S, color="Susceptible"), size=1.5, alpha=0.8)+
  geom_line(data=sub, aes(x=time, y=I, color="Infectious"), size=1.5, alpha=0.8)+
  geom_line(data=sub, aes(x=time, y=B, color="Clinical, not infectious"), size=1.5, alpha=0.8)+
  geom_line(data=sub, aes(x=time, y=R, color="Recovered"), size=1.5, alpha=0.8)+
  geom_line(data=sub, aes(x=time, y=D, color="Died"), size=1.5, alpha=0.8)+
  geom_line(data=sub, aes(x=time, y=C, color="To market"), size =1.5, alpha=0.8)+
  scale_color_manual(values = Pal1, name= "Disease state",
                     breaks=c("Susceptible","Infectious","Clinical, not infectious", "Recovered", 'Died', 'To market')) +
  # theme(text = element_text(size=20),
  #       axis.text.x = element_text(size=16))+
  labs(y= "% of herd", x = "Time (days)") #axis labels
```

### Other results

##### Milk loss

Code used to generate results for decreased milk production included in the manuscript result texts.

```
#set deathrates to 0 for basic simulation
parms$phi <-0; parms$c <- 0; parms$mu <- 0

#run simulation
out <- ode(state, time, SIRMmod, parms, rtol = 1e-15, maxsteps = 500000)

#calculate milk prodcution
milk <- out %>%
  as.data.frame()%>%
  filter(!(time %% 1))%>%
  mutate(milk_all = base.milk.prod*(S+ .5*I + 0.5*B + 0.75*R),
         milk_discard = base.milk.prod*(S+ 0.75*R),
         baseline = herd.size*base.milk.prod,
         Prop.Milk.All = milk_all/baseline,
         Prop.Milk.Discard = milk_discard/baseline)

#create table
data.frame(
  Milk = c("Keep all produced", "Discard from sick", "both"),
  Scenario = c("Lowest point", "Lowest point", "End of outbreak"),
  Day = c(milk$time[milk$Prop.Milk.All== min(milk$Prop.Milk.All)],
          milk$time[milk$Prop.Milk.Discard== min(milk$Prop.Milk.Discard)],
          45),
  Proportion = c(min(milk$Prop.Milk.All), 
                 min(milk$Prop.Milk.Discard),
                 milk$Prop.Milk.All[milk$time == 45])
  )
```

```
               Milk        Scenario Day Proportion
1 Keep all produced    Lowest point  20  0.9010347
2 Discard from sick    Lowest point  16  0.8503329
3              both End of outbreak  45  0.9122138
```
