## Appendix for "Challenges and lessons learned from preliminary modeling of with-in herd transmission of highly pathogenic avian influenza H_5_N_1_ in dairy cattle": bootstrap.min.js

```
/*!
  * Bootstrap v5.3.1 (https://getbootstrap.com/)
  * Copyright 2011-2023 The Bootstrap Authors (https://github.com/twbs/bootstrap/graphs/contributors)
  * Licensed under MIT (https://github.com/twbs/bootstrap/blob/main/LICENSE)
  */
!function(t,e){"object"==typeof exports&&"undefined"!=typeof module?module.exports=e():"function"==typeof define&&define.amd?define(e):(t="undefined"!=typeof globalThis?globalThis:t||self).bootstrap=e()}(this,(function(){"use strict";const t=new Map,e={set(e,i,n){t.has(e)||t.set(e,new Map);const s=t.get(e);s.has(i)||0===s.size?s.set(i,n):console.error(`Bootstrap doesn't allow more than one instance per element. Bound instance: ${Array.from(s.keys())[0]}.`)},get:(e,i)=>t.has(e)&&t.get(e).get(i)||null,remove(e,i){if(!t.has(e))return;const n=t.get(e);n.delete(i),0===n.size&&t.delete(e)}},i="transitionend",n=t=>(t&&window.CSS&&window.CSS.escape&&(t=t.replace(/#([^\s"#']+)/g,((t,e)=>`#${CSS.escape(e)}`))),t),s=t=>{t.dispatchEvent(new Event(i))},o=t=>!(!t||"object"!=typeof t)&&(void 0!==t.jquery&&(t=t[0]),void 0!==t.nodeType),r=t=>o(t)?t.jquery?t[0]:t:"string"==typeof t&&t.length>0?document.querySelector(n(t)):null,a=t=>{if(!o(t)||0===t.getClientRects().length)return!1;const e="visible"===getComputedStyle(t).getPropertyValue("visibility"),i=t.closest("details:not([open])");if(!i)return e;if(i!==t){const e=t.closest("summary");if(e&&e.parentNode!==i)return!1;if(null===e)return!1}return e},l=t=>!t||t.nodeType!==Node.ELEMENT_NODE||!!t.classList.contains("disabled")||(void 0!==t.disabled?t.disabled:t.hasAttribute("disabled")&&"false"!==t.getAttribute("disabled")),c=t=>{if(!document.documentElement.attachShadow)return null;if("function"==typeof t.getRootNode){const e=t.getRootNode();return e instanceof ShadowRoot?e:null}return t instanceof ShadowRoot?t:t.parentNode?c(t.parentNode):null},h=()=>{},d=t=>{t.offsetHeight},u=()=>window.jQuery&&!document.body.hasAttribute("data-bs-no-jquery")?window.jQuery:null,f=[],p=()=>"rtl"===document.documentElement.dir,m=t=>{var e;e=()=>{const e=u();if(e){const i=t.NAME,n=e.fn[i];e.fn[i]=t.jQueryInterface,e.fn[i].Constructor=t,e.fn[i].noConflict=()=>(e.fn[i]=n,t.jQueryInterface)}},"loading"===document.readyState?(f.length||document.addEventListener("DOMContentLoaded",(()=>{for(const t of f)t()})),f.push(e)):e()},g=(t,e=[],i=t)=>"function"==typeof t?t(...e):i,_=(t,e,n=!0)=>{if(!n)return void g(t);const o=(t=>{if(!t)return 0;let{transitionDuration:e,transitionDelay:i}=window.getComputedStyle(t);const n=Number.parseFloat(e),s=Number.parseFloat(i);return n||s?(e=e.split(",")[0],i=i.split(",")[0],1e3*(Number.parseFloat(e)+Number.parseFloat(i))):0})(e)+5;let r=!1;const a=({target:n})=>{n===e&&(r=!0,e.removeEventListener(i,a),g(t))};e.addEventListener(i,a),setTimeout((()=>{r||s(e)}),o)},b=(t,e,i,n)=>{const s=t.length;let o=t.indexOf(e);return-1===o?!i&&n?t[s-1]:t[0]:(o+=i?1:-1,n&&(o=(o+s)%s),t[Math.max(0,Math.min(o,s-1))])},v=/[^.]*(?=\..*)\.|.*/,y=/\..*/,w=/::\d+$/,A={};let E=1;const T={mouseenter:"mouseover",mouseleave:"mouseout"},C=new Set(["click","dblclick","mouseup","mousedown","contextmenu","mousewheel","DOMMouseScroll","mouseover","mouseout","mousemove","selectstart","selectend","keydown","keypress","keyup","orientationchange","touchstart","touchmove","touchend","touchcancel","pointerdown","pointermove","pointerup","pointerleave","pointercancel","gesturestart","gesturechange","gestureend","focus","blur","change","reset","select","submit","focusin","focusout","load","unload","beforeunload","resize","move","DOMContentLoaded","readystatechange","error","abort","scroll"]);function O(t,e){return e&&`${e}::${E++}`||t.uidEvent||E++}function x(t){const e=O(t);return t.uidEvent=e,A[e]=A[e]||{},A[e]}function k(t,e,i=null){return Object.values(t).find((t=>t.callable===e&&t.delegationSelector===i))}function L(t,e,i){const n="string"==typeof e,s=n?i:e||i;let o=I(t);return C.has(o)||(o=t),[n,s,o]}function S(t,e,i,n,s){if("string"!=typeof e||!t)return;let[o,r,a]=L(e,i,n);if(e in T){const t=t=>function(e){if(!e.relatedTarget||e.relatedTarget!==e.delegateTarget&&!e.delegateTarget.contains(e.relatedTarget))return t.call(this,e)};r=t(r)}const l=x(t),c=l[a]||(l[a]={}),h=k(c,r,o?i:null);if(h)return void(h.oneOff=h.oneOff&&s);const d=O(r,e.replace(v,"")),u=o?function(t,e,i){return function n(s){const o=t.querySelectorAll(e);for(let{target:r}=s;r&&r!==this;r=r.parentNode)for(const a of o)if(a===r)return P(s,{delegateTarget:r}),n.oneOff&&N.off(t,s.type,e,i),i.apply(r,[s])}}(t,i,r):function(t,e){return function i(n){return P(n,{delegateTarget:t}),i.oneOff&&N.off(t,n.type,e),e.apply(t,[n])}}(t,r);u.delegationSelector=o?i:null,u.callable=r,u.oneOff=s,u.uidEvent=d,c[d]=u,t.addEventListener(a,u,o)}function D(t,e,i,n,s){const o=k(e[i],n,s);o&&(t.removeEventListener(i,o,Boolean(s)),delete e[i][o.uidEvent])}function $(t,e,i,n){const s=e[i]||{};for(const[o,r]of Object.entries(s))o.includes(n)&&D(t,e,i,r.callable,r.delegationSelector)}function I(t){return t=t.replace(y,""),T[t]||t}const N={on(t,e,i,n){S(t,e,i,n,!1)},one(t,e,i,n){S(t,e,i,n,!0)},off(t,e,i,n){if("string"!=typeof e||!t)return;const[s,o,r]=L(e,i,n),a=r!==e,l=x(t),c=l[r]||{},h=e.startsWith(".");if(void 0===o){if(h)for(const i of Object.keys(l))$(t,l,i,e.slice(1));for(const[i,n]of Object.entries(c)){const s=i.replace(w,"");a&&!e.includes(s)||D(t,l,r,n.callable,n.delegationSelector)}}else{if(!Object.keys(c).length)return;D(t,l,r,o,s?i:null)}},trigger(t,e,i){if("string"!=typeof e||!t)return null;const n=u();let s=null,o=!0,r=!0,a=!1;e!==I(e)&&n&&(s=n.Event(e,i),n(t).trigger(s),o=!s.isPropagationStopped(),r=!s.isImmediatePropagationStopped(),a=s.isDefaultPrevented());const l=P(new Event(e,{bubbles:o,cancelable:!0}),i);return a&&l.preventDefault(),r&&t.dispatchEvent(l),l.defaultPrevented&&s&&s.preventDefault(),l}};function P(t,e={}){for(const[i,n]of Object.entries(e))try{t[i]=n}catch(e){Object.defineProperty(t,i,{configurable:!0,get:()=>n})}return t}function M(t){if("true"===t)return!0;if("false"===t)return!1;if(t===Number(t).toString())return Number(t);if(""===t||"null"===t)return null;if("string"!=typeof t)return t;try{return JSON.parse(decodeURIComponent(t))}catch(e){return t}}function j(t){return t.replace(/[A-Z]/g,(t=>`-${t.toLowerCase()}`))}const F={setDataAttribute(t,e,i){t.setAttribute(`data-bs-${j(e)}`,i)},removeDataAttribute(t,e){t.removeAttribute(`data-bs-${j(e)}`)},getDataAttributes(t){if(!t)return{};const e={},i=Object.keys(t.dataset).filter((t=>t.startsWith("bs")&&!t.startsWith("bsConfig")));for(const n of i){let i=n.replace(/^bs/,"");i=i.charAt(0).toLowerCase()+i.slice(1,i.length),e[i]=M(t.dataset[n])}return e},getDataAttribute:(t,e)=>M(t.getAttribute(`data-bs-${j(e)}`))};class H{static get Default(){return{}}static get DefaultType(){return{}}static get NAME(){throw new Error('You have to implement the static method "NAME", for each component!')}_getConfig(t){return t=this._mergeConfigObj(t),t=this._configAfterMerge(t),this._typeCheckConfig(t),t}_configAfterMerge(t){return t}_mergeConfigObj(t,e){const i=o(e)?F.getDataAttribute(e,"config"):{};return{...this.constructor.Default,..."object"==typeof i?i:{},...o(e)?F.getDataAttributes(e):{},..."object"==typeof t?t:{}}}_typeCheckConfig(t,e=this.constructor.DefaultType){for(const[n,s]of Object.entries(e)){const e=t[n],r=o(e)?"element":null==(i=e)?`${i}`:Object.prototype.toString.call(i).match(/\s([a-z]+)/i)[1].toLowerCase();if(!new RegExp(s).test(r))throw new TypeError(`${this.constructor.NAME.toUpperCase()}: Option "${n}" provided type "${r}" but expected type "${s}".`)}var i}}class W extends H{constructor(t,i){super(),(t=r(t))&&(this._element=t,this._config=this._getConfig(i),e.set(this._element,this.constructor.DATA_KEY,this))}dispose(){e.remove(this._element,this.constructor.DATA_KEY),N.off(this._element,this.constructor.EVENT_KEY);for(const t of Object.getOwnPropertyNames(this))this[t]=null}_queueCallback(t,e,i=!0){_(t,e,i)}_getConfig(t){return t=this._mergeConfigObj(t,this._element),t=this._configAfterMerge(t),this._typeCheckConfig(t),t}static getInstance(t){return e.get(r(t),this.DATA_KEY)}static getOrCreateInstance(t,e={}){return this.getInstance(t)||new this(t,"object"==typeof e?e:null)}static get VERSION(){return"5.3.1"}static get DATA_KEY(){return`bs.${this.NAME}`}static get EVENT_KEY(){return`.${this.DATA_KEY}`}static eventName(t){return`${t}${this.EVENT_KEY}`}}const B=t=>{let e=t.getAttribute("data-bs-target");if(!e||"#"===e){let i=t.getAttribute("href");if(!i||!i.includes("#")&&!i.startsWith("."))return null;i.includes("#")&&!i.startsWith("#")&&(i=`#${i.split("#")[1]}`),e=i&&"#"!==i?i.trim():null}return n(e)},z={find:(t,e=document.documentElement)=>[].concat(...Element.prototype.querySelectorAll.call(e,t)),findOne:(t,e=document.documentElement)=>Element.prototype.querySelector.call(e,t),children:(t,e)=>[].concat(...t.children).filter((t=>t.matches(e))),parents(t,e){const i=[];let n=t.parentNode.closest(e);for(;n;)i.push(n),n=n.parentNode.closest(e);return i},prev(t,e){let i=t.previousElementSibling;for(;i;){if(i.matches(e))return[i];i=i.previousElementSibling}return[]},next(t,e){let i=t.nextElementSibling;for(;i;){if(i.matches(e))return[i];i=i.nextElementSibling}return[]},focusableChildren(t){const e=["a","button","input","textarea","select","details","[tabindex]",'[contenteditable="true"]'].map((t=>`${t}:not([tabindex^="-"])`)).join(",");return this.find(e,t).filter((t=>!l(t)&&a(t)))},getSelectorFromElement(t){const e=B(t);return e&&z.findOne(e)?e:null},getElementFromSelector(t){const e=B(t);return e?z.findOne(e):null},getMultipleElementsFromSelector(t){const e=B(t);return e?z.find(e):[]}},R=(t,e="hide")=>{const i=`click.dismiss${t.EVENT_KEY}`,n=t.NAME;N.on(document,i,`[data-bs-dismiss="${n}"]`,(function(i){if(["A","AREA"].includes(this.tagName)&&i.preventDefault(),l(this))return;const s=z.getElementFromSelector(this)||this.closest(`.${n}`);t.getOrCreateInstance(s)[e]()}))},q=".bs.alert",V=`close${q}`,K=`closed${q}`;class Q extends W{static get NAME(){return"alert"}close(){if(N.trigger(this._element,V).defaultPrevented)return;this._element.classList.remove("show");const t=this._element.classList.contains("fade");this._queueCallback((()=>this._destroyElement()),this._element,t)}_destroyElement(){this._element.remove(),N.trigger(this._element,K),this.dispose()}static jQueryInterface(t){return this.each((function(){const e=Q.getOrCreateInstance(this);if("string"==typeof t){if(void 0===e[t]||t.startsWith("_")||"constructor"===t)throw new TypeError(`No method named "${t}"`);e[t](this)}}))}}R(Q,"close"),m(Q);const X='[data-bs-toggle="button"]';class Y extends W{static get NAME(){return"button"}toggle(){this._element.setAttribute("aria-pressed",this._element.classList.toggle("active"))}static jQueryInterface(t){return this.each((function(){const e=Y.getOrCreateInstance(this);"toggle"===t&&e[t]()}))}}N.on(document,"click.bs.button.data-api",X,(t=>{t.preventDefault();const e=t.target.closest(X);Y.getOrCreateInstance(e).toggle()})),m(Y);const U=".bs.swipe",G=`touchstart${U}`,J=`touchmove${U}`,Z=`touchend${U}`,tt=`pointerdown${U}`,et=`pointerup${U}`,it={endCallback:null,leftCallback:null,rightCallback:null},nt={endCallback:"(function|null)",leftCallback:"(function|null)",rightCallback:"(function|null)"};class st extends H{constructor(t,e){super(),this._element=t,t&&st.isSupported()&&(this._config=this._getConfig(e),this._deltaX=0,this._supportPointerEvents=Boolean(window.PointerEvent),this._initEvents())}static get Default(){return it}static get DefaultType(){return nt}static get NAME(){return"swipe"}dispose(){N.off(this._element,U)}_start(t){this._supportPointerEvents?this._eventIsPointerPenTouch(t)&&(this._deltaX=t.clientX):this._deltaX=t.touches[0].clientX}_end(t){this._eventIsPointerPenTouch(t)&&(this._deltaX=t.clientX-this._deltaX),this._handleSwipe(),g(this._config.endCallback)}_move(t){this._deltaX=t.touches&&t.touches.length>1?0:t.touches[0].clientX-this._deltaX}_handleSwipe(){const t=Math.abs(this._deltaX);if(t<=40)return;const e=t/this._deltaX;this._deltaX=0,e&&g(e>0?this._config.rightCallback:this._config.leftCallback)}_initEvents(){this._supportPointerEvents?(N.on(this._element,tt,(t=>this._start(t))),N.on(this._element,et,(t=>this._end(t))),this._element.classList.add("pointer-event")):(N.on(this._element,G,(t=>this._start(t))),N.on(this._element,J,(t=>this._move(t))),N.on(this._element,Z,(t=>this._end(t))))}_eventIsPointerPenTouch(t){return this._supportPointerEvents&&("pen"===t.pointerType||"touch"===t.pointerType)}static isSupported(){return"ontouchstart"in document.documentElement||navigator.maxTouchPoints>0}}const ot=".bs.carousel",rt=".data-api",at="next",lt="prev",ct="left",ht="right",dt=`slide${ot}`,ut=`slid${ot}`,ft=`keydown${ot}`,pt=`mouseenter${ot}`,mt=`mouseleave${ot}`,gt=`dragstart${ot}`,_t=`load${ot}${rt}`,bt=`click${ot}${rt}`,vt="carousel",yt="active",wt=".active",At=".carousel-item",Et=wt+At,Tt={ArrowLeft:ht,ArrowRight:ct},Ct={interval:5e3,keyboard:!0,pause:"hover",ride:!1,touch:!0,wrap:!0},Ot={interval:"(number|boolean)",keyboard:"boolean",pause:"(string|boolean)",ride:"(boolean|string)",touch:"boolean",wrap:"boolean"};class xt extends W{constructor(t,e){super(t,e),this._interval=null,this._activeElement=null,this._isSliding=!1,this.touchTimeout=null,this._swipeHelper=null,this._indicatorsElement=z.findOne(".carousel-indicators",this._element),this._addEventListeners(),this._config.ride===vt&&this.cycle()}static get Default(){return Ct}static get DefaultType(){return Ot}static get NAME(){return"carousel"}next(){this._slide(at)}nextWhenVisible(){!document.hidden&&a(this._element)&&this.next()}prev(){this._slide(lt)}pause(){this._isSliding&&s(this._element),this._clearInterval()}cycle(){this._clearInterval(),this._updateInterval(),this._interval=setInterval((()=>this.nextWhenVisible()),this._config.interval)}_maybeEnableCycle(){this._config.ride&&(this._isSliding?N.one(this._element,ut,(()=>this.cycle())):this.cycle())}to(t){const e=this._getItems();if(t>e.length-1||t<0)return;if(this._isSliding)return void N.one(this._element,ut,(()=>this.to(t)));const i=this._getItemIndex(this._getActive());if(i===t)return;const n=t>i?at:lt;this._slide(n,e[t])}dispose(){this._swipeHelper&&this._swipeHelper.dispose(),super.dispose()}_configAfterMerge(t){return t.defaultInterval=t.interval,t}_addEventListeners(){this._config.keyboard&&N.on(this._element,ft,(t=>this._keydown(t))),"hover"===this._config.pause&&(N.on(this._element,pt,(()=>this.pause())),N.on(this._element,mt,(()=>this._maybeEnableCycle()))),this._config.touch&&st.isSupported()&&this._addTouchEventListeners()}_addTouchEventListeners(){for(const t of z.find(".carousel-item img",this._element))N.on(t,gt,(t=>t.preventDefault()));const t={leftCallback:()=>this._slide(this._directionToOrder(ct)),rightCallback:()=>this._slide(this._directionToOrder(ht)),endCallback:()=>{"hover"===this._config.pause&&(this.pause(),this.touchTimeout&&clearTimeout(this.touchTimeout),this.touchTimeout=setTimeout((()=>this._maybeEnableCycle()),500+this._config.interval))}};this._swipeHelper=new st(this._element,t)}_keydown(t){if(/input|textarea/i.test(t.target.tagName))return;const e=Tt[t.key];e&&(t.preventDefault(),this._slide(this._directionToOrder(e)))}_getItemIndex(t){return this._getItems().indexOf(t)}_setActiveIndicatorElement(t){if(!this._indicatorsElement)return;const e=z.findOne(wt,this._indicatorsElement);e.classList.remove(yt),e.removeAttribute("aria-current");const i=z.findOne(`[data-bs-slide-to="${t}"]`,this._indicatorsElement);i&&(i.classList.add(yt),i.setAttribute("aria-current","true"))}_updateInterval(){const t=this._activeElement||this._getActive();if(!t)return;const e=Number.parseInt(t.getAttribute("data-bs-interval"),10);this._config.interval=e||this._config.defaultInterval}_slide(t,e=null){if(this._isSliding)return;const i=this._getActive(),n=t===at,s=e||b(this._getItems(),i,n,this._config.wrap);if(s===i)return;const o=this._getItemIndex(s),r=e=>N.trigger(this._element,e,{relatedTarget:s,direction:this._orderToDirection(t),from:this._getItemIndex(i),to:o});if(r(dt).defaultPrevented)return;if(!i||!s)return;const a=Boolean(this._interval);this.pause(),this._isSliding=!0,this._setActiveIndicatorElement(o),this._activeElement=s;const l=n?"carousel-item-start":"carousel-item-end",c=n?"carousel-item-next":"carousel-item-prev";s.classList.add(c),d(s),i.classList.add(l),s.classList.add(l),this._queueCallback((()=>{s.classList.remove(l,c),s.classList.add(yt),i.classList.remove(yt,c,l),this._isSliding=!1,r(ut)}),i,this._isAnimated()),a&&this.cycle()}_isAnimated(){return this._element.classList.contains("slide")}_getActive(){return z.findOne(Et,this._element)}_getItems(){return z.find(At,this._element)}_clearInterval(){this._interval&&(clearInterval(this._interval),this._interval=null)}_directionToOrder(t){return p()?t===ct?lt:at:t===ct?at:lt}_orderToDirection(t){return p()?t===lt?ct:ht:t===lt?ht:ct}static jQueryInterface(t){return this.each((function(){const e=xt.getOrCreateInstance(this,t);if("number"!=typeof t){if("string"==typeof t){if(void 0===e[t]||t.startsWith("_")||"constructor"===t)throw new TypeError(`No method named "${t}"`);e[t]()}}else e.to(t)}))}}N.on(document,bt,"[data-bs-slide], [data-bs-slide-to]",(function(t){const e=z.getElementFromSelector(this);if(!e||!e.classList.contains(vt))return;t.preventDefault();const i=xt.getOrCreateInstance(e),n=this.getAttribute("data-bs-slide-to");return n?(i.to(n),void i._maybeEnableCycle()):"next"===F.getDataAttribute(this,"slide")?(i.next(),void i._maybeEnableCycle()):(i.prev(),void i._maybeEnableCycle())})),N.on(window,_t,(()=>{const t=z.find('[data-bs-ride="carousel"]');for(const e of t)xt.getOrCreateInstance(e)})),m(xt);const kt=".bs.collapse",Lt=`show${kt}`,St=`shown${kt}`,Dt=`hide${kt}`,$t=`hidden${kt}`,It=`click${kt}.data-api`,Nt="show",Pt="collapse",Mt="collapsing",jt=`:scope .${Pt} .${Pt}`,Ft='[data-bs-toggle="collapse"]',Ht={parent:null,toggle:!0},Wt={parent:"(null|element)",toggle:"boolean"};class Bt extends W{constructor(t,e){super(t,e),this._isTransitioning=!1,this._triggerArray=[];const i=z.find(Ft);for(const t of i){const e=z.getSelectorFromElement(t),i=z.find(e).filter((t=>t===this._element));null!==e&&i.length&&this._triggerArray.push(t)}this._initializeChildren(),this._config.parent||this._addAriaAndCollapsedClass(this._triggerArray,this._isShown()),this._config.toggle&&this.toggle()}static get Default(){return Ht}static get DefaultType(){return Wt}static get NAME(){return"collapse"}toggle(){this._isShown()?this.hide():this.show()}show(){if(this._isTransitioning||this._isShown())return;let t=[];if(this._config.parent&&(t=this._getFirstLevelChildren(".collapse.show, .collapse.collapsing").filter((t=>t!==this._element)).map((t=>Bt.getOrCreateInstance(t,{toggle:!1})))),t.length&&t[0]._isTransitioning)return;if(N.trigger(this._element,Lt).defaultPrevented)return;for(const e of t)e.hide();const e=this._getDimension();this._element.classList.remove(Pt),this._element.classList.add(Mt),this._element.style[e]=0,this._addAriaAndCollapsedClass(this._triggerArray,!0),this._isTransitioning=!0;const i=`scroll${e[0].toUpperCase()+e.slice(1)}`;this._queueCallback((()=>{this._isTransitioning=!1,this._element.classList.remove(Mt),this._element.classList.add(Pt,Nt),this._element.style[e]="",N.trigger(this._element,St)}),this._element,!0),this._element.style[e]=`${this._element[i]}px`}hide(){if(this._isTransitioning||!this._isShown())return;if(N.trigger(this._element,Dt).defaultPrevented)return;const t=this._getDimension();this._element.style[t]=`${this._element.getBoundingClientRect()[t]}px`,d(this._element),this._element.classList.add(Mt),this._element.classList.remove(Pt,Nt);for(const t of this._triggerArray){const e=z.getElementFromSelector(t);e&&!this._isShown(e)&&this._addAriaAndCollapsedClass([t],!1)}this._isTransitioning=!0,this._element.style[t]="",this._queueCallback((()=>{this._isTransitioning=!1,this._element.classList.remove(Mt),this._element.classList.add(Pt),N.trigger(this._element,$t)}),this._element,!0)}_isShown(t=this._element){return t.classList.contains(Nt)}_configAfterMerge(t){return t.toggle=Boolean(t.toggle),t.parent=r(t.parent),t}_getDimension(){return this._element.classList.contains("collapse-horizontal")?"width":"height"}_initializeChildren(){if(!this._config.parent)return;const t=this._getFirstLevelChildren(Ft);for(const e of t){const t=z.getElementFromSelector(e);t&&this._addAriaAndCollapsedClass([e],this._isShown(t))}}_getFirstLevelChildren(t){const e=z.find(jt,this._config.parent);return z.find(t,this._config.parent).filter((t=>!e.includes(t)))}_addAriaAndCollapsedClass(t,e){if(t.length)for(const i of t)i.classList.toggle("collapsed",!e),i.setAttribute("aria-expanded",e)}static jQueryInterface(t){const e={};return"string"==typeof t&&/show|hide/.test(t)&&(e.toggle=!1),this.each((function(){const i=Bt.getOrCreateInstance(this,e);if("string"==typeof t){if(void 0===i[t])throw new TypeError(`No method named "${t}"`);i[t]()}}))}}N.on(document,It,Ft,(function(t){("A"===t.target.tagName||t.delegateTarget&&"A"===t.delegateTarget.tagName)&&t.preventDefault();for(const t of z.getMultipleElementsFromSelector(this))Bt.getOrCreateInstance(t,{toggle:!1}).toggle()})),m(Bt);var zt="top",Rt="bottom",qt="right",Vt="left",Kt="auto",Qt=[zt,Rt,qt,Vt],Xt="start",Yt="end",Ut="clippingParents",Gt="viewport",Jt="popper",Zt="reference",te=Qt.reduce((function(t,e){return t.concat([e+"-"+Xt,e+"-"+Yt])}),[]),ee=[].concat(Qt,[Kt]).reduce((function(t,e){return t.concat([e,e+"-"+Xt,e+"-"+Yt])}),[]),ie="beforeRead",ne="read",se="afterRead",oe="beforeMain",re="main",ae="afterMain",le="beforeWrite",ce="write",he="afterWrite",de=[ie,ne,se,oe,re,ae,le,ce,he];function ue(t){return t?(t.nodeName||"").toLowerCase():null}function fe(t){if(null==t)return window;if("[object Window]"!==t.toString()){var e=t.ownerDocument;return e&&e.defaultView||window}return t}function pe(t){return t instanceof fe(t).Element||t instanceof Element}function me(t){return t instanceof fe(t).HTMLElement||t instanceof HTMLElement}function ge(t){return"undefined"!=typeof ShadowRoot&&(t instanceof fe(t).ShadowRoot||t instanceof ShadowRoot)}const _e={name:"applyStyles",enabled:!0,phase:"write",fn:function(t){var e=t.state;Object.keys(e.elements).forEach((function(t){var i=e.styles[t]||{},n=e.attributes[t]||{},s=e.elements[t];me(s)&&ue(s)&&(Object.assign(s.style,i),Object.keys(n).forEach((function(t){var e=n[t];!1===e?s.removeAttribute(t):s.setAttribute(t,!0===e?"":e)})))}))},effect:function(t){var e=t.state,i={popper:{position:e.options.strategy,left:"0",top:"0",margin:"0"},arrow:{position:"absolute"},reference:{}};return Object.assign(e.elements.popper.style,i.popper),e.styles=i,e.elements.arrow&&Object.assign(e.elements.arrow.style,i.arrow),function(){Object.keys(e.elements).forEach((function(t){var n=e.elements[t],s=e.attributes[t]||{},o=Object.keys(e.styles.hasOwnProperty(t)?e.styles[t]:i[t]).reduce((function(t,e){return t[e]="",t}),{});me(n)&&ue(n)&&(Object.assign(n.style,o),Object.keys(s).forEach((function(t){n.removeAttribute(t)})))}))}},requires:["computeStyles"]};function be(t){return t.split("-")[0]}var ve=Math.max,ye=Math.min,we=Math.round;function Ae(){var t=navigator.userAgentData;return null!=t&&t.brands&&Array.isArray(t.brands)?t.brands.map((function(t){return t.brand+"/"+t.version})).join(" "):navigator.userAgent}function Ee(){return!/^((?!chrome|android).)*safari/i.test(Ae())}function Te(t,e,i){void 0===e&&(e=!1),void 0===i&&(i=!1);var n=t.getBoundingClientRect(),s=1,o=1;e&&me(t)&&(s=t.offsetWidth>0&&we(n.width)/t.offsetWidth||1,o=t.offsetHeight>0&&we(n.height)/t.offsetHeight||1);var r=(pe(t)?fe(t):window).visualViewport,a=!Ee()&&i,l=(n.left+(a&&r?r.offsetLeft:0))/s,c=(n.top+(a&&r?r.offsetTop:0))/o,h=n.width/s,d=n.height/o;return{width:h,height:d,top:c,right:l+h,bottom:c+d,left:l,x:l,y:c}}function Ce(t){var e=Te(t),i=t.offsetWidth,n=t.offsetHeight;return Math.abs(e.width-i)<=1&&(i=e.width),Math.abs(e.height-n)<=1&&(n=e.height),{x:t.offsetLeft,y:t.offsetTop,width:i,height:n}}function Oe(t,e){var i=e.getRootNode&&e.getRootNode();if(t.contains(e))return!0;if(i&&ge(i)){var n=e;do{if(n&&t.isSameNode(n))return!0;n=n.parentNode||n.host}while(n)}return!1}function xe(t){return fe(t).getComputedStyle(t)}function ke(t){return["table","td","th"].indexOf(ue(t))>=0}function Le(t){return((pe(t)?t.ownerDocument:t.document)||window.document).documentElement}function Se(t){return"html"===ue(t)?t:t.assignedSlot||t.parentNode||(ge(t)?t.host:null)||Le(t)}function De(t){return me(t)&&"fixed"!==xe(t).position?t.offsetParent:null}function $e(t){for(var e=fe(t),i=De(t);i&&ke(i)&&"static"===xe(i).position;)i=De(i);return i&&("html"===ue(i)||"body"===ue(i)&&"static"===xe(i).position)?e:i||function(t){var e=/firefox/i.test(Ae());if(/Trident/i.test(Ae())&&me(t)&&"fixed"===xe(t).position)return null;var i=Se(t);for(ge(i)&&(i=i.host);me(i)&&["html","body"].indexOf(ue(i))<0;){var n=xe(i);if("none"!==n.transform||"none"!==n.perspective||"paint"===n.contain||-1!==["transform","perspective"].indexOf(n.willChange)||e&&"filter"===n.willChange||e&&n.filter&&"none"!==n.filter)return i;i=i.parentNode}return null}(t)||e}function Ie(t){return["top","bottom"].indexOf(t)>=0?"x":"y"}function Ne(t,e,i){return ve(t,ye(e,i))}function Pe(t){return Object.assign({},{top:0,right:0,bottom:0,left:0},t)}function Me(t,e){return e.reduce((function(e,i){return e[i]=t,e}),{})}const je={name:"arrow",enabled:!0,phase:"main",fn:function(t){var e,i=t.state,n=t.name,s=t.options,o=i.elements.arrow,r=i.modifiersData.popperOffsets,a=be(i.placement),l=Ie(a),c=[Vt,qt].indexOf(a)>=0?"height":"width";if(o&&r){var h=function(t,e){return Pe("number"!=typeof(t="function"==typeof t?t(Object.assign({},e.rects,{placement:e.placement})):t)?t:Me(t,Qt))}(s.padding,i),d=Ce(o),u="y"===l?zt:Vt,f="y"===l?Rt:qt,p=i.rects.reference[c]+i.rects.reference[l]-r[l]-i.rects.popper[c],m=r[l]-i.rects.reference[l],g=$e(o),_=g?"y"===l?g.clientHeight||0:g.clientWidth||0:0,b=p/2-m/2,v=h[u],y=_-d[c]-h[f],w=_/2-d[c]/2+b,A=Ne(v,w,y),E=l;i.modifiersData[n]=((e={})[E]=A,e.centerOffset=A-w,e)}},effect:function(t){var e=t.state,i=t.options.element,n=void 0===i?"[data-popper-arrow]":i;null!=n&&("string"!=typeof n||(n=e.elements.popper.querySelector(n)))&&Oe(e.elements.popper,n)&&(e.elements.arrow=n)},requires:["popperOffsets"],requiresIfExists:["preventOverflow"]};function Fe(t){return t.split("-")[1]}var He={top:"auto",right:"auto",bottom:"auto",left:"auto"};function We(t){var e,i=t.popper,n=t.popperRect,s=t.placement,o=t.variation,r=t.offsets,a=t.position,l=t.gpuAcceleration,c=t.adaptive,h=t.roundOffsets,d=t.isFixed,u=r.x,f=void 0===u?0:u,p=r.y,m=void 0===p?0:p,g="function"==typeof h?h({x:f,y:m}):{x:f,y:m};f=g.x,m=g.y;var _=r.hasOwnProperty("x"),b=r.hasOwnProperty("y"),v=Vt,y=zt,w=window;if(c){var A=$e(i),E="clientHeight",T="clientWidth";A===fe(i)&&"static"!==xe(A=Le(i)).position&&"absolute"===a&&(E="scrollHeight",T="scrollWidth"),(s===zt||(s===Vt||s===qt)&&o===Yt)&&(y=Rt,m-=(d&&A===w&&w.visualViewport?w.visualViewport.height:A[E])-n.height,m*=l?1:-1),s!==Vt&&(s!==zt&&s!==Rt||o!==Yt)||(v=qt,f-=(d&&A===w&&w.visualViewport?w.visualViewport.width:A[T])-n.width,f*=l?1:-1)}var C,O=Object.assign({position:a},c&&He),x=!0===h?function(t,e){var i=t.x,n=t.y,s=e.devicePixelRatio||1;return{x:we(i*s)/s||0,y:we(n*s)/s||0}}({x:f,y:m},fe(i)):{x:f,y:m};return f=x.x,m=x.y,l?Object.assign({},O,((C={})[y]=b?"0":"",C[v]=_?"0":"",C.transform=(w.devicePixelRatio||1)<=1?"translate("+f+"px, "+m+"px)":"translate3d("+f+"px, "+m+"px, 0)",C)):Object.assign({},O,((e={})[y]=b?m+"px":"",e[v]=_?f+"px":"",e.transform="",e))}const Be={name:"computeStyles",enabled:!0,phase:"beforeWrite",fn:function(t){var e=t.state,i=t.options,n=i.gpuAcceleration,s=void 0===n||n,o=i.adaptive,r=void 0===o||o,a=i.roundOffsets,l=void 0===a||a,c={placement:be(e.placement),variation:Fe(e.placement),popper:e.elements.popper,popperRect:e.rects.popper,gpuAcceleration:s,isFixed:"fixed"===e.options.strategy};null!=e.modifiersData.popperOffsets&&(e.styles.popper=Object.assign({},e.styles.popper,We(Object.assign({},c,{offsets:e.modifiersData.popperOffsets,position:e.options.strategy,adaptive:r,roundOffsets:l})))),null!=e.modifiersData.arrow&&(e.styles.arrow=Object.assign({},e.styles.arrow,We(Object.assign({},c,{offsets:e.modifiersData.arrow,position:"absolute",adaptive:!1,roundOffsets:l})))),e.attributes.popper=Object.assign({},e.attributes.popper,{"data-popper-placement":e.placement})},data:{}};var ze={passive:!0};const Re={name:"eventListeners",enabled:!0,phase:"write",fn:function(){},effect:function(t){var e=t.state,i=t.instance,n=t.options,s=n.scroll,o=void 0===s||s,r=n.resize,a=void 0===r||r,l=fe(e.elements.popper),c=[].concat(e.scrollParents.reference,e.scrollParents.popper);return o&&c.forEach((function(t){t.addEventListener("scroll",i.update,ze)})),a&&l.addEventListener("resize",i.update,ze),function(){o&&c.forEach((function(t){t.removeEventListener("scroll",i.update,ze)})),a&&l.removeEventListener("resize",i.update,ze)}},data:{}};var qe={left:"right",right:"left",bottom:"top",top:"bottom"};function Ve(t){return t.replace(/left|right|bottom|top/g,(function(t){return qe[t]}))}var Ke={start:"end",end:"start"};function Qe(t){return t.replace(/start|end/g,(function(t){return Ke[t]}))}function Xe(t){var e=fe(t);return{scrollLeft:e.pageXOffset,scrollTop:e.pageYOffset}}function Ye(t){return Te(Le(t)).left+Xe(t).scrollLeft}function Ue(t){var e=xe(t),i=e.overflow,n=e.overflowX,s=e.overflowY;return/auto|scroll|overlay|hidden/.test(i+s+n)}function Ge(t){return["html","body","#document"].indexOf(ue(t))>=0?t.ownerDocument.body:me(t)&&Ue(t)?t:Ge(Se(t))}function Je(t,e){var i;void 0===e&&(e=[]);var n=Ge(t),s=n===(null==(i=t.ownerDocument)?void 0:i.body),o=fe(n),r=s?[o].concat(o.visualViewport||[],Ue(n)?n:[]):n,a=e.concat(r);return s?a:a.concat(Je(Se(r)))}function Ze(t){return Object.assign({},t,{left:t.x,top:t.y,right:t.x+t.width,bottom:t.y+t.height})}function ti(t,e,i){return e===Gt?Ze(function(t,e){var i=fe(t),n=Le(t),s=i.visualViewport,o=n.clientWidth,r=n.clientHeight,a=0,l=0;if(s){o=s.width,r=s.height;var c=Ee();(c||!c&&"fixed"===e)&&(a=s.offsetLeft,l=s.offsetTop)}return{width:o,height:r,x:a+Ye(t),y:l}}(t,i)):pe(e)?function(t,e){var i=Te(t,!1,"fixed"===e);return i.top=i.top+t.clientTop,i.left=i.left+t.clientLeft,i.bottom=i.top+t.clientHeight,i.right=i.left+t.clientWidth,i.width=t.clientWidth,i.height=t.clientHeight,i.x=i.left,i.y=i.top,i}(e,i):Ze(function(t){var e,i=Le(t),n=Xe(t),s=null==(e=t.ownerDocument)?void 0:e.body,o=ve(i.scrollWidth,i.clientWidth,s?s.scrollWidth:0,s?s.clientWidth:0),r=ve(i.scrollHeight,i.clientHeight,s?s.scrollHeight:0,s?s.clientHeight:0),a=-n.scrollLeft+Ye(t),l=-n.scrollTop;return"rtl"===xe(s||i).direction&&(a+=ve(i.clientWidth,s?s.clientWidth:0)-o),{width:o,height:r,x:a,y:l}}(Le(t)))}function ei(t){var e,i=t.reference,n=t.element,s=t.placement,o=s?be(s):null,r=s?Fe(s):null,a=i.x+i.width/2-n.width/2,l=i.y+i.height/2-n.height/2;switch(o){case zt:e={x:a,y:i.y-n.height};break;case Rt:e={x:a,y:i.y+i.height};break;case qt:e={x:i.x+i.width,y:l};break;case Vt:e={x:i.x-n.width,y:l};break;default:e={x:i.x,y:i.y}}var c=o?Ie(o):null;if(null!=c){var h="y"===c?"height":"width";switch(r){case Xt:e[c]=e[c]-(i[h]/2-n[h]/2);break;case Yt:e[c]=e[c]+(i[h]/2-n[h]/2)}}return e}function ii(t,e){void 0===e&&(e={});var i=e,n=i.placement,s=void 0===n?t.placement:n,o=i.strategy,r=void 0===o?t.strategy:o,a=i.boundary,l=void 0===a?Ut:a,c=i.rootBoundary,h=void 0===c?Gt:c,d=i.elementContext,u=void 0===d?Jt:d,f=i.altBoundary,p=void 0!==f&&f,m=i.padding,g=void 0===m?0:m,_=Pe("number"!=typeof g?g:Me(g,Qt)),b=u===Jt?Zt:Jt,v=t.rects.popper,y=t.elements[p?b:u],w=function(t,e,i,n){var s="clippingParents"===e?function(t){var e=Je(Se(t)),i=["absolute","fixed"].indexOf(xe(t).position)>=0&&me(t)?$e(t):t;return pe(i)?e.filter((function(t){return pe(t)&&Oe(t,i)&&"body"!==ue(t)})):[]}(t):[].concat(e),o=[].concat(s,[i]),r=o[0],a=o.reduce((function(e,i){var s=ti(t,i,n);return e.top=ve(s.top,e.top),e.right=ye(s.right,e.right),e.bottom=ye(s.bottom,e.bottom),e.left=ve(s.left,e.left),e}),ti(t,r,n));return a.width=a.right-a.left,a.height=a.bottom-a.top,a.x=a.left,a.y=a.top,a}(pe(y)?y:y.contextElement||Le(t.elements.popper),l,h,r),A=Te(t.elements.reference),E=ei({reference:A,element:v,strategy:"absolute",placement:s}),T=Ze(Object.assign({},v,E)),C=u===Jt?T:A,O={top:w.top-C.top+_.top,bottom:C.bottom-w.bottom+_.bottom,left:w.left-C.left+_.left,right:C.right-w.right+_.right},x=t.modifiersData.offset;if(u===Jt&&x){var k=x[s];Object.keys(O).forEach((function(t){var e=[qt,Rt].indexOf(t)>=0?1:-1,i=[zt,Rt].indexOf(t)>=0?"y":"x";O[t]+=k[i]*e}))}return O}function ni(t,e){void 0===e&&(e={});var i=e,n=i.placement,s=i.boundary,o=i.rootBoundary,r=i.padding,a=i.flipVariations,l=i.allowedAutoPlacements,c=void 0===l?ee:l,h=Fe(n),d=h?a?te:te.filter((function(t){return Fe(t)===h})):Qt,u=d.filter((function(t){return c.indexOf(t)>=0}));0===u.length&&(u=d);var f=u.reduce((function(e,i){return e[i]=ii(t,{placement:i,boundary:s,rootBoundary:o,padding:r})[be(i)],e}),{});return Object.keys(f).sort((function(t,e){return f[t]-f[e]}))}const si={name:"flip",enabled:!0,phase:"main",fn:function(t){var e=t.state,i=t.options,n=t.name;if(!e.modifiersData[n]._skip){for(var s=i.mainAxis,o=void 0===s||s,r=i.altAxis,a=void 0===r||r,l=i.fallbackPlacements,c=i.padding,h=i.boundary,d=i.rootBoundary,u=i.altBoundary,f=i.flipVariations,p=void 0===f||f,m=i.allowedAutoPlacements,g=e.options.placement,_=be(g),b=l||(_!==g&&p?function(t){if(be(t)===Kt)return[];var e=Ve(t);return[Qe(t),e,Qe(e)]}(g):[Ve(g)]),v=[g].concat(b).reduce((function(t,i){return t.concat(be(i)===Kt?ni(e,{placement:i,boundary:h,rootBoundary:d,padding:c,flipVariations:p,allowedAutoPlacements:m}):i)}),[]),y=e.rects.reference,w=e.rects.popper,A=new Map,E=!0,T=v[0],C=0;C<v.length;C++){var O=v[C],x=be(O),k=Fe(O)===Xt,L=[zt,Rt].indexOf(x)>=0,S=L?"width":"height",D=ii(e,{placement:O,boundary:h,rootBoundary:d,altBoundary:u,padding:c}),$=L?k?qt:Vt:k?Rt:zt;y[S]>w[S]&&($=Ve($));var I=Ve($),N=[];if(o&&N.push(D[x]<=0),a&&N.push(D[$]<=0,D[I]<=0),N.every((function(t){return t}))){T=O,E=!1;break}A.set(O,N)}if(E)for(var P=function(t){var e=v.find((function(e){var i=A.get(e);if(i)return i.slice(0,t).every((function(t){return t}))}));if(e)return T=e,"break"},M=p?3:1;M>0&&"break"!==P(M);M--);e.placement!==T&&(e.modifiersData[n]._skip=!0,e.placement=T,e.reset=!0)}},requiresIfExists:["offset"],data:{_skip:!1}};function oi(t,e,i){return void 0===i&&(i={x:0,y:0}),{top:t.top-e.height-i.y,right:t.right-e.width+i.x,bottom:t.bottom-e.height+i.y,left:t.left-e.width-i.x}}function ri(t){return[zt,qt,Rt,Vt].some((function(e){return t[e]>=0}))}const ai={name:"hide",enabled:!0,phase:"main",requiresIfExists:["preventOverflow"],fn:function(t){var e=t.state,i=t.name,n=e.rects.reference,s=e.rects.popper,o=e.modifiersData.preventOverflow,r=ii(e,{elementContext:"reference"}),a=ii(e,{altBoundary:!0}),l=oi(r,n),c=oi(a,s,o),h=ri(l),d=ri(c);e.modifiersData[i]={referenceClippingOffsets:l,popperEscapeOffsets:c,isReferenceHidden:h,hasPopperEscaped:d},e.attributes.popper=Object.assign({},e.attributes.popper,{"data-popper-reference-hidden":h,"data-popper-escaped":d})}},li={name:"offset",enabled:!0,phase:"main",requires:["popperOffsets"],fn:function(t){var e=t.state,i=t.options,n=t.name,s=i.offset,o=void 0===s?[0,0]:s,r=ee.reduce((function(t,i){return t[i]=function(t,e,i){var n=be(t),s=[Vt,zt].indexOf(n)>=0?-1:1,o="function"==typeof i?i(Object.assign({},e,{placement:t})):i,r=o[0],a=o[1];return r=r||0,a=(a||0)*s,[Vt,qt].indexOf(n)>=0?{x:a,y:r}:{x:r,y:a}}(i,e.rects,o),t}),{}),a=r[e.placement],l=a.x,c=a.y;null!=e.modifiersData.popperOffsets&&(e.modifiersData.popperOffsets.x+=l,e.modifiersData.popperOffsets.y+=c),e.modifiersData[n]=r}},ci={name:"popperOffsets",enabled:!0,phase:"read",fn:function(t){var e=t.state,i=t.name;e.modifiersData[i]=ei({reference:e.rects.reference,element:e.rects.popper,strategy:"absolute",placement:e.placement})},data:{}},hi={name:"preventOverflow",enabled:!0,phase:"main",fn:function(t){var e=t.state,i=t.options,n=t.name,s=i.mainAxis,o=void 0===s||s,r=i.altAxis,a=void 0!==r&&r,l=i.boundary,c=i.rootBoundary,h=i.altBoundary,d=i.padding,u=i.tether,f=void 0===u||u,p=i.tetherOffset,m=void 0===p?0:p,g=ii(e,{boundary:l,rootBoundary:c,padding:d,altBoundary:h}),_=be(e.placement),b=Fe(e.placement),v=!b,y=Ie(_),w="x"===y?"y":"x",A=e.modifiersData.popperOffsets,E=e.rects.reference,T=e.rects.popper,C="function"==typeof m?m(Object.assign({},e.rects,{placement:e.placement})):m,O="number"==typeof C?{mainAxis:C,altAxis:C}:Object.assign({mainAxis:0,altAxis:0},C),x=e.modifiersData.offset?e.modifiersData.offset[e.placement]:null,k={x:0,y:0};if(A){if(o){var L,S="y"===y?zt:Vt,D="y"===y?Rt:qt,$="y"===y?"height":"width",I=A[y],N=I+g[S],P=I-g[D],M=f?-T[$]/2:0,j=b===Xt?E[$]:T[$],F=b===Xt?-T[$]:-E[$],H=e.elements.arrow,W=f&&H?Ce(H):{width:0,height:0},B=e.modifiersData["arrow#persistent"]?e.modifiersData["arrow#persistent"].padding:{top:0,right:0,bottom:0,left:0},z=B[S],R=B[D],q=Ne(0,E[$],W[$]),V=v?E[$]/2-M-q-z-O.mainAxis:j-q-z-O.mainAxis,K=v?-E[$]/2+M+q+R+O.mainAxis:F+q+R+O.mainAxis,Q=e.elements.arrow&&$e(e.elements.arrow),X=Q?"y"===y?Q.clientTop||0:Q.clientLeft||0:0,Y=null!=(L=null==x?void 0:x[y])?L:0,U=I+K-Y,G=Ne(f?ye(N,I+V-Y-X):N,I,f?ve(P,U):P);A[y]=G,k[y]=G-I}if(a){var J,Z="x"===y?zt:Vt,tt="x"===y?Rt:qt,et=A[w],it="y"===w?"height":"width",nt=et+g[Z],st=et-g[tt],ot=-1!==[zt,Vt].indexOf(_),rt=null!=(J=null==x?void 0:x[w])?J:0,at=ot?nt:et-E[it]-T[it]-rt+O.altAxis,lt=ot?et+E[it]+T[it]-rt-O.altAxis:st,ct=f&&ot?function(t,e,i){var n=Ne(t,e,i);return n>i?i:n}(at,et,lt):Ne(f?at:nt,et,f?lt:st);A[w]=ct,k[w]=ct-et}e.modifiersData[n]=k}},requiresIfExists:["offset"]};function di(t,e,i){void 0===i&&(i=!1);var n,s,o=me(e),r=me(e)&&function(t){var e=t.getBoundingClientRect(),i=we(e.width)/t.offsetWidth||1,n=we(e.height)/t.offsetHeight||1;return 1!==i||1!==n}(e),a=Le(e),l=Te(t,r,i),c={scrollLeft:0,scrollTop:0},h={x:0,y:0};return(o||!o&&!i)&&(("body"!==ue(e)||Ue(a))&&(c=(n=e)!==fe(n)&&me(n)?{scrollLeft:(s=n).scrollLeft,scrollTop:s.scrollTop}:Xe(n)),me(e)?((h=Te(e,!0)).x+=e.clientLeft,h.y+=e.clientTop):a&&(h.x=Ye(a))),{x:l.left+c.scrollLeft-h.x,y:l.top+c.scrollTop-h.y,width:l.width,height:l.height}}function ui(t){var e=new Map,i=new Set,n=[];function s(t){i.add(t.name),[].concat(t.requires||[],t.requiresIfExists||[]).forEach((function(t){if(!i.has(t)){var n=e.get(t);n&&s(n)}})),n.push(t)}return t.forEach((function(t){e.set(t.name,t)})),t.forEach((function(t){i.has(t.name)||s(t)})),n}var fi={placement:"bottom",modifiers:[],strategy:"absolute"};function pi(){for(var t=arguments.length,e=new Array(t),i=0;i<t;i++)e[i]=arguments[i];return!e.some((function(t){return!(t&&"function"==typeof t.getBoundingClientRect)}))}function mi(t){void 0===t&&(t={});var e=t,i=e.defaultModifiers,n=void 0===i?[]:i,s=e.defaultOptions,o=void 0===s?fi:s;return function(t,e,i){void 0===i&&(i=o);var s,r,a={placement:"bottom",orderedModifiers:[],options:Object.assign({},fi,o),modifiersData:{},elements:{reference:t,popper:e},attributes:{},styles:{}},l=[],c=!1,h={state:a,setOptions:function(i){var s="function"==typeof i?i(a.options):i;d(),a.options=Object.assign({},o,a.options,s),a.scrollParents={reference:pe(t)?Je(t):t.contextElement?Je(t.contextElement):[],popper:Je(e)};var r,c,u=function(t){var e=ui(t);return de.reduce((function(t,i){return t.concat(e.filter((function(t){return t.phase===i})))}),[])}((r=[].concat(n,a.options.modifiers),c=r.reduce((function(t,e){var i=t[e.name];return t[e.name]=i?Object.assign({},i,e,{options:Object.assign({},i.options,e.options),data:Object.assign({},i.data,e.data)}):e,t}),{}),Object.keys(c).map((function(t){return c[t]}))));return a.orderedModifiers=u.filter((function(t){return t.enabled})),a.orderedModifiers.forEach((function(t){var e=t.name,i=t.options,n=void 0===i?{}:i,s=t.effect;if("function"==typeof s){var o=s({state:a,name:e,instance:h,options:n});l.push(o||function(){})}})),h.update()},forceUpdate:function(){if(!c){var t=a.elements,e=t.reference,i=t.popper;if(pi(e,i)){a.rects={reference:di(e,$e(i),"fixed"===a.options.strategy),popper:Ce(i)},a.reset=!1,a.placement=a.options.placement,a.orderedModifiers.forEach((function(t){return a.modifiersData[t.name]=Object.assign({},t.data)}));for(var n=0;n<a.orderedModifiers.length;n++)if(!0!==a.reset){var s=a.orderedModifiers[n],o=s.fn,r=s.options,l=void 0===r?{}:r,d=s.name;"function"==typeof o&&(a=o({state:a,options:l,name:d,instance:h})||a)}else a.reset=!1,n=-1}}},update:(s=function(){return new Promise((function(t){h.forceUpdate(),t(a)}))},function(){return r||(r=new Promise((function(t){Promise.resolve().then((function(){r=void 0,t(s())}))}))),r}),destroy:function(){d(),c=!0}};if(!pi(t,e))return h;function d(){l.forEach((function(t){return t()})),l=[]}return h.setOptions(i).then((function(t){!c&&i.onFirstUpdate&&i.onFirstUpdate(t)})),h}}var gi=mi(),_i=mi({defaultModifiers:[Re,ci,Be,_e]}),bi=mi({defaultModifiers:[Re,ci,Be,_e,li,si,hi,je,ai]});const vi=Object.freeze(Object.defineProperty({__proto__:null,afterMain:ae,afterRead:se,afterWrite:he,applyStyles:_e,arrow:je,auto:Kt,basePlacements:Qt,beforeMain:oe,beforeRead:ie,beforeWrite:le,bottom:Rt,clippingParents:Ut,computeStyles:Be,createPopper:bi,createPopperBase:gi,createPopperLite:_i,detectOverflow:ii,end:Yt,eventListeners:Re,flip:si,hide:ai,left:Vt,main:re,modifierPhases:de,offset:li,placements:ee,popper:Jt,popperGenerator:mi,popperOffsets:ci,preventOverflow:hi,read:ne,reference:Zt,right:qt,start:Xt,top:zt,variationPlacements:te,viewport:Gt,write:ce},Symbol.toStringTag,{value:"Module"})),yi="dropdown",wi=".bs.dropdown",Ai=".data-api",Ei="ArrowUp",Ti="ArrowDown",Ci=`hide${wi}`,Oi=`hidden${wi}`,xi=`show${wi}`,ki=`shown${wi}`,Li=`click${wi}${Ai}`,Si=`keydown${wi}${Ai}`,Di=`keyup${wi}${Ai}`,$i="show",Ii='[data-bs-toggle="dropdown"]:not(.disabled):not(:disabled)',Ni=`${Ii}.${$i}`,Pi=".dropdown-menu",Mi=p()?"top-end":"top-start",ji=p()?"top-start":"top-end",Fi=p()?"bottom-end":"bottom-start",Hi=p()?"bottom-start":"bottom-end",Wi=p()?"left-start":"right-start",Bi=p()?"right-start":"left-start",zi={autoClose:!0,boundary:"clippingParents",display:"dynamic",offset:[0,2],popperConfig:null,reference:"toggle"},Ri={autoClose:"(boolean|string)",boundary:"(string|element)",display:"string",offset:"(array|string|function)",popperConfig:"(null|object|function)",reference:"(string|element|object)"};class qi extends W{constructor(t,e){super(t,e),this._popper=null,this._parent=this._element.parentNode,this._menu=z.next(this._element,Pi)[0]||z.prev(this._element,Pi)[0]||z.findOne(Pi,this._parent),this._inNavbar=this._detectNavbar()}static get Default(){return zi}static get DefaultType(){return Ri}static get NAME(){return yi}toggle(){return this._isShown()?this.hide():this.show()}show(){if(l(this._element)||this._isShown())return;const t={relatedTarget:this._element};if(!N.trigger(this._element,xi,t).defaultPrevented){if(this._createPopper(),"ontouchstart"in document.documentElement&&!this._parent.closest(".navbar-nav"))for(const t of[].concat(...document.body.children))N.on(t,"mouseover",h);this._element.focus(),this._element.setAttribute("aria-expanded",!0),this._menu.classList.add($i),this._element.classList.add($i),N.trigger(this._element,ki,t)}}hide(){if(l(this._element)||!this._isShown())return;const t={relatedTarget:this._element};this._completeHide(t)}dispose(){this._popper&&this._popper.destroy(),super.dispose()}update(){this._inNavbar=this._detectNavbar(),this._popper&&this._popper.update()}_completeHide(t){if(!N.trigger(this._element,Ci,t).defaultPrevented){if("ontouchstart"in document.documentElement)for(const t of[].concat(...document.body.children))N.off(t,"mouseover",h);this._popper&&this._popper.destroy(),this._menu.classList.remove($i),this._element.classList.remove($i),this._element.setAttribute("aria-expanded","false"),F.removeDataAttribute(this._menu,"popper"),N.trigger(this._element,Oi,t)}}_getConfig(t){if("object"==typeof(t=super._getConfig(t)).reference&&!o(t.reference)&&"function"!=typeof t.reference.getBoundingClientRect)throw new TypeError(`${yi.toUpperCase()}: Option "reference" provided type "object" without a required "getBoundingClientRect" method.`);return t}_createPopper(){if(void 0===vi)throw new TypeError("Bootstrap's dropdowns require Popper (https://popper.js.org)");let t=this._element;"parent"===this._config.reference?t=this._parent:o(this._config.reference)?t=r(this._config.reference):"object"==typeof this._config.reference&&(t=this._config.reference);const e=this._getPopperConfig();this._popper=bi(t,this._menu,e)}_isShown(){return this._menu.classList.contains($i)}_getPlacement(){const t=this._parent;if(t.classList.contains("dropend"))return Wi;if(t.classList.contains("dropstart"))return Bi;if(t.classList.contains("dropup-center"))return"top";if(t.classList.contains("dropdown-center"))return"bottom";const e="end"===getComputedStyle(this._menu).getPropertyValue("--bs-position").trim();return t.classList.contains("dropup")?e?ji:Mi:e?Hi:Fi}_detectNavbar(){return null!==this._element.closest(".navbar")}_getOffset(){const{offset:t}=this._config;return"string"==typeof t?t.split(",").map((t=>Number.parseInt(t,10))):"function"==typeof t?e=>t(e,this._element):t}_getPopperConfig(){const t={placement:this._getPlacement(),modifiers:[{name:"preventOverflow",options:{boundary:this._config.boundary}},{name:"offset",options:{offset:this._getOffset()}}]};return(this._inNavbar||"static"===this._config.display)&&(F.setDataAttribute(this._menu,"popper","static"),t.modifiers=[{name:"applyStyles",enabled:!1}]),{...t,...g(this._config.popperConfig,[t])}}_selectMenuItem({key:t,target:e}){const i=z.find(".dropdown-menu .dropdown-item:not(.disabled):not(:disabled)",this._menu).filter((t=>a(t)));i.length&&b(i,e,t===Ti,!i.includes(e)).focus()}static jQueryInterface(t){return this.each((function(){const e=qi.getOrCreateInstance(this,t);if("string"==typeof t){if(void 0===e[t])throw new TypeError(`No method named "${t}"`);e[t]()}}))}static clearMenus(t){if(2===t.button||"keyup"===t.type&&"Tab"!==t.key)return;const e=z.find(Ni);for(const i of e){const e=qi.getInstance(i);if(!e||!1===e._config.autoClose)continue;const n=t.composedPath(),s=n.includes(e._menu);if(n.includes(e._element)||"inside"===e._config.autoClose&&!s||"outside"===e._config.autoClose&&s)continue;if(e._menu.contains(t.target)&&("keyup"===t.type&&"Tab"===t.key||/input|select|option|textarea|form/i.test(t.target.tagName)))continue;const o={relatedTarget:e._element};"click"===t.type&&(o.clickEvent=t),e._completeHide(o)}}static dataApiKeydownHandler(t){const e=/input|textarea/i.test(t.target.tagName),i="Escape"===t.key,n=[Ei,Ti].includes(t.key);if(!n&&!i)return;if(e&&!i)return;t.preventDefault();const s=this.matches(Ii)?this:z.prev(this,Ii)[0]||z.next(this,Ii)[0]||z.findOne(Ii,t.delegateTarget.parentNode),o=qi.getOrCreateInstance(s);if(n)return t.stopPropagation(),o.show(),void o._selectMenuItem(t);o._isShown()&&(t.stopPropagation(),o.hide(),s.focus())}}N.on(document,Si,Ii,qi.dataApiKeydownHandler),N.on(document,Si,Pi,qi.dataApiKeydownHandler),N.on(document,Li,qi.clearMenus),N.on(document,Di,qi.clearMenus),N.on(document,Li,Ii,(function(t){t.preventDefault(),qi.getOrCreateInstance(this).toggle()})),m(qi);const Vi="backdrop",Ki="show",Qi=`mousedown.bs.${Vi}`,Xi={className:"modal-backdrop",clickCallback:null,isAnimated:!1,isVisible:!0,rootElement:"body"},Yi={className:"string",clickCallback:"(function|null)",isAnimated:"boolean",isVisible:"boolean",rootElement:"(element|string)"};class Ui extends H{constructor(t){super(),this._config=this._getConfig(t),this._isAppended=!1,this._element=null}static get Default(){return Xi}static get DefaultType(){return Yi}static get NAME(){return Vi}show(t){if(!this._config.isVisible)return void g(t);this._append();const e=this._getElement();this._config.isAnimated&&d(e),e.classList.add(Ki),this._emulateAnimation((()=>{g(t)}))}hide(t){this._config.isVisible?(this._getElement().classList.remove(Ki),this._emulateAnimation((()=>{this.dispose(),g(t)}))):g(t)}dispose(){this._isAppended&&(N.off(this._element,Qi),this._element.remove(),this._isAppended=!1)}_getElement(){if(!this._element){const t=document.createElement("div");t.className=this._config.className,this._config.isAnimated&&t.classList.add("fade"),this._element=t}return this._element}_configAfterMerge(t){return t.rootElement=r(t.rootElement),t}_append(){if(this._isAppended)return;const t=this._getElement();this._config.rootElement.append(t),N.on(t,Qi,(()=>{g(this._config.clickCallback)})),this._isAppended=!0}_emulateAnimation(t){_(t,this._getElement(),this._config.isAnimated)}}const Gi=".bs.focustrap",Ji=`focusin${Gi}`,Zi=`keydown.tab${Gi}`,tn="backward",en={autofocus:!0,trapElement:null},nn={autofocus:"boolean",trapElement:"element"};class sn extends H{constructor(t){super(),this._config=this._getConfig(t),this._isActive=!1,this._lastTabNavDirection=null}static get Default(){return en}static get DefaultType(){return nn}static get NAME(){return"focustrap"}activate(){this._isActive||(this._config.autofocus&&this._config.trapElement.focus(),N.off(document,Gi),N.on(document,Ji,(t=>this._handleFocusin(t))),N.on(document,Zi,(t=>this._handleKeydown(t))),this._isActive=!0)}deactivate(){this._isActive&&(this._isActive=!1,N.off(document,Gi))}_handleFocusin(t){const{trapElement:e}=this._config;if(t.target===document||t.target===e||e.contains(t.target))return;const i=z.focusableChildren(e);0===i.length?e.focus():this._lastTabNavDirection===tn?i[i.length-1].focus():i[0].focus()}_handleKeydown(t){"Tab"===t.key&&(this._lastTabNavDirection=t.shiftKey?tn:"forward")}}const on=".fixed-top, .fixed-bottom, .is-fixed, .sticky-top",rn=".sticky-top",an="padding-right",ln="margin-right";class cn{constructor(){this._element=document.body}getWidth(){const t=document.documentElement.clientWidth;return Math.abs(window.innerWidth-t)}hide(){const t=this.getWidth();this._disableOverFlow(),this._setElementAttributes(this._element,an,(e=>e+t)),this._setElementAttributes(on,an,(e=>e+t)),this._setElementAttributes(rn,ln,(e=>e-t))}reset(){this._resetElementAttributes(this._element,"overflow"),this._resetElementAttributes(this._element,an),this._resetElementAttributes(on,an),this._resetElementAttributes(rn,ln)}isOverflowing(){return this.getWidth()>0}_disableOverFlow(){this._saveInitialAttribute(this._element,"overflow"),this._element.style.overflow="hidden"}_setElementAttributes(t,e,i){const n=this.getWidth();this._applyManipulationCallback(t,(t=>{if(t!==this._element&&window.innerWidth>t.clientWidth+n)return;this._saveInitialAttribute(t,e);const s=window.getComputedStyle(t).getPropertyValue(e);t.style.setProperty(e,`${i(Number.parseFloat(s))}px`)}))}_saveInitialAttribute(t,e){const i=t.style.getPropertyValue(e);i&&F.setDataAttribute(t,e,i)}_resetElementAttributes(t,e){this._applyManipulationCallback(t,(t=>{const i=F.getDataAttribute(t,e);null!==i?(F.removeDataAttribute(t,e),t.style.setProperty(e,i)):t.style.removeProperty(e)}))}_applyManipulationCallback(t,e){if(o(t))e(t);else for(const i of z.find(t,this._element))e(i)}}const hn=".bs.modal",dn=`hide${hn}`,un=`hidePrevented${hn}`,fn=`hidden${hn}`,pn=`show${hn}`,mn=`shown${hn}`,gn=`resize${hn}`,_n=`click.dismiss${hn}`,bn=`mousedown.dismiss${hn}`,vn=`keydown.dismiss${hn}`,yn=`click${hn}.data-api`,wn="modal-open",An="show",En="modal-static",Tn={backdrop:!0,focus:!0,keyboard:!0},Cn={backdrop:"(boolean|string)",focus:"boolean",keyboard:"boolean"};class On extends W{constructor(t,e){super(t,e),this._dialog=z.findOne(".modal-dialog",this._element),this._backdrop=this._initializeBackDrop(),this._focustrap=this._initializeFocusTrap(),this._isShown=!1,this._isTransitioning=!1,this._scrollBar=new cn,this._addEventListeners()}static get Default(){return Tn}static get DefaultType(){return Cn}static get NAME(){return"modal"}toggle(t){return this._isShown?this.hide():this.show(t)}show(t){this._isShown||this._isTransitioning||N.trigger(this._element,pn,{relatedTarget:t}).defaultPrevented||(this._isShown=!0,this._isTransitioning=!0,this._scrollBar.hide(),document.body.classList.add(wn),this._adjustDialog(),this._backdrop.show((()=>this._showElement(t))))}hide(){this._isShown&&!this._isTransitioning&&(N.trigger(this._element,dn).defaultPrevented||(this._isShown=!1,this._isTransitioning=!0,this._focustrap.deactivate(),this._element.classList.remove(An),this._queueCallback((()=>this._hideModal()),this._element,this._isAnimated())))}dispose(){N.off(window,hn),N.off(this._dialog,hn),this._backdrop.dispose(),this._focustrap.deactivate(),super.dispose()}handleUpdate(){this._adjustDialog()}_initializeBackDrop(){return new Ui({isVisible:Boolean(this._config.backdrop),isAnimated:this._isAnimated()})}_initializeFocusTrap(){return new sn({trapElement:this._element})}_showElement(t){document.body.contains(this._element)||document.body.append(this._element),this._element.style.display="block",this._element.removeAttribute("aria-hidden"),this._element.setAttribute("aria-modal",!0),this._element.setAttribute("role","dialog"),this._element.scrollTop=0;const e=z.findOne(".modal-body",this._dialog);e&&(e.scrollTop=0),d(this._element),this._element.classList.add(An),this._queueCallback((()=>{this._config.focus&&this._focustrap.activate(),this._isTransitioning=!1,N.trigger(this._element,mn,{relatedTarget:t})}),this._dialog,this._isAnimated())}_addEventListeners(){N.on(this._element,vn,(t=>{"Escape"===t.key&&(this._config.keyboard?this.hide():this._triggerBackdropTransition())})),N.on(window,gn,(()=>{this._isShown&&!this._isTransitioning&&this._adjustDialog()})),N.on(this._element,bn,(t=>{N.one(this._element,_n,(e=>{this._element===t.target&&this._element===e.target&&("static"!==this._config.backdrop?this._config.backdrop&&this.hide():this._triggerBackdropTransition())}))}))}_hideModal(){this._element.style.display="none",this._element.setAttribute("aria-hidden",!0),this._element.removeAttribute("aria-modal"),this._element.removeAttribute("role"),this._isTransitioning=!1,this._backdrop.hide((()=>{document.body.classList.remove(wn),this._resetAdjustments(),this._scrollBar.reset(),N.trigger(this._element,fn)}))}_isAnimated(){return this._element.classList.contains("fade")}_triggerBackdropTransition(){if(N.trigger(this._element,un).defaultPrevented)return;const t=this._element.scrollHeight>document.documentElement.clientHeight,e=this._element.style.overflowY;"hidden"===e||this._element.classList.contains(En)||(t||(this._element.style.overflowY="hidden"),this._element.classList.add(En),this._queueCallback((()=>{this._element.classList.remove(En),this._queueCallback((()=>{this._element.style.overflowY=e}),this._dialog)}),this._dialog),this._element.focus())}_adjustDialog(){const t=this._element.scrollHeight>document.documentElement.clientHeight,e=this._scrollBar.getWidth(),i=e>0;if(i&&!t){const t=p()?"paddingLeft":"paddingRight";this._element.style[t]=`${e}px`}if(!i&&t){const t=p()?"paddingRight":"paddingLeft";this._element.style[t]=`${e}px`}}_resetAdjustments(){this._element.style.paddingLeft="",this._element.style.paddingRight=""}static jQueryInterface(t,e){return this.each((function(){const i=On.getOrCreateInstance(this,t);if("string"==typeof t){if(void 0===i[t])throw new TypeError(`No method named "${t}"`);i[t](e)}}))}}N.on(document,yn,'[data-bs-toggle="modal"]',(function(t){const e=z.getElementFromSelector(this);["A","AREA"].includes(this.tagName)&&t.preventDefault(),N.one(e,pn,(t=>{t.defaultPrevented||N.one(e,fn,(()=>{a(this)&&this.focus()}))}));const i=z.findOne(".modal.show");i&&On.getInstance(i).hide(),On.getOrCreateInstance(e).toggle(this)})),R(On),m(On);const xn=".bs.offcanvas",kn=".data-api",Ln=`load${xn}${kn}`,Sn="show",Dn="showing",$n="hiding",In=".offcanvas.show",Nn=`show${xn}`,Pn=`shown${xn}`,Mn=`hide${xn}`,jn=`hidePrevented${xn}`,Fn=`hidden${xn}`,Hn=`resize${xn}`,Wn=`click${xn}${kn}`,Bn=`keydown.dismiss${xn}`,zn={backdrop:!0,keyboard:!0,scroll:!1},Rn={backdrop:"(boolean|string)",keyboard:"boolean",scroll:"boolean"};class qn extends W{constructor(t,e){super(t,e),this._isShown=!1,this._backdrop=this._initializeBackDrop(),this._focustrap=this._initializeFocusTrap(),this._addEventListeners()}static get Default(){return zn}static get DefaultType(){return Rn}static get NAME(){return"offcanvas"}toggle(t){return this._isShown?this.hide():this.show(t)}show(t){this._isShown||N.trigger(this._element,Nn,{relatedTarget:t}).defaultPrevented||(this._isShown=!0,this._backdrop.show(),this._config.scroll||(new cn).hide(),this._element.setAttribute("aria-modal",!0),this._element.setAttribute("role","dialog"),this._element.classList.add(Dn),this._queueCallback((()=>{this._config.scroll&&!this._config.backdrop||this._focustrap.activate(),this._element.classList.add(Sn),this._element.classList.remove(Dn),N.trigger(this._element,Pn,{relatedTarget:t})}),this._element,!0))}hide(){this._isShown&&(N.trigger(this._element,Mn).defaultPrevented||(this._focustrap.deactivate(),this._element.blur(),this._isShown=!1,this._element.classList.add($n),this._backdrop.hide(),this._queueCallback((()=>{this._element.classList.remove(Sn,$n),this._element.removeAttribute("aria-modal"),this._element.removeAttribute("role"),this._config.scroll||(new cn).reset(),N.trigger(this._element,Fn)}),this._element,!0)))}dispose(){this._backdrop.dispose(),this._focustrap.deactivate(),super.dispose()}_initializeBackDrop(){const t=Boolean(this._config.backdrop);return new Ui({className:"offcanvas-backdrop",isVisible:t,isAnimated:!0,rootElement:this._element.parentNode,clickCallback:t?()=>{"static"!==this._config.backdrop?this.hide():N.trigger(this._element,jn)}:null})}_initializeFocusTrap(){return new sn({trapElement:this._element})}_addEventListeners(){N.on(this._element,Bn,(t=>{"Escape"===t.key&&(this._config.keyboard?this.hide():N.trigger(this._element,jn))}))}static jQueryInterface(t){return this.each((function(){const e=qn.getOrCreateInstance(this,t);if("string"==typeof t){if(void 0===e[t]||t.startsWith("_")||"constructor"===t)throw new TypeError(`No method named "${t}"`);e[t](this)}}))}}N.on(document,Wn,'[data-bs-toggle="offcanvas"]',(function(t){const e=z.getElementFromSelector(this);if(["A","AREA"].includes(this.tagName)&&t.preventDefault(),l(this))return;N.one(e,Fn,(()=>{a(this)&&this.focus()}));const i=z.findOne(In);i&&i!==e&&qn.getInstance(i).hide(),qn.getOrCreateInstance(e).toggle(this)})),N.on(window,Ln,(()=>{for(const t of z.find(In))qn.getOrCreateInstance(t).show()})),N.on(window,Hn,(()=>{for(const t of z.find("[aria-modal][class*=show][class*=offcanvas-]"))"fixed"!==getComputedStyle(t).position&&qn.getOrCreateInstance(t).hide()})),R(qn),m(qn);const Vn={"*":["class","dir","id","lang","role",/^aria-[\w-]*$/i],a:["target","href","title","rel"],area:[],b:[],br:[],col:[],code:[],div:[],em:[],hr:[],h1:[],h2:[],h3:[],h4:[],h5:[],h6:[],i:[],img:["src","srcset","alt","title","width","height"],li:[],ol:[],p:[],pre:[],s:[],small:[],span:[],sub:[],sup:[],strong:[],u:[],ul:[]},Kn=new Set(["background","cite","href","itemtype","longdesc","poster","src","xlink:href"]),Qn=/^(?!javascript:)(?:[a-z0-9+.-]+:|[^&:/?#]*(?:[/?#]|$))/i,Xn=(t,e)=>{const i=t.nodeName.toLowerCase();return e.includes(i)?!Kn.has(i)||Boolean(Qn.test(t.nodeValue)):e.filter((t=>t instanceof RegExp)).some((t=>t.test(i)))},Yn={allowList:Vn,content:{},extraClass:"",html:!1,sanitize:!0,sanitizeFn:null,template:"<div></div>"},Un={allowList:"object",content:"object",extraClass:"(string|function)",html:"boolean",sanitize:"boolean",sanitizeFn:"(null|function)",template:"string"},Gn={entry:"(string|element|function|null)",selector:"(string|element)"};class Jn extends H{constructor(t){super(),this._config=this._getConfig(t)}static get Default(){return Yn}static get DefaultType(){return Un}static get NAME(){return"TemplateFactory"}getContent(){return Object.values(this._config.content).map((t=>this._resolvePossibleFunction(t))).filter(Boolean)}hasContent(){return this.getContent().length>0}changeContent(t){return this._checkContent(t),this._config.content={...this._config.content,...t},this}toHtml(){const t=document.createElement("div");t.innerHTML=this._maybeSanitize(this._config.template);for(const[e,i]of Object.entries(this._config.content))this._setContent(t,i,e);const e=t.children[0],i=this._resolvePossibleFunction(this._config.extraClass);return i&&e.classList.add(...i.split(" ")),e}_typeCheckConfig(t){super._typeCheckConfig(t),this._checkContent(t.content)}_checkContent(t){for(const[e,i]of Object.entries(t))super._typeCheckConfig({selector:e,entry:i},Gn)}_setContent(t,e,i){const n=z.findOne(i,t);n&&((e=this._resolvePossibleFunction(e))?o(e)?this._putElementInTemplate(r(e),n):this._config.html?n.innerHTML=this._maybeSanitize(e):n.textContent=e:n.remove())}_maybeSanitize(t){return this._config.sanitize?function(t,e,i){if(!t.length)return t;if(i&&"function"==typeof i)return i(t);const n=(new window.DOMParser).parseFromString(t,"text/html"),s=[].concat(...n.body.querySelectorAll("*"));for(const t of s){const i=t.nodeName.toLowerCase();if(!Object.keys(e).includes(i)){t.remove();continue}const n=[].concat(...t.attributes),s=[].concat(e["*"]||[],e[i]||[]);for(const e of n)Xn(e,s)||t.removeAttribute(e.nodeName)}return n.body.innerHTML}(t,this._config.allowList,this._config.sanitizeFn):t}_resolvePossibleFunction(t){return g(t,[this])}_putElementInTemplate(t,e){if(this._config.html)return e.innerHTML="",void e.append(t);e.textContent=t.textContent}}const Zn=new Set(["sanitize","allowList","sanitizeFn"]),ts="fade",es="show",is=".modal",ns="hide.bs.modal",ss="hover",os="focus",rs={AUTO:"auto",TOP:"top",RIGHT:p()?"left":"right",BOTTOM:"bottom",LEFT:p()?"right":"left"},as={allowList:Vn,animation:!0,boundary:"clippingParents",container:!1,customClass:"",delay:0,fallbackPlacements:["top","right","bottom","left"],html:!1,offset:[0,6],placement:"top",popperConfig:null,sanitize:!0,sanitizeFn:null,selector:!1,template:'<div class="tooltip" role="tooltip"><div class="tooltip-arrow"></div><div class="tooltip-inner"></div></div>',title:"",trigger:"hover focus"},ls={allowList:"object",animation:"boolean",boundary:"(string|element)",container:"(string|element|boolean)",customClass:"(string|function)",delay:"(number|object)",fallbackPlacements:"array",html:"boolean",offset:"(array|string|function)",placement:"(string|function)",popperConfig:"(null|object|function)",sanitize:"boolean",sanitizeFn:"(null|function)",selector:"(string|boolean)",template:"string",title:"(string|element|function)",trigger:"string"};class cs extends W{constructor(t,e){if(void 0===vi)throw new TypeError("Bootstrap's tooltips require Popper (https://popper.js.org)");super(t,e),this._isEnabled=!0,this._timeout=0,this._isHovered=null,this._activeTrigger={},this._popper=null,this._templateFactory=null,this._newContent=null,this.tip=null,this._setListeners(),this._config.selector||this._fixTitle()}static get Default(){return as}static get DefaultType(){return ls}static get NAME(){return"tooltip"}enable(){this._isEnabled=!0}disable(){this._isEnabled=!1}toggleEnabled(){this._isEnabled=!this._isEnabled}toggle(){this._isEnabled&&(this._activeTrigger.click=!this._activeTrigger.click,this._isShown()?this._leave():this._enter())}dispose(){clearTimeout(this._timeout),N.off(this._element.closest(is),ns,this._hideModalHandler),this._element.getAttribute("data-bs-original-title")&&this._element.setAttribute("title",this._element.getAttribute("data-bs-original-title")),this._disposePopper(),super.dispose()}show(){if("none"===this._element.style.display)throw new Error("Please use show on visible elements");if(!this._isWithContent()||!this._isEnabled)return;const t=N.trigger(this._element,this.constructor.eventName("show")),e=(c(this._element)||this._element.ownerDocument.documentElement).contains(this._element);if(t.defaultPrevented||!e)return;this._disposePopper();const i=this._getTipElement();this._element.setAttribute("aria-describedby",i.getAttribute("id"));const{container:n}=this._config;if(this._element.ownerDocument.documentElement.contains(this.tip)||(n.append(i),N.trigger(this._element,this.constructor.eventName("inserted"))),this._popper=this._createPopper(i),i.classList.add(es),"ontouchstart"in document.documentElement)for(const t of[].concat(...document.body.children))N.on(t,"mouseover",h);this._queueCallback((()=>{N.trigger(this._element,this.constructor.eventName("shown")),!1===this._isHovered&&this._leave(),this._isHovered=!1}),this.tip,this._isAnimated())}hide(){if(this._isShown()&&!N.trigger(this._element,this.constructor.eventName("hide")).defaultPrevented){if(this._getTipElement().classList.remove(es),"ontouchstart"in document.documentElement)for(const t of[].concat(...document.body.children))N.off(t,"mouseover",h);this._activeTrigger.click=!1,this._activeTrigger[os]=!1,this._activeTrigger[ss]=!1,this._isHovered=null,this._queueCallback((()=>{this._isWithActiveTrigger()||(this._isHovered||this._disposePopper(),this._element.removeAttribute("aria-describedby"),N.trigger(this._element,this.constructor.eventName("hidden")))}),this.tip,this._isAnimated())}}update(){this._popper&&this._popper.update()}_isWithContent(){return Boolean(this._getTitle())}_getTipElement(){return this.tip||(this.tip=this._createTipElement(this._newContent||this._getContentForTemplate())),this.tip}_createTipElement(t){const e=this._getTemplateFactory(t).toHtml();if(!e)return null;e.classList.remove(ts,es),e.classList.add(`bs-${this.constructor.NAME}-auto`);const i=(t=>{do{t+=Math.floor(1e6*Math.random())}while(document.getElementById(t));return t})(this.constructor.NAME).toString();return e.setAttribute("id",i),this._isAnimated()&&e.classList.add(ts),e}setContent(t){this._newContent=t,this._isShown()&&(this._disposePopper(),this.show())}_getTemplateFactory(t){return this._templateFactory?this._templateFactory.changeContent(t):this._templateFactory=new Jn({...this._config,content:t,extraClass:this._resolvePossibleFunction(this._config.customClass)}),this._templateFactory}_getContentForTemplate(){return{".tooltip-inner":this._getTitle()}}_getTitle(){return this._resolvePossibleFunction(this._config.title)||this._element.getAttribute("data-bs-original-title")}_initializeOnDelegatedTarget(t){return this.constructor.getOrCreateInstance(t.delegateTarget,this._getDelegateConfig())}_isAnimated(){return this._config.animation||this.tip&&this.tip.classList.contains(ts)}_isShown(){return this.tip&&this.tip.classList.contains(es)}_createPopper(t){const e=g(this._config.placement,[this,t,this._element]),i=rs[e.toUpperCase()];return bi(this._element,t,this._getPopperConfig(i))}_getOffset(){const{offset:t}=this._config;return"string"==typeof t?t.split(",").map((t=>Number.parseInt(t,10))):"function"==typeof t?e=>t(e,this._element):t}_resolvePossibleFunction(t){return g(t,[this._element])}_getPopperConfig(t){const e={placement:t,modifiers:[{name:"flip",options:{fallbackPlacements:this._config.fallbackPlacements}},{name:"offset",options:{offset:this._getOffset()}},{name:"preventOverflow",options:{boundary:this._config.boundary}},{name:"arrow",options:{element:`.${this.constructor.NAME}-arrow`}},{name:"preSetPlacement",enabled:!0,phase:"beforeMain",fn:t=>{this._getTipElement().setAttribute("data-popper-placement",t.state.placement)}}]};return{...e,...g(this._config.popperConfig,[e])}}_setListeners(){const t=this._config.trigger.split(" ");for(const e of t)if("click"===e)N.on(this._element,this.constructor.eventName("click"),this._config.selector,(t=>{this._initializeOnDelegatedTarget(t).toggle()}));else if("manual"!==e){const t=e===ss?this.constructor.eventName("mouseenter"):this.constructor.eventName("focusin"),i=e===ss?this.constructor.eventName("mouseleave"):this.constructor.eventName("focusout");N.on(this._element,t,this._config.selector,(t=>{const e=this._initializeOnDelegatedTarget(t);e._activeTrigger["focusin"===t.type?os:ss]=!0,e._enter()})),N.on(this._element,i,this._config.selector,(t=>{const e=this._initializeOnDelegatedTarget(t);e._activeTrigger["focusout"===t.type?os:ss]=e._element.contains(t.relatedTarget),e._leave()}))}this._hideModalHandler=()=>{this._element&&this.hide()},N.on(this._element.closest(is),ns,this._hideModalHandler)}_fixTitle(){const t=this._element.getAttribute("title");t&&(this._element.getAttribute("aria-label")||this._element.textContent.trim()||this._element.setAttribute("aria-label",t),this._element.setAttribute("data-bs-original-title",t),this._element.removeAttribute("title"))}_enter(){this._isShown()||this._isHovered?this._isHovered=!0:(this._isHovered=!0,this._setTimeout((()=>{this._isHovered&&this.show()}),this._config.delay.show))}_leave(){this._isWithActiveTrigger()||(this._isHovered=!1,this._setTimeout((()=>{this._isHovered||this.hide()}),this._config.delay.hide))}_setTimeout(t,e){clearTimeout(this._timeout),this._timeout=setTimeout(t,e)}_isWithActiveTrigger(){return Object.values(this._activeTrigger).includes(!0)}_getConfig(t){const e=F.getDataAttributes(this._element);for(const t of Object.keys(e))Zn.has(t)&&delete e[t];return t={...e,..."object"==typeof t&&t?t:{}},t=this._mergeConfigObj(t),t=this._configAfterMerge(t),this._typeCheckConfig(t),t}_configAfterMerge(t){return t.container=!1===t.container?document.body:r(t.container),"number"==typeof t.delay&&(t.delay={show:t.delay,hide:t.delay}),"number"==typeof t.title&&(t.title=t.title.toString()),"number"==typeof t.content&&(t.content=t.content.toString()),t}_getDelegateConfig(){const t={};for(const[e,i]of Object.entries(this._config))this.constructor.Default[e]!==i&&(t[e]=i);return t.selector=!1,t.trigger="manual",t}_disposePopper(){this._popper&&(this._popper.destroy(),this._popper=null),this.tip&&(this.tip.remove(),this.tip=null)}static jQueryInterface(t){return this.each((function(){const e=cs.getOrCreateInstance(this,t);if("string"==typeof t){if(void 0===e[t])throw new TypeError(`No method named "${t}"`);e[t]()}}))}}m(cs);const hs={...cs.Default,content:"",offset:[0,8],placement:"right",template:'<div class="popover" role="tooltip"><div class="popover-arrow"></div><h3 class="popover-header"></h3><div class="popover-body"></div></div>',trigger:"click"},ds={...cs.DefaultType,content:"(null|string|element|function)"};class us extends cs{static get Default(){return hs}static get DefaultType(){return ds}static get NAME(){return"popover"}_isWithContent(){return this._getTitle()||this._getContent()}_getContentForTemplate(){return{".popover-header":this._getTitle(),".popover-body":this._getContent()}}_getContent(){return this._resolvePossibleFunction(this._config.content)}static jQueryInterface(t){return this.each((function(){const e=us.getOrCreateInstance(this,t);if("string"==typeof t){if(void 0===e[t])throw new TypeError(`No method named "${t}"`);e[t]()}}))}}m(us);const fs=".bs.scrollspy",ps=`activate${fs}`,ms=`click${fs}`,gs=`load${fs}.data-api`,_s="active",bs="[href]",vs=".nav-link",ys=`${vs}, .nav-item > ${vs}, .list-group-item`,ws={offset:null,rootMargin:"0px 0px -25%",smoothScroll:!1,target:null,threshold:[.1,.5,1]},As={offset:"(number|null)",rootMargin:"string",smoothScroll:"boolean",target:"element",threshold:"array"};class Es extends W{constructor(t,e){super(t,e),this._targetLinks=new Map,this._observableSections=new Map,this._rootElement="visible"===getComputedStyle(this._element).overflowY?null:this._element,this._activeTarget=null,this._observer=null,this._previousScrollData={visibleEntryTop:0,parentScrollTop:0},this.refresh()}static get Default(){return ws}static get DefaultType(){return As}static get NAME(){return"scrollspy"}refresh(){this._initializeTargetsAndObservables(),this._maybeEnableSmoothScroll(),this._observer?this._observer.disconnect():this._observer=this._getNewObserver();for(const t of this._observableSections.values())this._observer.observe(t)}dispose(){this._observer.disconnect(),super.dispose()}_configAfterMerge(t){return t.target=r(t.target)||document.body,t.rootMargin=t.offset?`${t.offset}px 0px -30%`:t.rootMargin,"string"==typeof t.threshold&&(t.threshold=t.threshold.split(",").map((t=>Number.parseFloat(t)))),t}_maybeEnableSmoothScroll(){this._config.smoothScroll&&(N.off(this._config.target,ms),N.on(this._config.target,ms,bs,(t=>{const e=this._observableSections.get(t.target.hash);if(e){t.preventDefault();const i=this._rootElement||window,n=e.offsetTop-this._element.offsetTop;if(i.scrollTo)return void i.scrollTo({top:n,behavior:"smooth"});i.scrollTop=n}})))}_getNewObserver(){const t={root:this._rootElement,threshold:this._config.threshold,rootMargin:this._config.rootMargin};return new IntersectionObserver((t=>this._observerCallback(t)),t)}_observerCallback(t){const e=t=>this._targetLinks.get(`#${t.target.id}`),i=t=>{this._previousScrollData.visibleEntryTop=t.target.offsetTop,this._process(e(t))},n=(this._rootElement||document.documentElement).scrollTop,s=n>=this._previousScrollData.parentScrollTop;this._previousScrollData.parentScrollTop=n;for(const o of t){if(!o.isIntersecting){this._activeTarget=null,this._clearActiveClass(e(o));continue}const t=o.target.offsetTop>=this._previousScrollData.visibleEntryTop;if(s&&t){if(i(o),!n)return}else s||t||i(o)}}_initializeTargetsAndObservables(){this._targetLinks=new Map,this._observableSections=new Map;const t=z.find(bs,this._config.target);for(const e of t){if(!e.hash||l(e))continue;const t=z.findOne(decodeURI(e.hash),this._element);a(t)&&(this._targetLinks.set(decodeURI(e.hash),e),this._observableSections.set(e.hash,t))}}_process(t){this._activeTarget!==t&&(this._clearActiveClass(this._config.target),this._activeTarget=t,t.classList.add(_s),this._activateParents(t),N.trigger(this._element,ps,{relatedTarget:t}))}_activateParents(t){if(t.classList.contains("dropdown-item"))z.findOne(".dropdown-toggle",t.closest(".dropdown")).classList.add(_s);else for(const e of z.parents(t,".nav, .list-group"))for(const t of z.prev(e,ys))t.classList.add(_s)}_clearActiveClass(t){t.classList.remove(_s);const e=z.find(`${bs}.${_s}`,t);for(const t of e)t.classList.remove(_s)}static jQueryInterface(t){return this.each((function(){const e=Es.getOrCreateInstance(this,t);if("string"==typeof t){if(void 0===e[t]||t.startsWith("_")||"constructor"===t)throw new TypeError(`No method named "${t}"`);e[t]()}}))}}N.on(window,gs,(()=>{for(const t of z.find('[data-bs-spy="scroll"]'))Es.getOrCreateInstance(t)})),m(Es);const Ts=".bs.tab",Cs=`hide${Ts}`,Os=`hidden${Ts}`,xs=`show${Ts}`,ks=`shown${Ts}`,Ls=`click${Ts}`,Ss=`keydown${Ts}`,Ds=`load${Ts}`,$s="ArrowLeft",Is="ArrowRight",Ns="ArrowUp",Ps="ArrowDown",Ms="Home",js="End",Fs="active",Hs="fade",Ws="show",Bs=":not(.dropdown-toggle)",zs='[data-bs-toggle="tab"], [data-bs-toggle="pill"], [data-bs-toggle="list"]',Rs=`.nav-link${Bs}, .list-group-item${Bs}, [role="tab"]${Bs}, ${zs}`,qs=`.${Fs}[data-bs-toggle="tab"], .${Fs}[data-bs-toggle="pill"], .${Fs}[data-bs-toggle="list"]`;class Vs extends W{constructor(t){super(t),this._parent=this._element.closest('.list-group, .nav, [role="tablist"]'),this._parent&&(this._setInitialAttributes(this._parent,this._getChildren()),N.on(this._element,Ss,(t=>this._keydown(t))))}static get NAME(){return"tab"}show(){const t=this._element;if(this._elemIsActive(t))return;const e=this._getActiveElem(),i=e?N.trigger(e,Cs,{relatedTarget:t}):null;N.trigger(t,xs,{relatedTarget:e}).defaultPrevented||i&&i.defaultPrevented||(this._deactivate(e,t),this._activate(t,e))}_activate(t,e){t&&(t.classList.add(Fs),this._activate(z.getElementFromSelector(t)),this._queueCallback((()=>{"tab"===t.getAttribute("role")?(t.removeAttribute("tabindex"),t.setAttribute("aria-selected",!0),this._toggleDropDown(t,!0),N.trigger(t,ks,{relatedTarget:e})):t.classList.add(Ws)}),t,t.classList.contains(Hs)))}_deactivate(t,e){t&&(t.classList.remove(Fs),t.blur(),this._deactivate(z.getElementFromSelector(t)),this._queueCallback((()=>{"tab"===t.getAttribute("role")?(t.setAttribute("aria-selected",!1),t.setAttribute("tabindex","-1"),this._toggleDropDown(t,!1),N.trigger(t,Os,{relatedTarget:e})):t.classList.remove(Ws)}),t,t.classList.contains(Hs)))}_keydown(t){if(![$s,Is,Ns,Ps,Ms,js].includes(t.key))return;t.stopPropagation(),t.preventDefault();const e=this._getChildren().filter((t=>!l(t)));let i;if([Ms,js].includes(t.key))i=e[t.key===Ms?0:e.length-1];else{const n=[Is,Ps].includes(t.key);i=b(e,t.target,n,!0)}i&&(i.focus({preventScroll:!0}),Vs.getOrCreateInstance(i).show())}_getChildren(){return z.find(Rs,this._parent)}_getActiveElem(){return this._getChildren().find((t=>this._elemIsActive(t)))||null}_setInitialAttributes(t,e){this._setAttributeIfNotExists(t,"role","tablist");for(const t of e)this._setInitialAttributesOnChild(t)}_setInitialAttributesOnChild(t){t=this._getInnerElement(t);const e=this._elemIsActive(t),i=this._getOuterElement(t);t.setAttribute("aria-selected",e),i!==t&&this._setAttributeIfNotExists(i,"role","presentation"),e||t.setAttribute("tabindex","-1"),this._setAttributeIfNotExists(t,"role","tab"),this._setInitialAttributesOnTargetPanel(t)}_setInitialAttributesOnTargetPanel(t){const e=z.getElementFromSelector(t);e&&(this._setAttributeIfNotExists(e,"role","tabpanel"),t.id&&this._setAttributeIfNotExists(e,"aria-labelledby",`${t.id}`))}_toggleDropDown(t,e){const i=this._getOuterElement(t);if(!i.classList.contains("dropdown"))return;const n=(t,n)=>{const s=z.findOne(t,i);s&&s.classList.toggle(n,e)};n(".dropdown-toggle",Fs),n(".dropdown-menu",Ws),i.setAttribute("aria-expanded",e)}_setAttributeIfNotExists(t,e,i){t.hasAttribute(e)||t.setAttribute(e,i)}_elemIsActive(t){return t.classList.contains(Fs)}_getInnerElement(t){return t.matches(Rs)?t:z.findOne(Rs,t)}_getOuterElement(t){return t.closest(".nav-item, .list-group-item")||t}static jQueryInterface(t){return this.each((function(){const e=Vs.getOrCreateInstance(this);if("string"==typeof t){if(void 0===e[t]||t.startsWith("_")||"constructor"===t)throw new TypeError(`No method named "${t}"`);e[t]()}}))}}N.on(document,Ls,zs,(function(t){["A","AREA"].includes(this.tagName)&&t.preventDefault(),l(this)||Vs.getOrCreateInstance(this).show()})),N.on(window,Ds,(()=>{for(const t of z.find(qs))Vs.getOrCreateInstance(t)})),m(Vs);const Ks=".bs.toast",Qs=`mouseover${Ks}`,Xs=`mouseout${Ks}`,Ys=`focusin${Ks}`,Us=`focusout${Ks}`,Gs=`hide${Ks}`,Js=`hidden${Ks}`,Zs=`show${Ks}`,to=`shown${Ks}`,eo="hide",io="show",no="showing",so={animation:"boolean",autohide:"boolean",delay:"number"},oo={animation:!0,autohide:!0,delay:5e3};class ro extends W{constructor(t,e){super(t,e),this._timeout=null,this._hasMouseInteraction=!1,this._hasKeyboardInteraction=!1,this._setListeners()}static get Default(){return oo}static get DefaultType(){return so}static get NAME(){return"toast"}show(){N.trigger(this._element,Zs).defaultPrevented||(this._clearTimeout(),this._config.animation&&this._element.classList.add("fade"),this._element.classList.remove(eo),d(this._element),this._element.classList.add(io,no),this._queueCallback((()=>{this._element.classList.remove(no),N.trigger(this._element,to),this._maybeScheduleHide()}),this._element,this._config.animation))}hide(){this.isShown()&&(N.trigger(this._element,Gs).defaultPrevented||(this._element.classList.add(no),this._queueCallback((()=>{this._element.classList.add(eo),this._element.classList.remove(no,io),N.trigger(this._element,Js)}),this._element,this._config.animation)))}dispose(){this._clearTimeout(),this.isShown()&&this._element.classList.remove(io),super.dispose()}isShown(){return this._element.classList.contains(io)}_maybeScheduleHide(){this._config.autohide&&(this._hasMouseInteraction||this._hasKeyboardInteraction||(this._timeout=setTimeout((()=>{this.hide()}),this._config.delay)))}_onInteraction(t,e){switch(t.type){case"mouseover":case"mouseout":this._hasMouseInteraction=e;break;case"focusin":case"focusout":this._hasKeyboardInteraction=e}if(e)return void this._clearTimeout();const i=t.relatedTarget;this._element===i||this._element.contains(i)||this._maybeScheduleHide()}_setListeners(){N.on(this._element,Qs,(t=>this._onInteraction(t,!0))),N.on(this._element,Xs,(t=>this._onInteraction(t,!1))),N.on(this._element,Ys,(t=>this._onInteraction(t,!0))),N.on(this._element,Us,(t=>this._onInteraction(t,!1)))}_clearTimeout(){clearTimeout(this._timeout),this._timeout=null}static jQueryInterface(t){return this.each((function(){const e=ro.getOrCreateInstance(this,t);if("string"==typeof t){if(void 0===e[t])throw new TypeError(`No method named "${t}"`);e[t](this)}}))}}return R(ro),m(ro),{Alert:Q,Button:Y,Carousel:xt,Collapse:Bt,Dropdown:qi,Modal:On,Offcanvas:qn,Popover:us,ScrollSpy:Es,Tab:Vs,Toast:ro,Tooltip:cs}}));
//# sourceMappingURL=bootstrap.bundle.min.js.map
```
