## Appendix for "Challenges and lessons learned from preliminary modeling of with-in herd transmission of highly pathogenic avian influenza H_5_N_1_ in dairy cattle": clipboard.min.js

```
/*!
 * clipboard.js v2.0.11
 * https://clipboardjs.com/
 *
 * Licensed MIT © Zeno Rocha
 */
!function(t,e){"object"==typeof exports&&"object"==typeof module?module.exports=e():"function"==typeof define&&define.amd?define([],e):"object"==typeof exports?exports.ClipboardJS=e():t.ClipboardJS=e()}(this,function(){return n={686:function(t,e,n){"use strict";n.d(e,{default:function(){return b}});var e=n(279),i=n.n(e),e=n(370),u=n.n(e),e=n(817),r=n.n(e);function c(t){try{return document.execCommand(t)}catch(t){return}}var a=function(t){t=r()(t);return c("cut"),t};function o(t,e){var n,o,t=(n=t,o="rtl"===document.documentElement.getAttribute("dir"),(t=document.createElement("textarea")).style.fontSize="12pt",t.style.border="0",t.style.padding="0",t.style.margin="0",t.style.position="absolute",t.style[o?"right":"left"]="-9999px",o=window.pageYOffset||document.documentElement.scrollTop,t.style.top="".concat(o,"px"),t.setAttribute("readonly",""),t.value=n,t);return e.container.appendChild(t),e=r()(t),c("copy"),t.remove(),e}var f=function(t){var e=1<arguments.length&&void 0!==arguments[1]?arguments[1]:{container:document.body},n="";return"string"==typeof t?n=o(t,e):t instanceof HTMLInputElement&&!["text","search","url","tel","password"].includes(null==t?void 0:t.type)?n=o(t.value,e):(n=r()(t),c("copy")),n};function l(t){return(l="function"==typeof Symbol&&"symbol"==typeof Symbol.iterator?function(t){return typeof t}:function(t){return t&&"function"==typeof Symbol&&t.constructor===Symbol&&t!==Symbol.prototype?"symbol":typeof t})(t)}var s=function(){var t=0<arguments.length&&void 0!==arguments[0]?arguments[0]:{},e=t.action,n=void 0===e?"copy":e,o=t.container,e=t.target,t=t.text;if("copy"!==n&&"cut"!==n)throw new Error('Invalid "action" value, use either "copy" or "cut"');if(void 0!==e){if(!e||"object"!==l(e)||1!==e.nodeType)throw new Error('Invalid "target" value, use a valid Element');if("copy"===n&&e.hasAttribute("disabled"))throw new Error('Invalid "target" attribute. Please use "readonly" instead of "disabled" attribute');if("cut"===n&&(e.hasAttribute("readonly")||e.hasAttribute("disabled")))throw new Error('Invalid "target" attribute. You can\'t cut text from elements with "readonly" or "disabled" attributes')}return t?f(t,{container:o}):e?"cut"===n?a(e):f(e,{container:o}):void 0};function p(t){return(p="function"==typeof Symbol&&"symbol"==typeof Symbol.iterator?function(t){return typeof t}:function(t){return t&&"function"==typeof Symbol&&t.constructor===Symbol&&t!==Symbol.prototype?"symbol":typeof t})(t)}function d(t,e){for(var n=0;n<e.length;n++){var o=e[n];o.enumerable=o.enumerable||!1,o.configurable=!0,"value"in o&&(o.writable=!0),Object.defineProperty(t,o.key,o)}}function y(t,e){return(y=Object.setPrototypeOf||function(t,e){return t.__proto__=e,t})(t,e)}function h(n){var o=function(){if("undefined"==typeof Reflect||!Reflect.construct)return!1;if(Reflect.construct.sham)return!1;if("function"==typeof Proxy)return!0;try{return Date.prototype.toString.call(Reflect.construct(Date,[],function(){})),!0}catch(t){return!1}}();return function(){var t,e=v(n);return t=o?(t=v(this).constructor,Reflect.construct(e,arguments,t)):e.apply(this,arguments),e=this,!(t=t)||"object"!==p(t)&&"function"!=typeof t?function(t){if(void 0!==t)return t;throw new ReferenceError("this hasn't been initialised - super() hasn't been called")}(e):t}}function v(t){return(v=Object.setPrototypeOf?Object.getPrototypeOf:function(t){return t.__proto__||Object.getPrototypeOf(t)})(t)}function m(t,e){t="data-clipboard-".concat(t);if(e.hasAttribute(t))return e.getAttribute(t)}var b=function(){!function(t,e){if("function"!=typeof e&&null!==e)throw new TypeError("Super expression must either be null or a function");t.prototype=Object.create(e&&e.prototype,{constructor:{value:t,writable:!0,configurable:!0}}),e&&y(t,e)}(r,i());var t,e,n,o=h(r);function r(t,e){var n;return function(t){if(!(t instanceof r))throw new TypeError("Cannot call a class as a function")}(this),(n=o.call(this)).resolveOptions(e),n.listenClick(t),n}return t=r,n=[{key:"copy",value:function(t){var e=1<arguments.length&&void 0!==arguments[1]?arguments[1]:{container:document.body};return f(t,e)}},{key:"cut",value:function(t){return a(t)}},{key:"isSupported",value:function(){var t=0<arguments.length&&void 0!==arguments[0]?arguments[0]:["copy","cut"],t="string"==typeof t?[t]:t,e=!!document.queryCommandSupported;return t.forEach(function(t){e=e&&!!document.queryCommandSupported(t)}),e}}],(e=[{key:"resolveOptions",value:function(){var t=0<arguments.length&&void 0!==arguments[0]?arguments[0]:{};this.action="function"==typeof t.action?t.action:this.defaultAction,this.target="function"==typeof t.target?t.target:this.defaultTarget,this.text="function"==typeof t.text?t.text:this.defaultText,this.container="object"===p(t.container)?t.container:document.body}},{key:"listenClick",value:function(t){var e=this;this.listener=u()(t,"click",function(t){return e.onClick(t)})}},{key:"onClick",value:function(t){var e=t.delegateTarget||t.currentTarget,n=this.action(e)||"copy",t=s({action:n,container:this.container,target:this.target(e),text:this.text(e)});this.emit(t?"success":"error",{action:n,text:t,trigger:e,clearSelection:function(){e&&e.focus(),window.getSelection().removeAllRanges()}})}},{key:"defaultAction",value:function(t){return m("action",t)}},{key:"defaultTarget",value:function(t){t=m("target",t);if(t)return document.querySelector(t)}},{key:"defaultText",value:function(t){return m("text",t)}},{key:"destroy",value:function(){this.listener.destroy()}}])&&d(t.prototype,e),n&&d(t,n),r}()},828:function(t){var e;"undefined"==typeof Element||Element.prototype.matches||((e=Element.prototype).matches=e.matchesSelector||e.mozMatchesSelector||e.msMatchesSelector||e.oMatchesSelector||e.webkitMatchesSelector),t.exports=function(t,e){for(;t&&9!==t.nodeType;){if("function"==typeof t.matches&&t.matches(e))return t;t=t.parentNode}}},438:function(t,e,n){var u=n(828);function i(t,e,n,o,r){var i=function(e,n,t,o){return function(t){t.delegateTarget=u(t.target,n),t.delegateTarget&&o.call(e,t)}}.apply(this,arguments);return t.addEventListener(n,i,r),{destroy:function(){t.removeEventListener(n,i,r)}}}t.exports=function(t,e,n,o,r){return"function"==typeof t.addEventListener?i.apply(null,arguments):"function"==typeof n?i.bind(null,document).apply(null,arguments):("string"==typeof t&&(t=document.querySelectorAll(t)),Array.prototype.map.call(t,function(t){return i(t,e,n,o,r)}))}},879:function(t,n){n.node=function(t){return void 0!==t&&t instanceof HTMLElement&&1===t.nodeType},n.nodeList=function(t){var e=Object.prototype.toString.call(t);return void 0!==t&&("[object NodeList]"===e||"[object HTMLCollection]"===e)&&"length"in t&&(0===t.length||n.node(t[0]))},n.string=function(t){return"string"==typeof t||t instanceof String},n.fn=function(t){return"[object Function]"===Object.prototype.toString.call(t)}},370:function(t,e,n){var f=n(879),l=n(438);t.exports=function(t,e,n){if(!t&&!e&&!n)throw new Error("Missing required arguments");if(!f.string(e))throw new TypeError("Second argument must be a String");if(!f.fn(n))throw new TypeError("Third argument must be a Function");if(f.node(t))return c=e,a=n,(u=t).addEventListener(c,a),{destroy:function(){u.removeEventListener(c,a)}};if(f.nodeList(t))return o=t,r=e,i=n,Array.prototype.forEach.call(o,function(t){t.addEventListener(r,i)}),{destroy:function(){Array.prototype.forEach.call(o,function(t){t.removeEventListener(r,i)})}};if(f.string(t))return t=t,e=e,n=n,l(document.body,t,e,n);throw new TypeError("First argument must be a String, HTMLElement, HTMLCollection, or NodeList");var o,r,i,u,c,a}},817:function(t){t.exports=function(t){var e,n="SELECT"===t.nodeName?(t.focus(),t.value):"INPUT"===t.nodeName||"TEXTAREA"===t.nodeName?((e=t.hasAttribute("readonly"))||t.setAttribute("readonly",""),t.select(),t.setSelectionRange(0,t.value.length),e||t.removeAttribute("readonly"),t.value):(t.hasAttribute("contenteditable")&&t.focus(),n=window.getSelection(),(e=document.createRange()).selectNodeContents(t),n.removeAllRanges(),n.addRange(e),n.toString());return n}},279:function(t){function e(){}e.prototype={on:function(t,e,n){var o=this.e||(this.e={});return(o[t]||(o[t]=[])).push({fn:e,ctx:n}),this},once:function(t,e,n){var o=this;function r(){o.off(t,r),e.apply(n,arguments)}return r._=e,this.on(t,r,n)},emit:function(t){for(var e=[].slice.call(arguments,1),n=((this.e||(this.e={}))[t]||[]).slice(),o=0,r=n.length;o<r;o++)n[o].fn.apply(n[o].ctx,e);return this},off:function(t,e){var n=this.e||(this.e={}),o=n[t],r=[];if(o&&e)for(var i=0,u=o.length;i<u;i++)o[i].fn!==e&&o[i].fn._!==e&&r.push(o[i]);return r.length?n[t]=r:delete n[t],this}},t.exports=e,t.exports.TinyEmitter=e}},r={},o.n=function(t){var e=t&&t.__esModule?function(){return t.default}:function(){return t};return o.d(e,{a:e}),e},o.d=function(t,e){for(var n in e)o.o(e,n)&&!o.o(t,n)&&Object.defineProperty(t,n,{enumerable:!0,get:e[n]})},o.o=function(t,e){return Object.prototype.hasOwnProperty.call(t,e)},o(686).default;function o(t){if(r[t])return r[t].exports;var e=r[t]={exports:{}};return n[t](e,e.exports,o),e.exports}var n,r});
```
