## Appendix for "Challenges and lessons learned from preliminary modeling of with-in herd transmission of highly pathogenic avian influenza H_5_N_1_ in dairy cattle": anchor.min.js

```
// @license magnet:?xt=urn:btih:d3d9a9a6595521f9666a5e94cc830dab83b65699&dn=expat.txt Expat
//
// AnchorJS - v5.0.0 - 2023-01-18
// https://www.bryanbraun.com/anchorjs/
// Copyright (c) 2023 Bryan Braun; Licensed MIT
//
// @license magnet:?xt=urn:btih:d3d9a9a6595521f9666a5e94cc830dab83b65699&dn=expat.txt Expat
!function(A,e){"use strict";"function"==typeof define&&define.amd?define([],e):"object"==typeof module&&module.exports?module.exports=e():(A.AnchorJS=e(),A.anchors=new A.AnchorJS)}(globalThis,function(){"use strict";return function(A){function u(A){A.icon=Object.prototype.hasOwnProperty.call(A,"icon")?A.icon:"",A.visible=Object.prototype.hasOwnProperty.call(A,"visible")?A.visible:"hover",A.placement=Object.prototype.hasOwnProperty.call(A,"placement")?A.placement:"right",A.ariaLabel=Object.prototype.hasOwnProperty.call(A,"ariaLabel")?A.ariaLabel:"Anchor",A.class=Object.prototype.hasOwnProperty.call(A,"class")?A.class:"",A.base=Object.prototype.hasOwnProperty.call(A,"base")?A.base:"",A.truncate=Object.prototype.hasOwnProperty.call(A,"truncate")?Math.floor(A.truncate):64,A.titleText=Object.prototype.hasOwnProperty.call(A,"titleText")?A.titleText:""}function d(A){var e;if("string"==typeof A||A instanceof String)e=[].slice.call(document.querySelectorAll(A));else{if(!(Array.isArray(A)||A instanceof NodeList))throw new TypeError("The selector provided to AnchorJS was invalid.");e=[].slice.call(A)}return e}this.options=A||{},this.elements=[],u(this.options),this.add=function(A){var e,t,o,i,n,s,a,r,l,c,h,p=[];if(u(this.options),0!==(e=d(A=A||"h2, h3, h4, h5, h6")).length){for(null===document.head.querySelector("style.anchorjs")&&((A=document.createElement("style")).className="anchorjs",A.appendChild(document.createTextNode("")),void 0===(h=document.head.querySelector('[rel="stylesheet"],style'))?document.head.appendChild(A):document.head.insertBefore(A,h),A.sheet.insertRule(".anchorjs-link{opacity:0;text-decoration:none;-webkit-font-smoothing:antialiased;-moz-osx-font-smoothing:grayscale}",A.sheet.cssRules.length),A.sheet.insertRule(":hover>.anchorjs-link,.anchorjs-link:focus{opacity:1}",A.sheet.cssRules.length),A.sheet.insertRule("[data-anchorjs-icon]::after{content:attr(data-anchorjs-icon)}",A.sheet.cssRules.length),A.sheet.insertRule('@font-face{font-family:anchorjs-icons;src:url(data:n/a;base64,AAEAAAALAIAAAwAwT1MvMg8yG2cAAAE4AAAAYGNtYXDp3gC3AAABpAAAAExnYXNwAAAAEAAAA9wAAAAIZ2x5ZlQCcfwAAAH4AAABCGhlYWQHFvHyAAAAvAAAADZoaGVhBnACFwAAAPQAAAAkaG10eASAADEAAAGYAAAADGxvY2EACACEAAAB8AAAAAhtYXhwAAYAVwAAARgAAAAgbmFtZQGOH9cAAAMAAAAAunBvc3QAAwAAAAADvAAAACAAAQAAAAEAAHzE2p9fDzz1AAkEAAAAAADRecUWAAAAANQA6R8AAAAAAoACwAAAAAgAAgAAAAAAAAABAAADwP/AAAACgAAA/9MCrQABAAAAAAAAAAAAAAAAAAAAAwABAAAAAwBVAAIAAAAAAAIAAAAAAAAAAAAAAAAAAAAAAAMCQAGQAAUAAAKZAswAAACPApkCzAAAAesAMwEJAAAAAAAAAAAAAAAAAAAAARAAAAAAAAAAAAAAAAAAAAAAQAAg//0DwP/AAEADwABAAAAAAQAAAAAAAAAAAAAAIAAAAAAAAAIAAAACgAAxAAAAAwAAAAMAAAAcAAEAAwAAABwAAwABAAAAHAAEADAAAAAIAAgAAgAAACDpy//9//8AAAAg6cv//f///+EWNwADAAEAAAAAAAAAAAAAAAAACACEAAEAAAAAAAAAAAAAAAAxAAACAAQARAKAAsAAKwBUAAABIiYnJjQ3NzY2MzIWFxYUBwcGIicmNDc3NjQnJiYjIgYHBwYUFxYUBwYGIwciJicmNDc3NjIXFhQHBwYUFxYWMzI2Nzc2NCcmNDc2MhcWFAcHBgYjARQGDAUtLXoWOR8fORYtLTgKGwoKCjgaGg0gEhIgDXoaGgkJBQwHdR85Fi0tOAobCgoKOBoaDSASEiANehoaCQkKGwotLXoWOR8BMwUFLYEuehYXFxYugC44CQkKGwo4GkoaDQ0NDXoaShoKGwoFBe8XFi6ALjgJCQobCjgaShoNDQ0NehpKGgobCgoKLYEuehYXAAAADACWAAEAAAAAAAEACAAAAAEAAAAAAAIAAwAIAAEAAAAAAAMACAAAAAEAAAAAAAQACAAAAAEAAAAAAAUAAQALAAEAAAAAAAYACAAAAAMAAQQJAAEAEAAMAAMAAQQJAAIABgAcAAMAAQQJAAMAEAAMAAMAAQQJAAQAEAAMAAMAAQQJAAUAAgAiAAMAAQQJAAYAEAAMYW5jaG9yanM0MDBAAGEAbgBjAGgAbwByAGoAcwA0ADAAMABAAAAAAwAAAAAAAAAAAAAAAAAAAAAAAAAAAAAAAAAAAAAAAAABAAH//wAP) format("truetype")}',A.sheet.cssRules.length)),h=document.querySelectorAll("[id]"),t=[].map.call(h,function(A){return A.id}),i=0;i<e.length;i++)if(this.hasAnchorJSLink(e[i]))p.push(i);else{if(e[i].hasAttribute("id"))o=e[i].getAttribute("id");else if(e[i].hasAttribute("data-anchor-id"))o=e[i].getAttribute("data-anchor-id");else{for(r=a=this.urlify(e[i].textContent),s=0;n=t.indexOf(r=void 0!==n?a+"-"+s:r),s+=1,-1!==n;);n=void 0,t.push(r),e[i].setAttribute("id",r),o=r}(l=document.createElement("a")).className="anchorjs-link "+this.options.class,l.setAttribute("aria-label",this.options.ariaLabel),l.setAttribute("data-anchorjs-icon",this.options.icon),this.options.titleText&&(l.title=this.options.titleText),c=document.querySelector("base")?window.location.pathname+window.location.search:"",c=this.options.base||c,l.href=c+"#"+o,"always"===this.options.visible&&(l.style.opacity="1"),""===this.options.icon&&(l.style.font="1em/1 anchorjs-icons","left"===this.options.placement)&&(l.style.lineHeight="inherit"),"left"===this.options.placement?(l.style.position="absolute",l.style.marginLeft="-1.25em",l.style.paddingRight=".25em",l.style.paddingLeft=".25em",e[i].insertBefore(l,e[i].firstChild)):(l.style.marginLeft=".1875em",l.style.paddingRight=".1875em",l.style.paddingLeft=".1875em",e[i].appendChild(l))}for(i=0;i<p.length;i++)e.splice(p[i]-i,1);this.elements=this.elements.concat(e)}return this},this.remove=function(A){for(var e,t,o=d(A),i=0;i<o.length;i++)(t=o[i].querySelector(".anchorjs-link"))&&(-1!==(e=this.elements.indexOf(o[i]))&&this.elements.splice(e,1),o[i].removeChild(t));return this},this.removeAll=function(){this.remove(this.elements)},this.urlify=function(A){var e=document.createElement("textarea");return e.innerHTML=A,A=e.value,this.options.truncate||u(this.options),A.trim().replace(/'/gi,"").replace(/[& +$,:;=?@"#{}|^~[`%!'<>\]./()*\\\n\t\b\v\u00A0]/g,"-").replace(/-{2,}/g,"-").substring(0,this.options.truncate).replace(/^-+|-+$/gm,"").toLowerCase()},this.hasAnchorJSLink=function(A){var e=A.firstChild&&-1<(" "+A.firstChild.className+" ").indexOf(" anchorjs-link "),A=A.lastChild&&-1<(" "+A.lastChild.className+" ").indexOf(" anchorjs-link ");return e||A||!1}}});
// @license-end
```
