## Appendix for "Challenges and lessons learned from preliminary modeling of with-in herd transmission of highly pathogenic avian influenza H_5_N_1_ in dairy cattle": popper.min.js

```
/**
 * @popperjs/core v2.11.7 - MIT License
 */

!function(e,t){"object"==typeof exports&&"undefined"!=typeof module?t(exports):"function"==typeof define&&define.amd?define(["exports"],t):t((e="undefined"!=typeof globalThis?globalThis:e||self).Popper={})}(this,(function(e){"use strict";function t(e){if(null==e)return window;if("[object Window]"!==e.toString()){var t=e.ownerDocument;return t&&t.defaultView||window}return e}function n(e){return e instanceof t(e).Element||e instanceof Element}function r(e){return e instanceof t(e).HTMLElement||e instanceof HTMLElement}function o(e){return"undefined"!=typeof ShadowRoot&&(e instanceof t(e).ShadowRoot||e instanceof ShadowRoot)}var i=Math.max,a=Math.min,s=Math.round;function f(){var e=navigator.userAgentData;return null!=e&&e.brands&&Array.isArray(e.brands)?e.brands.map((function(e){return e.brand+"/"+e.version})).join(" "):navigator.userAgent}function c(){return!/^((?!chrome|android).)*safari/i.test(f())}function p(e,o,i){void 0===o&&(o=!1),void 0===i&&(i=!1);var a=e.getBoundingClientRect(),f=1,p=1;o&&r(e)&&(f=e.offsetWidth>0&&s(a.width)/e.offsetWidth||1,p=e.offsetHeight>0&&s(a.height)/e.offsetHeight||1);var u=(n(e)?t(e):window).visualViewport,l=!c()&&i,d=(a.left+(l&&u?u.offsetLeft:0))/f,h=(a.top+(l&&u?u.offsetTop:0))/p,m=a.width/f,v=a.height/p;return{width:m,height:v,top:h,right:d+m,bottom:h+v,left:d,x:d,y:h}}function u(e){var n=t(e);return{scrollLeft:n.pageXOffset,scrollTop:n.pageYOffset}}function l(e){return e?(e.nodeName||"").toLowerCase():null}function d(e){return((n(e)?e.ownerDocument:e.document)||window.document).documentElement}function h(e){return p(d(e)).left+u(e).scrollLeft}function m(e){return t(e).getComputedStyle(e)}function v(e){var t=m(e),n=t.overflow,r=t.overflowX,o=t.overflowY;return/auto|scroll|overlay|hidden/.test(n+o+r)}function y(e,n,o){void 0===o&&(o=!1);var i,a,f=r(n),c=r(n)&&function(e){var t=e.getBoundingClientRect(),n=s(t.width)/e.offsetWidth||1,r=s(t.height)/e.offsetHeight||1;return 1!==n||1!==r}(n),m=d(n),y=p(e,c,o),g={scrollLeft:0,scrollTop:0},b={x:0,y:0};return(f||!f&&!o)&&(("body"!==l(n)||v(m))&&(g=(i=n)!==t(i)&&r(i)?{scrollLeft:(a=i).scrollLeft,scrollTop:a.scrollTop}:u(i)),r(n)?((b=p(n,!0)).x+=n.clientLeft,b.y+=n.clientTop):m&&(b.x=h(m))),{x:y.left+g.scrollLeft-b.x,y:y.top+g.scrollTop-b.y,width:y.width,height:y.height}}function g(e){var t=p(e),n=e.offsetWidth,r=e.offsetHeight;return Math.abs(t.width-n)<=1&&(n=t.width),Math.abs(t.height-r)<=1&&(r=t.height),{x:e.offsetLeft,y:e.offsetTop,width:n,height:r}}function b(e){return"html"===l(e)?e:e.assignedSlot||e.parentNode||(o(e)?e.host:null)||d(e)}function x(e){return["html","body","#document"].indexOf(l(e))>=0?e.ownerDocument.body:r(e)&&v(e)?e:x(b(e))}function w(e,n){var r;void 0===n&&(n=[]);var o=x(e),i=o===(null==(r=e.ownerDocument)?void 0:r.body),a=t(o),s=i?[a].concat(a.visualViewport||[],v(o)?o:[]):o,f=n.concat(s);return i?f:f.concat(w(b(s)))}function O(e){return["table","td","th"].indexOf(l(e))>=0}function j(e){return r(e)&&"fixed"!==m(e).position?e.offsetParent:null}function E(e){for(var n=t(e),i=j(e);i&&O(i)&&"static"===m(i).position;)i=j(i);return i&&("html"===l(i)||"body"===l(i)&&"static"===m(i).position)?n:i||function(e){var t=/firefox/i.test(f());if(/Trident/i.test(f())&&r(e)&&"fixed"===m(e).position)return null;var n=b(e);for(o(n)&&(n=n.host);r(n)&&["html","body"].indexOf(l(n))<0;){var i=m(n);if("none"!==i.transform||"none"!==i.perspective||"paint"===i.contain||-1!==["transform","perspective"].indexOf(i.willChange)||t&&"filter"===i.willChange||t&&i.filter&&"none"!==i.filter)return n;n=n.parentNode}return null}(e)||n}var D="top",A="bottom",L="right",P="left",M="auto",k=[D,A,L,P],W="start",B="end",H="viewport",T="popper",R=k.reduce((function(e,t){return e.concat([t+"-"+W,t+"-"+B])}),[]),S=[].concat(k,[M]).reduce((function(e,t){return e.concat([t,t+"-"+W,t+"-"+B])}),[]),V=["beforeRead","read","afterRead","beforeMain","main","afterMain","beforeWrite","write","afterWrite"];function q(e){var t=new Map,n=new Set,r=[];function o(e){n.add(e.name),[].concat(e.requires||[],e.requiresIfExists||[]).forEach((function(e){if(!n.has(e)){var r=t.get(e);r&&o(r)}})),r.push(e)}return e.forEach((function(e){t.set(e.name,e)})),e.forEach((function(e){n.has(e.name)||o(e)})),r}function C(e){return e.split("-")[0]}function N(e,t){var n=t.getRootNode&&t.getRootNode();if(e.contains(t))return!0;if(n&&o(n)){var r=t;do{if(r&&e.isSameNode(r))return!0;r=r.parentNode||r.host}while(r)}return!1}function I(e){return Object.assign({},e,{left:e.x,top:e.y,right:e.x+e.width,bottom:e.y+e.height})}function _(e,r,o){return r===H?I(function(e,n){var r=t(e),o=d(e),i=r.visualViewport,a=o.clientWidth,s=o.clientHeight,f=0,p=0;if(i){a=i.width,s=i.height;var u=c();(u||!u&&"fixed"===n)&&(f=i.offsetLeft,p=i.offsetTop)}return{width:a,height:s,x:f+h(e),y:p}}(e,o)):n(r)?function(e,t){var n=p(e,!1,"fixed"===t);return n.top=n.top+e.clientTop,n.left=n.left+e.clientLeft,n.bottom=n.top+e.clientHeight,n.right=n.left+e.clientWidth,n.width=e.clientWidth,n.height=e.clientHeight,n.x=n.left,n.y=n.top,n}(r,o):I(function(e){var t,n=d(e),r=u(e),o=null==(t=e.ownerDocument)?void 0:t.body,a=i(n.scrollWidth,n.clientWidth,o?o.scrollWidth:0,o?o.clientWidth:0),s=i(n.scrollHeight,n.clientHeight,o?o.scrollHeight:0,o?o.clientHeight:0),f=-r.scrollLeft+h(e),c=-r.scrollTop;return"rtl"===m(o||n).direction&&(f+=i(n.clientWidth,o?o.clientWidth:0)-a),{width:a,height:s,x:f,y:c}}(d(e)))}function F(e,t,o,s){var f="clippingParents"===t?function(e){var t=w(b(e)),o=["absolute","fixed"].indexOf(m(e).position)>=0&&r(e)?E(e):e;return n(o)?t.filter((function(e){return n(e)&&N(e,o)&&"body"!==l(e)})):[]}(e):[].concat(t),c=[].concat(f,[o]),p=c[0],u=c.reduce((function(t,n){var r=_(e,n,s);return t.top=i(r.top,t.top),t.right=a(r.right,t.right),t.bottom=a(r.bottom,t.bottom),t.left=i(r.left,t.left),t}),_(e,p,s));return u.width=u.right-u.left,u.height=u.bottom-u.top,u.x=u.left,u.y=u.top,u}function U(e){return e.split("-")[1]}function z(e){return["top","bottom"].indexOf(e)>=0?"x":"y"}function X(e){var t,n=e.reference,r=e.element,o=e.placement,i=o?C(o):null,a=o?U(o):null,s=n.x+n.width/2-r.width/2,f=n.y+n.height/2-r.height/2;switch(i){case D:t={x:s,y:n.y-r.height};break;case A:t={x:s,y:n.y+n.height};break;case L:t={x:n.x+n.width,y:f};break;case P:t={x:n.x-r.width,y:f};break;default:t={x:n.x,y:n.y}}var c=i?z(i):null;if(null!=c){var p="y"===c?"height":"width";switch(a){case W:t[c]=t[c]-(n[p]/2-r[p]/2);break;case B:t[c]=t[c]+(n[p]/2-r[p]/2)}}return t}function Y(e){return Object.assign({},{top:0,right:0,bottom:0,left:0},e)}function G(e,t){return t.reduce((function(t,n){return t[n]=e,t}),{})}function J(e,t){void 0===t&&(t={});var r=t,o=r.placement,i=void 0===o?e.placement:o,a=r.strategy,s=void 0===a?e.strategy:a,f=r.boundary,c=void 0===f?"clippingParents":f,u=r.rootBoundary,l=void 0===u?H:u,h=r.elementContext,m=void 0===h?T:h,v=r.altBoundary,y=void 0!==v&&v,g=r.padding,b=void 0===g?0:g,x=Y("number"!=typeof b?b:G(b,k)),w=m===T?"reference":T,O=e.rects.popper,j=e.elements[y?w:m],E=F(n(j)?j:j.contextElement||d(e.elements.popper),c,l,s),P=p(e.elements.reference),M=X({reference:P,element:O,strategy:"absolute",placement:i}),W=I(Object.assign({},O,M)),B=m===T?W:P,R={top:E.top-B.top+x.top,bottom:B.bottom-E.bottom+x.bottom,left:E.left-B.left+x.left,right:B.right-E.right+x.right},S=e.modifiersData.offset;if(m===T&&S){var V=S[i];Object.keys(R).forEach((function(e){var t=[L,A].indexOf(e)>=0?1:-1,n=[D,A].indexOf(e)>=0?"y":"x";R[e]+=V[n]*t}))}return R}var K={placement:"bottom",modifiers:[],strategy:"absolute"};function Q(){for(var e=arguments.length,t=new Array(e),n=0;n<e;n++)t[n]=arguments[n];return!t.some((function(e){return!(e&&"function"==typeof e.getBoundingClientRect)}))}function Z(e){void 0===e&&(e={});var t=e,r=t.defaultModifiers,o=void 0===r?[]:r,i=t.defaultOptions,a=void 0===i?K:i;return function(e,t,r){void 0===r&&(r=a);var i,s,f={placement:"bottom",orderedModifiers:[],options:Object.assign({},K,a),modifiersData:{},elements:{reference:e,popper:t},attributes:{},styles:{}},c=[],p=!1,u={state:f,setOptions:function(r){var i="function"==typeof r?r(f.options):r;l(),f.options=Object.assign({},a,f.options,i),f.scrollParents={reference:n(e)?w(e):e.contextElement?w(e.contextElement):[],popper:w(t)};var s,p,d=function(e){var t=q(e);return V.reduce((function(e,n){return e.concat(t.filter((function(e){return e.phase===n})))}),[])}((s=[].concat(o,f.options.modifiers),p=s.reduce((function(e,t){var n=e[t.name];return e[t.name]=n?Object.assign({},n,t,{options:Object.assign({},n.options,t.options),data:Object.assign({},n.data,t.data)}):t,e}),{}),Object.keys(p).map((function(e){return p[e]}))));return f.orderedModifiers=d.filter((function(e){return e.enabled})),f.orderedModifiers.forEach((function(e){var t=e.name,n=e.options,r=void 0===n?{}:n,o=e.effect;if("function"==typeof o){var i=o({state:f,name:t,instance:u,options:r}),a=function(){};c.push(i||a)}})),u.update()},forceUpdate:function(){if(!p){var e=f.elements,t=e.reference,n=e.popper;if(Q(t,n)){f.rects={reference:y(t,E(n),"fixed"===f.options.strategy),popper:g(n)},f.reset=!1,f.placement=f.options.placement,f.orderedModifiers.forEach((function(e){return f.modifiersData[e.name]=Object.assign({},e.data)}));for(var r=0;r<f.orderedModifiers.length;r++)if(!0!==f.reset){var o=f.orderedModifiers[r],i=o.fn,a=o.options,s=void 0===a?{}:a,c=o.name;"function"==typeof i&&(f=i({state:f,options:s,name:c,instance:u})||f)}else f.reset=!1,r=-1}}},update:(i=function(){return new Promise((function(e){u.forceUpdate(),e(f)}))},function(){return s||(s=new Promise((function(e){Promise.resolve().then((function(){s=void 0,e(i())}))}))),s}),destroy:function(){l(),p=!0}};if(!Q(e,t))return u;function l(){c.forEach((function(e){return e()})),c=[]}return u.setOptions(r).then((function(e){!p&&r.onFirstUpdate&&r.onFirstUpdate(e)})),u}}var $={passive:!0};var ee={name:"eventListeners",enabled:!0,phase:"write",fn:function(){},effect:function(e){var n=e.state,r=e.instance,o=e.options,i=o.scroll,a=void 0===i||i,s=o.resize,f=void 0===s||s,c=t(n.elements.popper),p=[].concat(n.scrollParents.reference,n.scrollParents.popper);return a&&p.forEach((function(e){e.addEventListener("scroll",r.update,$)})),f&&c.addEventListener("resize",r.update,$),function(){a&&p.forEach((function(e){e.removeEventListener("scroll",r.update,$)})),f&&c.removeEventListener("resize",r.update,$)}},data:{}};var te={name:"popperOffsets",enabled:!0,phase:"read",fn:function(e){var t=e.state,n=e.name;t.modifiersData[n]=X({reference:t.rects.reference,element:t.rects.popper,strategy:"absolute",placement:t.placement})},data:{}},ne={top:"auto",right:"auto",bottom:"auto",left:"auto"};function re(e){var n,r=e.popper,o=e.popperRect,i=e.placement,a=e.variation,f=e.offsets,c=e.position,p=e.gpuAcceleration,u=e.adaptive,l=e.roundOffsets,h=e.isFixed,v=f.x,y=void 0===v?0:v,g=f.y,b=void 0===g?0:g,x="function"==typeof l?l({x:y,y:b}):{x:y,y:b};y=x.x,b=x.y;var w=f.hasOwnProperty("x"),O=f.hasOwnProperty("y"),j=P,M=D,k=window;if(u){var W=E(r),H="clientHeight",T="clientWidth";if(W===t(r)&&"static"!==m(W=d(r)).position&&"absolute"===c&&(H="scrollHeight",T="scrollWidth"),W=W,i===D||(i===P||i===L)&&a===B)M=A,b-=(h&&W===k&&k.visualViewport?k.visualViewport.height:W[H])-o.height,b*=p?1:-1;if(i===P||(i===D||i===A)&&a===B)j=L,y-=(h&&W===k&&k.visualViewport?k.visualViewport.width:W[T])-o.width,y*=p?1:-1}var R,S=Object.assign({position:c},u&&ne),V=!0===l?function(e,t){var n=e.x,r=e.y,o=t.devicePixelRatio||1;return{x:s(n*o)/o||0,y:s(r*o)/o||0}}({x:y,y:b},t(r)):{x:y,y:b};return y=V.x,b=V.y,p?Object.assign({},S,((R={})[M]=O?"0":"",R[j]=w?"0":"",R.transform=(k.devicePixelRatio||1)<=1?"translate("+y+"px, "+b+"px)":"translate3d("+y+"px, "+b+"px, 0)",R)):Object.assign({},S,((n={})[M]=O?b+"px":"",n[j]=w?y+"px":"",n.transform="",n))}var oe={name:"computeStyles",enabled:!0,phase:"beforeWrite",fn:function(e){var t=e.state,n=e.options,r=n.gpuAcceleration,o=void 0===r||r,i=n.adaptive,a=void 0===i||i,s=n.roundOffsets,f=void 0===s||s,c={placement:C(t.placement),variation:U(t.placement),popper:t.elements.popper,popperRect:t.rects.popper,gpuAcceleration:o,isFixed:"fixed"===t.options.strategy};null!=t.modifiersData.popperOffsets&&(t.styles.popper=Object.assign({},t.styles.popper,re(Object.assign({},c,{offsets:t.modifiersData.popperOffsets,position:t.options.strategy,adaptive:a,roundOffsets:f})))),null!=t.modifiersData.arrow&&(t.styles.arrow=Object.assign({},t.styles.arrow,re(Object.assign({},c,{offsets:t.modifiersData.arrow,position:"absolute",adaptive:!1,roundOffsets:f})))),t.attributes.popper=Object.assign({},t.attributes.popper,{"data-popper-placement":t.placement})},data:{}};var ie={name:"applyStyles",enabled:!0,phase:"write",fn:function(e){var t=e.state;Object.keys(t.elements).forEach((function(e){var n=t.styles[e]||{},o=t.attributes[e]||{},i=t.elements[e];r(i)&&l(i)&&(Object.assign(i.style,n),Object.keys(o).forEach((function(e){var t=o[e];!1===t?i.removeAttribute(e):i.setAttribute(e,!0===t?"":t)})))}))},effect:function(e){var t=e.state,n={popper:{position:t.options.strategy,left:"0",top:"0",margin:"0"},arrow:{position:"absolute"},reference:{}};return Object.assign(t.elements.popper.style,n.popper),t.styles=n,t.elements.arrow&&Object.assign(t.elements.arrow.style,n.arrow),function(){Object.keys(t.elements).forEach((function(e){var o=t.elements[e],i=t.attributes[e]||{},a=Object.keys(t.styles.hasOwnProperty(e)?t.styles[e]:n[e]).reduce((function(e,t){return e[t]="",e}),{});r(o)&&l(o)&&(Object.assign(o.style,a),Object.keys(i).forEach((function(e){o.removeAttribute(e)})))}))}},requires:["computeStyles"]};var ae={name:"offset",enabled:!0,phase:"main",requires:["popperOffsets"],fn:function(e){var t=e.state,n=e.options,r=e.name,o=n.offset,i=void 0===o?[0,0]:o,a=S.reduce((function(e,n){return e[n]=function(e,t,n){var r=C(e),o=[P,D].indexOf(r)>=0?-1:1,i="function"==typeof n?n(Object.assign({},t,{placement:e})):n,a=i[0],s=i[1];return a=a||0,s=(s||0)*o,[P,L].indexOf(r)>=0?{x:s,y:a}:{x:a,y:s}}(n,t.rects,i),e}),{}),s=a[t.placement],f=s.x,c=s.y;null!=t.modifiersData.popperOffsets&&(t.modifiersData.popperOffsets.x+=f,t.modifiersData.popperOffsets.y+=c),t.modifiersData[r]=a}},se={left:"right",right:"left",bottom:"top",top:"bottom"};function fe(e){return e.replace(/left|right|bottom|top/g,(function(e){return se[e]}))}var ce={start:"end",end:"start"};function pe(e){return e.replace(/start|end/g,(function(e){return ce[e]}))}function ue(e,t){void 0===t&&(t={});var n=t,r=n.placement,o=n.boundary,i=n.rootBoundary,a=n.padding,s=n.flipVariations,f=n.allowedAutoPlacements,c=void 0===f?S:f,p=U(r),u=p?s?R:R.filter((function(e){return U(e)===p})):k,l=u.filter((function(e){return c.indexOf(e)>=0}));0===l.length&&(l=u);var d=l.reduce((function(t,n){return t[n]=J(e,{placement:n,boundary:o,rootBoundary:i,padding:a})[C(n)],t}),{});return Object.keys(d).sort((function(e,t){return d[e]-d[t]}))}var le={name:"flip",enabled:!0,phase:"main",fn:function(e){var t=e.state,n=e.options,r=e.name;if(!t.modifiersData[r]._skip){for(var o=n.mainAxis,i=void 0===o||o,a=n.altAxis,s=void 0===a||a,f=n.fallbackPlacements,c=n.padding,p=n.boundary,u=n.rootBoundary,l=n.altBoundary,d=n.flipVariations,h=void 0===d||d,m=n.allowedAutoPlacements,v=t.options.placement,y=C(v),g=f||(y===v||!h?[fe(v)]:function(e){if(C(e)===M)return[];var t=fe(e);return[pe(e),t,pe(t)]}(v)),b=[v].concat(g).reduce((function(e,n){return e.concat(C(n)===M?ue(t,{placement:n,boundary:p,rootBoundary:u,padding:c,flipVariations:h,allowedAutoPlacements:m}):n)}),[]),x=t.rects.reference,w=t.rects.popper,O=new Map,j=!0,E=b[0],k=0;k<b.length;k++){var B=b[k],H=C(B),T=U(B)===W,R=[D,A].indexOf(H)>=0,S=R?"width":"height",V=J(t,{placement:B,boundary:p,rootBoundary:u,altBoundary:l,padding:c}),q=R?T?L:P:T?A:D;x[S]>w[S]&&(q=fe(q));var N=fe(q),I=[];if(i&&I.push(V[H]<=0),s&&I.push(V[q]<=0,V[N]<=0),I.every((function(e){return e}))){E=B,j=!1;break}O.set(B,I)}if(j)for(var _=function(e){var t=b.find((function(t){var n=O.get(t);if(n)return n.slice(0,e).every((function(e){return e}))}));if(t)return E=t,"break"},F=h?3:1;F>0;F--){if("break"===_(F))break}t.placement!==E&&(t.modifiersData[r]._skip=!0,t.placement=E,t.reset=!0)}},requiresIfExists:["offset"],data:{_skip:!1}};function de(e,t,n){return i(e,a(t,n))}var he={name:"preventOverflow",enabled:!0,phase:"main",fn:function(e){var t=e.state,n=e.options,r=e.name,o=n.mainAxis,s=void 0===o||o,f=n.altAxis,c=void 0!==f&&f,p=n.boundary,u=n.rootBoundary,l=n.altBoundary,d=n.padding,h=n.tether,m=void 0===h||h,v=n.tetherOffset,y=void 0===v?0:v,b=J(t,{boundary:p,rootBoundary:u,padding:d,altBoundary:l}),x=C(t.placement),w=U(t.placement),O=!w,j=z(x),M="x"===j?"y":"x",k=t.modifiersData.popperOffsets,B=t.rects.reference,H=t.rects.popper,T="function"==typeof y?y(Object.assign({},t.rects,{placement:t.placement})):y,R="number"==typeof T?{mainAxis:T,altAxis:T}:Object.assign({mainAxis:0,altAxis:0},T),S=t.modifiersData.offset?t.modifiersData.offset[t.placement]:null,V={x:0,y:0};if(k){if(s){var q,N="y"===j?D:P,I="y"===j?A:L,_="y"===j?"height":"width",F=k[j],X=F+b[N],Y=F-b[I],G=m?-H[_]/2:0,K=w===W?B[_]:H[_],Q=w===W?-H[_]:-B[_],Z=t.elements.arrow,$=m&&Z?g(Z):{width:0,height:0},ee=t.modifiersData["arrow#persistent"]?t.modifiersData["arrow#persistent"].padding:{top:0,right:0,bottom:0,left:0},te=ee[N],ne=ee[I],re=de(0,B[_],$[_]),oe=O?B[_]/2-G-re-te-R.mainAxis:K-re-te-R.mainAxis,ie=O?-B[_]/2+G+re+ne+R.mainAxis:Q+re+ne+R.mainAxis,ae=t.elements.arrow&&E(t.elements.arrow),se=ae?"y"===j?ae.clientTop||0:ae.clientLeft||0:0,fe=null!=(q=null==S?void 0:S[j])?q:0,ce=F+ie-fe,pe=de(m?a(X,F+oe-fe-se):X,F,m?i(Y,ce):Y);k[j]=pe,V[j]=pe-F}if(c){var ue,le="x"===j?D:P,he="x"===j?A:L,me=k[M],ve="y"===M?"height":"width",ye=me+b[le],ge=me-b[he],be=-1!==[D,P].indexOf(x),xe=null!=(ue=null==S?void 0:S[M])?ue:0,we=be?ye:me-B[ve]-H[ve]-xe+R.altAxis,Oe=be?me+B[ve]+H[ve]-xe-R.altAxis:ge,je=m&&be?function(e,t,n){var r=de(e,t,n);return r>n?n:r}(we,me,Oe):de(m?we:ye,me,m?Oe:ge);k[M]=je,V[M]=je-me}t.modifiersData[r]=V}},requiresIfExists:["offset"]};var me={name:"arrow",enabled:!0,phase:"main",fn:function(e){var t,n=e.state,r=e.name,o=e.options,i=n.elements.arrow,a=n.modifiersData.popperOffsets,s=C(n.placement),f=z(s),c=[P,L].indexOf(s)>=0?"height":"width";if(i&&a){var p=function(e,t){return Y("number"!=typeof(e="function"==typeof e?e(Object.assign({},t.rects,{placement:t.placement})):e)?e:G(e,k))}(o.padding,n),u=g(i),l="y"===f?D:P,d="y"===f?A:L,h=n.rects.reference[c]+n.rects.reference[f]-a[f]-n.rects.popper[c],m=a[f]-n.rects.reference[f],v=E(i),y=v?"y"===f?v.clientHeight||0:v.clientWidth||0:0,b=h/2-m/2,x=p[l],w=y-u[c]-p[d],O=y/2-u[c]/2+b,j=de(x,O,w),M=f;n.modifiersData[r]=((t={})[M]=j,t.centerOffset=j-O,t)}},effect:function(e){var t=e.state,n=e.options.element,r=void 0===n?"[data-popper-arrow]":n;null!=r&&("string"!=typeof r||(r=t.elements.popper.querySelector(r)))&&N(t.elements.popper,r)&&(t.elements.arrow=r)},requires:["popperOffsets"],requiresIfExists:["preventOverflow"]};function ve(e,t,n){return void 0===n&&(n={x:0,y:0}),{top:e.top-t.height-n.y,right:e.right-t.width+n.x,bottom:e.bottom-t.height+n.y,left:e.left-t.width-n.x}}function ye(e){return[D,L,A,P].some((function(t){return e[t]>=0}))}var ge={name:"hide",enabled:!0,phase:"main",requiresIfExists:["preventOverflow"],fn:function(e){var t=e.state,n=e.name,r=t.rects.reference,o=t.rects.popper,i=t.modifiersData.preventOverflow,a=J(t,{elementContext:"reference"}),s=J(t,{altBoundary:!0}),f=ve(a,r),c=ve(s,o,i),p=ye(f),u=ye(c);t.modifiersData[n]={referenceClippingOffsets:f,popperEscapeOffsets:c,isReferenceHidden:p,hasPopperEscaped:u},t.attributes.popper=Object.assign({},t.attributes.popper,{"data-popper-reference-hidden":p,"data-popper-escaped":u})}},be=Z({defaultModifiers:[ee,te,oe,ie]}),xe=[ee,te,oe,ie,ae,le,he,me,ge],we=Z({defaultModifiers:xe});e.applyStyles=ie,e.arrow=me,e.computeStyles=oe,e.createPopper=we,e.createPopperLite=be,e.defaultModifiers=xe,e.detectOverflow=J,e.eventListeners=ee,e.flip=le,e.hide=ge,e.offset=ae,e.popperGenerator=Z,e.popperOffsets=te,e.preventOverflow=he,Object.defineProperty(e,"__esModule",{value:!0})}));
```
