## Appendix for "Challenges and lessons learned from preliminary modeling of with-in herd transmission of highly pathogenic avian influenza H_5_N_1_ in dairy cattle": quarto.js

```
const sectionChanged = new CustomEvent("quarto-sectionChanged", {
  detail: {},
  bubbles: true,
  cancelable: false,
  composed: false,
});

const layoutMarginEls = () => {
  // Find any conflicting margin elements and add margins to the
  // top to prevent overlap
  const marginChildren = window.document.querySelectorAll(
    ".column-margin.column-container > *, .margin-caption, .aside"
  );

  let lastBottom = 0;
  for (const marginChild of marginChildren) {
    if (marginChild.offsetParent !== null) {
      // clear the top margin so we recompute it
      marginChild.style.marginTop = null;
      const top = marginChild.getBoundingClientRect().top + window.scrollY;
      if (top < lastBottom) {
        const marginChildStyle = window.getComputedStyle(marginChild);
        const marginBottom = parseFloat(marginChildStyle["marginBottom"]);
        const margin = lastBottom - top + marginBottom;
        marginChild.style.marginTop = `${margin}px`;
      }
      const styles = window.getComputedStyle(marginChild);
      const marginTop = parseFloat(styles["marginTop"]);
      lastBottom = top + marginChild.getBoundingClientRect().height + marginTop;
    }
  }
};

window.document.addEventListener("DOMContentLoaded", function (_event) {
  // Recompute the position of margin elements anytime the body size changes
  if (window.ResizeObserver) {
    const resizeObserver = new window.ResizeObserver(
      throttle(() => {
        layoutMarginEls();
        if (
          window.document.body.getBoundingClientRect().width < 990 &&
          isReaderMode()
        ) {
          quartoToggleReader();
        }
      }, 50)
    );
    resizeObserver.observe(window.document.body);
  }

  const tocEl = window.document.querySelector('nav.toc-active[role="doc-toc"]');
  const sidebarEl = window.document.getElementById("quarto-sidebar");
  const leftTocEl = window.document.getElementById("quarto-sidebar-toc-left");
  const marginSidebarEl = window.document.getElementById(
    "quarto-margin-sidebar"
  );
  // function to determine whether the element has a previous sibling that is active
  const prevSiblingIsActiveLink = (el) => {
    const sibling = el.previousElementSibling;
    if (sibling && sibling.tagName === "A") {
      return sibling.classList.contains("active");
    } else {
      return false;
    }
  };

  // fire slideEnter for bootstrap tab activations (for htmlwidget resize behavior)
  function fireSlideEnter(e) {
    const event = window.document.createEvent("Event");
    event.initEvent("slideenter", true, true);
    window.document.dispatchEvent(event);
  }
  const tabs = window.document.querySelectorAll('a[data-bs-toggle="tab"]');
  tabs.forEach((tab) => {
    tab.addEventListener("shown.bs.tab", fireSlideEnter);
  });

  // fire slideEnter for tabby tab activations (for htmlwidget resize behavior)
  document.addEventListener("tabby", fireSlideEnter, false);

  // Track scrolling and mark TOC links as active
  // get table of contents and sidebar (bail if we don't have at least one)
  const tocLinks = tocEl
    ? [...tocEl.querySelectorAll("a[data-scroll-target]")]
    : [];
  const makeActive = (link) => tocLinks[link].classList.add("active");
  const removeActive = (link) => tocLinks[link].classList.remove("active");
  const removeAllActive = () =>
    [...Array(tocLinks.length).keys()].forEach((link) => removeActive(link));

  // activate the anchor for a section associated with this TOC entry
  tocLinks.forEach((link) => {
    link.addEventListener("click", () => {
      if (link.href.indexOf("#") !== -1) {
        const anchor = link.href.split("#")[1];
        const heading = window.document.querySelector(
          `[data-anchor-id=${anchor}]`
        );
        if (heading) {
          // Add the class
          heading.classList.add("reveal-anchorjs-link");

          // function to show the anchor
          const handleMouseout = () => {
            heading.classList.remove("reveal-anchorjs-link");
            heading.removeEventListener("mouseout", handleMouseout);
          };

          // add a function to clear the anchor when the user mouses out of it
          heading.addEventListener("mouseout", handleMouseout);
        }
      }
    });
  });

  const sections = tocLinks.map((link) => {
    const target = link.getAttribute("data-scroll-target");
    if (target.startsWith("#")) {
      return window.document.getElementById(decodeURI(`${target.slice(1)}`));
    } else {
      return window.document.querySelector(decodeURI(`${target}`));
    }
  });

  const sectionMargin = 200;
  let currentActive = 0;
  // track whether we've initialized state the first time
  let init = false;

  const updateActiveLink = () => {
    // The index from bottom to top (e.g. reversed list)
    let sectionIndex = -1;
    if (
      window.innerHeight + window.pageYOffset >=
      window.document.body.offsetHeight
    ) {
      sectionIndex = 0;
    } else {
      sectionIndex = [...sections].reverse().findIndex((section) => {
        if (section) {
          return window.pageYOffset >= section.offsetTop - sectionMargin;
        } else {
          return false;
        }
      });
    }
    if (sectionIndex > -1) {
      const current = sections.length - sectionIndex - 1;
      if (current !== currentActive) {
        removeAllActive();
        currentActive = current;
        makeActive(current);
        if (init) {
          window.dispatchEvent(sectionChanged);
        }
        init = true;
      }
    }
  };

  const inHiddenRegion = (top, bottom, hiddenRegions) => {
    for (const region of hiddenRegions) {
      if (top <= region.bottom && bottom >= region.top) {
        return true;
      }
    }
    return false;
  };

  const categorySelector = "header.quarto-title-block .quarto-category";
  const activateCategories = (href) => {
    // Find any categories
    // Surround them with a link pointing back to:
    // #category=Authoring
    try {
      const categoryEls = window.document.querySelectorAll(categorySelector);
      for (const categoryEl of categoryEls) {
        const categoryText = categoryEl.textContent;
        if (categoryText) {
          const link = `${href}#category=${encodeURIComponent(categoryText)}`;
          const linkEl = window.document.createElement("a");
          linkEl.setAttribute("href", link);
          for (const child of categoryEl.childNodes) {
            linkEl.append(child);
          }
          categoryEl.appendChild(linkEl);
        }
      }
    } catch {
      // Ignore errors
    }
  };
  function hasTitleCategories() {
    return window.document.querySelector(categorySelector) !== null;
  }

  function offsetRelativeUrl(url) {
    const offset = getMeta("quarto:offset");
    return offset ? offset + url : url;
  }

  function offsetAbsoluteUrl(url) {
    const offset = getMeta("quarto:offset");
    const baseUrl = new URL(offset, window.location);

    const projRelativeUrl = url.replace(baseUrl, "");
    if (projRelativeUrl.startsWith("/")) {
      return projRelativeUrl;
    } else {
      return "/" + projRelativeUrl;
    }
  }

  // read a meta tag value
  function getMeta(metaName) {
    const metas = window.document.getElementsByTagName("meta");
    for (let i = 0; i < metas.length; i++) {
      if (metas[i].getAttribute("name") === metaName) {
        return metas[i].getAttribute("content");
      }
    }
    return "";
  }

  async function findAndActivateCategories() {
    const currentPagePath = offsetAbsoluteUrl(window.location.href);
    const response = await fetch(offsetRelativeUrl("listings.json"));
    if (response.status == 200) {
      return response.json().then(function (listingPaths) {
        const listingHrefs = [];
        for (const listingPath of listingPaths) {
          const pathWithoutLeadingSlash = listingPath.listing.substring(1);
          for (const item of listingPath.items) {
            if (
              item === currentPagePath ||
              item === currentPagePath + "index.html"
            ) {
              // Resolve this path against the offset to be sure
              // we already are using the correct path to the listing
              // (this adjusts the listing urls to be rooted against
              // whatever root the page is actually running against)
              const relative = offsetRelativeUrl(pathWithoutLeadingSlash);
              const baseUrl = window.location;
              const resolvedPath = new URL(relative, baseUrl);
              listingHrefs.push(resolvedPath.pathname);
              break;
            }
          }
        }

        // Look up the tree for a nearby linting and use that if we find one
        const nearestListing = findNearestParentListing(
          offsetAbsoluteUrl(window.location.pathname),
          listingHrefs
        );
        if (nearestListing) {
          activateCategories(nearestListing);
        } else {
          // See if the referrer is a listing page for this item
          const referredRelativePath = offsetAbsoluteUrl(document.referrer);
          const referrerListing = listingHrefs.find((listingHref) => {
            const isListingReferrer =
              listingHref === referredRelativePath ||
              listingHref === referredRelativePath + "index.html";
            return isListingReferrer;
          });

          if (referrerListing) {
            // Try to use the referrer if possible
            activateCategories(referrerListing);
          } else if (listingHrefs.length > 0) {
            // Otherwise, just fall back to the first listing
            activateCategories(listingHrefs[0]);
          }
        }
      });
    }
  }
  if (hasTitleCategories()) {
    findAndActivateCategories();
  }

  const findNearestParentListing = (href, listingHrefs) => {
    if (!href || !listingHrefs) {
      return undefined;
    }
    // Look up the tree for a nearby linting and use that if we find one
    const relativeParts = href.substring(1).split("/");
    while (relativeParts.length > 0) {
      const path = relativeParts.join("/");
      for (const listingHref of listingHrefs) {
        if (listingHref.startsWith(path)) {
          return listingHref;
        }
      }
      relativeParts.pop();
    }

    return undefined;
  };

  const manageSidebarVisiblity = (el, placeholderDescriptor) => {
    let isVisible = true;
    let elRect;

    return (hiddenRegions) => {
      if (el === null) {
        return;
      }

      // Find the last element of the TOC
      const lastChildEl = el.lastElementChild;

      if (lastChildEl) {
        // Converts the sidebar to a menu
        const convertToMenu = () => {
          for (const child of el.children) {
            child.style.opacity = 0;
            child.style.overflow = "hidden";
          }

          nexttick(() => {
            const toggleContainer = window.document.createElement("div");
            toggleContainer.style.width = "100%";
            toggleContainer.classList.add("zindex-over-content");
            toggleContainer.classList.add("quarto-sidebar-toggle");
            toggleContainer.classList.add("headroom-target"); // Marks this to be managed by headeroom
            toggleContainer.id = placeholderDescriptor.id;
            toggleContainer.style.position = "fixed";

            const toggleIcon = window.document.createElement("i");
            toggleIcon.classList.add("quarto-sidebar-toggle-icon");
            toggleIcon.classList.add("bi");
            toggleIcon.classList.add("bi-caret-down-fill");

            const toggleTitle = window.document.createElement("div");
            const titleEl = window.document.body.querySelector(
              placeholderDescriptor.titleSelector
            );
            if (titleEl) {
              toggleTitle.append(
                titleEl.textContent || titleEl.innerText,
                toggleIcon
              );
            }
            toggleTitle.classList.add("zindex-over-content");
            toggleTitle.classList.add("quarto-sidebar-toggle-title");
            toggleContainer.append(toggleTitle);

            const toggleContents = window.document.createElement("div");
            toggleContents.classList = el.classList;
            toggleContents.classList.add("zindex-over-content");
            toggleContents.classList.add("quarto-sidebar-toggle-contents");
            for (const child of el.children) {
              if (child.id === "toc-title") {
                continue;
              }

              const clone = child.cloneNode(true);
              clone.style.opacity = 1;
              clone.style.display = null;
              toggleContents.append(clone);
            }
            toggleContents.style.height = "0px";
            const positionToggle = () => {
              // position the element (top left of parent, same width as parent)
              if (!elRect) {
                elRect = el.getBoundingClientRect();
              }
              toggleContainer.style.left = `${elRect.left}px`;
              toggleContainer.style.top = `${elRect.top}px`;
              toggleContainer.style.width = `${elRect.width}px`;
            };
            positionToggle();

            toggleContainer.append(toggleContents);
            el.parentElement.prepend(toggleContainer);

            // Process clicks
            let tocShowing = false;
            // Allow the caller to control whether this is dismissed
            // when it is clicked (e.g. sidebar navigation supports
            // opening and closing the nav tree, so don't dismiss on click)
            const clickEl = placeholderDescriptor.dismissOnClick
              ? toggleContainer
              : toggleTitle;

            const closeToggle = () => {
              if (tocShowing) {
                toggleContainer.classList.remove("expanded");
                toggleContents.style.height = "0px";
                tocShowing = false;
              }
            };

            // Get rid of any expanded toggle if the user scrolls
            window.document.addEventListener(
              "scroll",
              throttle(() => {
                closeToggle();
              }, 50)
            );

            // Handle positioning of the toggle
            window.addEventListener(
              "resize",
              throttle(() => {
                elRect = undefined;
                positionToggle();
              }, 50)
            );

            window.addEventListener("quarto-hrChanged", () => {
              elRect = undefined;
            });

            // Process the click
            clickEl.onclick = () => {
              if (!tocShowing) {
                toggleContainer.classList.add("expanded");
                toggleContents.style.height = null;
                tocShowing = true;
              } else {
                closeToggle();
              }
            };
          });
        };

        // Converts a sidebar from a menu back to a sidebar
        const convertToSidebar = () => {
          for (const child of el.children) {
            child.style.opacity = 1;
            child.style.overflow = null;
          }

          const placeholderEl = window.document.getElementById(
            placeholderDescriptor.id
          );
          if (placeholderEl) {
            placeholderEl.remove();
          }

          el.classList.remove("rollup");
        };

        if (isReaderMode()) {
          convertToMenu();
          isVisible = false;
        } else {
          // Find the top and bottom o the element that is being managed
          const elTop = el.offsetTop;
          const elBottom =
            elTop + lastChildEl.offsetTop + lastChildEl.offsetHeight;

          if (!isVisible) {
            // If the element is current not visible reveal if there are
            // no conflicts with overlay regions
            if (!inHiddenRegion(elTop, elBottom, hiddenRegions)) {
              convertToSidebar();
              isVisible = true;
            }
          } else {
            // If the element is visible, hide it if it conflicts with overlay regions
            // and insert a placeholder toggle (or if we're in reader mode)
            if (inHiddenRegion(elTop, elBottom, hiddenRegions)) {
              convertToMenu();
              isVisible = false;
            }
          }
        }
      }
    };
  };

  const tabEls = document.querySelectorAll('a[data-bs-toggle="tab"]');
  for (const tabEl of tabEls) {
    const id = tabEl.getAttribute("data-bs-target");
    if (id) {
      const columnEl = document.querySelector(
        `${id} .column-margin, .tabset-margin-content`
      );
      if (columnEl)
        tabEl.addEventListener("shown.bs.tab", function (event) {
          const el = event.srcElement;
          if (el) {
            const visibleCls = `${el.id}-margin-content`;
            // walk up until we find a parent tabset
            let panelTabsetEl = el.parentElement;
            while (panelTabsetEl) {
              if (panelTabsetEl.classList.contains("panel-tabset")) {
                break;
              }
              panelTabsetEl = panelTabsetEl.parentElement;
            }

            if (panelTabsetEl) {
              const prevSib = panelTabsetEl.previousElementSibling;
              if (
                prevSib &&
                prevSib.classList.contains("tabset-margin-container")
              ) {
                const childNodes = prevSib.querySelectorAll(
                  ".tabset-margin-content"
                );
                for (const childEl of childNodes) {
                  if (childEl.classList.contains(visibleCls)) {
                    childEl.classList.remove("collapse");
                  } else {
                    childEl.classList.add("collapse");
                  }
                }
              }
            }
          }

          layoutMarginEls();
        });
    }
  }

  // Manage the visibility of the toc and the sidebar
  const marginScrollVisibility = manageSidebarVisiblity(marginSidebarEl, {
    id: "quarto-toc-toggle",
    titleSelector: "#toc-title",
    dismissOnClick: true,
  });
  const sidebarScrollVisiblity = manageSidebarVisiblity(sidebarEl, {
    id: "quarto-sidebarnav-toggle",
    titleSelector: ".title",
    dismissOnClick: false,
  });
  let tocLeftScrollVisibility;
  if (leftTocEl) {
    tocLeftScrollVisibility = manageSidebarVisiblity(leftTocEl, {
      id: "quarto-lefttoc-toggle",
      titleSelector: "#toc-title",
      dismissOnClick: true,
    });
  }

  // Find the first element that uses formatting in special columns
  const conflictingEls = window.document.body.querySelectorAll(
    '[class^="column-"], [class*=" column-"], aside, [class*="margin-caption"], [class*=" margin-caption"], [class*="margin-ref"], [class*=" margin-ref"]'
  );

  // Filter all the possibly conflicting elements into ones
  // the do conflict on the left or ride side
  const arrConflictingEls = Array.from(conflictingEls);
  const leftSideConflictEls = arrConflictingEls.filter((el) => {
    if (el.tagName === "ASIDE") {
      return false;
    }
    return Array.from(el.classList).find((className) => {
      return (
        className !== "column-body" &&
        className.startsWith("column-") &&
        !className.endsWith("right") &&
        !className.endsWith("container") &&
        className !== "column-margin"
      );
    });
  });
  const rightSideConflictEls = arrConflictingEls.filter((el) => {
    if (el.tagName === "ASIDE") {
      return true;
    }

    const hasMarginCaption = Array.from(el.classList).find((className) => {
      return className == "margin-caption";
    });
    if (hasMarginCaption) {
      return true;
    }

    return Array.from(el.classList).find((className) => {
      return (
        className !== "column-body" &&
        !className.endsWith("container") &&
        className.startsWith("column-") &&
        !className.endsWith("left")
      );
    });
  });

  const kOverlapPaddingSize = 10;
  function toRegions(els) {
    return els.map((el) => {
      const boundRect = el.getBoundingClientRect();
      const top =
        boundRect.top +
        document.documentElement.scrollTop -
        kOverlapPaddingSize;
      return {
        top,
        bottom: top + el.scrollHeight + 2 * kOverlapPaddingSize,
      };
    });
  }

  let hasObserved = false;
  const visibleItemObserver = (els) => {
    let visibleElements = [...els];
    const intersectionObserver = new IntersectionObserver(
      (entries, _observer) => {
        entries.forEach((entry) => {
          if (entry.isIntersecting) {
            if (visibleElements.indexOf(entry.target) === -1) {
              visibleElements.push(entry.target);
            }
          } else {
            visibleElements = visibleElements.filter((visibleEntry) => {
              return visibleEntry !== entry;
            });
          }
        });

        if (!hasObserved) {
          hideOverlappedSidebars();
        }
        hasObserved = true;
      },
      {}
    );
    els.forEach((el) => {
      intersectionObserver.observe(el);
    });

    return {
      getVisibleEntries: () => {
        return visibleElements;
      },
    };
  };

  const rightElementObserver = visibleItemObserver(rightSideConflictEls);
  const leftElementObserver = visibleItemObserver(leftSideConflictEls);

  const hideOverlappedSidebars = () => {
    marginScrollVisibility(toRegions(rightElementObserver.getVisibleEntries()));
    sidebarScrollVisiblity(toRegions(leftElementObserver.getVisibleEntries()));
    if (tocLeftScrollVisibility) {
      tocLeftScrollVisibility(
        toRegions(leftElementObserver.getVisibleEntries())
      );
    }
  };

  window.quartoToggleReader = () => {
    // Applies a slow class (or removes it)
    // to update the transition speed
    const slowTransition = (slow) => {
      const manageTransition = (id, slow) => {
        const el = document.getElementById(id);
        if (el) {
          if (slow) {
            el.classList.add("slow");
          } else {
            el.classList.remove("slow");
          }
        }
      };

      manageTransition("TOC", slow);
      manageTransition("quarto-sidebar", slow);
    };
    const readerMode = !isReaderMode();
    setReaderModeValue(readerMode);

    // If we're entering reader mode, slow the transition
    if (readerMode) {
      slowTransition(readerMode);
    }
    highlightReaderToggle(readerMode);
    hideOverlappedSidebars();

    // If we're exiting reader mode, restore the non-slow transition
    if (!readerMode) {
      slowTransition(!readerMode);
    }
  };

  const highlightReaderToggle = (readerMode) => {
    const els = document.querySelectorAll(".quarto-reader-toggle");
    if (els) {
      els.forEach((el) => {
        if (readerMode) {
          el.classList.add("reader");
        } else {
          el.classList.remove("reader");
        }
      });
    }
  };

  const setReaderModeValue = (val) => {
    if (window.location.protocol !== "file:") {
      window.localStorage.setItem("quarto-reader-mode", val);
    } else {
      localReaderMode = val;
    }
  };

  const isReaderMode = () => {
    if (window.location.protocol !== "file:") {
      return window.localStorage.getItem("quarto-reader-mode") === "true";
    } else {
      return localReaderMode;
    }
  };
  let localReaderMode = null;

  const tocOpenDepthStr = tocEl?.getAttribute("data-toc-expanded");
  const tocOpenDepth = tocOpenDepthStr ? Number(tocOpenDepthStr) : 1;

  // Walk the TOC and collapse/expand nodes
  // Nodes are expanded if:
  // - they are top level
  // - they have children that are 'active' links
  // - they are directly below an link that is 'active'
  const walk = (el, depth) => {
    // Tick depth when we enter a UL
    if (el.tagName === "UL") {
      depth = depth + 1;
    }

    // It this is active link
    let isActiveNode = false;
    if (el.tagName === "A" && el.classList.contains("active")) {
      isActiveNode = true;
    }

    // See if there is an active child to this element
    let hasActiveChild = false;
    for (child of el.children) {
      hasActiveChild = walk(child, depth) || hasActiveChild;
    }

    // Process the collapse state if this is an UL
    if (el.tagName === "UL") {
      if (tocOpenDepth === -1 && depth > 1) {
        el.classList.add("collapse");
      } else if (
        depth <= tocOpenDepth ||
        hasActiveChild ||
        prevSiblingIsActiveLink(el)
      ) {
        el.classList.remove("collapse");
      } else {
        el.classList.add("collapse");
      }

      // untick depth when we leave a UL
      depth = depth - 1;
    }
    return hasActiveChild || isActiveNode;
  };

  // walk the TOC and expand / collapse any items that should be shown

  if (tocEl) {
    walk(tocEl, 0);
    updateActiveLink();
  }

  // Throttle the scroll event and walk peridiocally
  window.document.addEventListener(
    "scroll",
    throttle(() => {
      if (tocEl) {
        updateActiveLink();
        walk(tocEl, 0);
      }
      if (!isReaderMode()) {
        hideOverlappedSidebars();
      }
    }, 5)
  );
  window.addEventListener(
    "resize",
    throttle(() => {
      if (!isReaderMode()) {
        hideOverlappedSidebars();
      }
    }, 10)
  );
  hideOverlappedSidebars();
  highlightReaderToggle(isReaderMode());
});

// grouped tabsets
window.addEventListener("pageshow", (_event) => {
  function getTabSettings() {
    const data = localStorage.getItem("quarto-persistent-tabsets-data");
    if (!data) {
      localStorage.setItem("quarto-persistent-tabsets-data", "{}");
      return {};
    }
    if (data) {
      return JSON.parse(data);
    }
  }

  function setTabSettings(data) {
    localStorage.setItem(
      "quarto-persistent-tabsets-data",
      JSON.stringify(data)
    );
  }

  function setTabState(groupName, groupValue) {
    const data = getTabSettings();
    data[groupName] = groupValue;
    setTabSettings(data);
  }

  function toggleTab(tab, active) {
    const tabPanelId = tab.getAttribute("aria-controls");
    const tabPanel = document.getElementById(tabPanelId);
    if (active) {
      tab.classList.add("active");
      tabPanel.classList.add("active");
    } else {
      tab.classList.remove("active");
      tabPanel.classList.remove("active");
    }
  }

  function toggleAll(selectedGroup, selectorsToSync) {
    for (const [thisGroup, tabs] of Object.entries(selectorsToSync)) {
      const active = selectedGroup === thisGroup;
      for (const tab of tabs) {
        toggleTab(tab, active);
      }
    }
  }

  function findSelectorsToSyncByLanguage() {
    const result = {};
    const tabs = Array.from(
      document.querySelectorAll(`div[data-group] a[id^='tabset-']`)
    );
    for (const item of tabs) {
      const div = item.parentElement.parentElement.parentElement;
      const group = div.getAttribute("data-group");
      if (!result[group]) {
        result[group] = {};
      }
      const selectorsToSync = result[group];
      const value = item.innerHTML;
      if (!selectorsToSync[value]) {
        selectorsToSync[value] = [];
      }
      selectorsToSync[value].push(item);
    }
    return result;
  }

  function setupSelectorSync() {
    const selectorsToSync = findSelectorsToSyncByLanguage();
    Object.entries(selectorsToSync).forEach(([group, tabSetsByValue]) => {
      Object.entries(tabSetsByValue).forEach(([value, items]) => {
        items.forEach((item) => {
          item.addEventListener("click", (_event) => {
            setTabState(group, value);
            toggleAll(value, selectorsToSync[group]);
          });
        });
      });
    });
    return selectorsToSync;
  }

  const selectorsToSync = setupSelectorSync();
  for (const [group, selectedName] of Object.entries(getTabSettings())) {
    const selectors = selectorsToSync[group];
    // it's possible that stale state gives us empty selections, so we explicitly check here.
    if (selectors) {
      toggleAll(selectedName, selectors);
    }
  }
});

function throttle(func, wait) {
  let waiting = false;
  return function () {
    if (!waiting) {
      func.apply(this, arguments);
      waiting = true;
      setTimeout(function () {
        waiting = false;
      }, wait);
    }
  };
}

function nexttick(func) {
  return setTimeout(func, 0);
}
```
