## Appendix for "Challenges and lessons learned from preliminary modeling of with-in herd transmission of highly pathogenic avian influenza H_5_N_1_ in dairy cattle": tippy.umd.min.js

```
!function(e,t){"object"==typeof exports&&"undefined"!=typeof module?module.exports=t(require("@popperjs/core")):"function"==typeof define&&define.amd?define(["@popperjs/core"],t):(e=e||self).tippy=t(e.Popper)}(this,(function(e){"use strict";var t={passive:!0,capture:!0},n=function(){return document.body};function r(e,t,n){if(Array.isArray(e)){var r=e[t];return null==r?Array.isArray(n)?n[t]:n:r}return e}function o(e,t){var n={}.toString.call(e);return 0===n.indexOf("[object")&&n.indexOf(t+"]")>-1}function i(e,t){return"function"==typeof e?e.apply(void 0,t):e}function a(e,t){return 0===t?e:function(r){clearTimeout(n),n=setTimeout((function(){e(r)}),t)};var n}function s(e,t){var n=Object.assign({},e);return t.forEach((function(e){delete n[e]})),n}function u(e){return[].concat(e)}function c(e,t){-1===e.indexOf(t)&&e.push(t)}function p(e){return e.split("-")[0]}function f(e){return[].slice.call(e)}function l(e){return Object.keys(e).reduce((function(t,n){return void 0!==e[n]&&(t[n]=e[n]),t}),{})}function d(){return document.createElement("div")}function v(e){return["Element","Fragment"].some((function(t){return o(e,t)}))}function m(e){return o(e,"MouseEvent")}function g(e){return!(!e||!e._tippy||e._tippy.reference!==e)}function h(e){return v(e)?[e]:function(e){return o(e,"NodeList")}(e)?f(e):Array.isArray(e)?e:f(document.querySelectorAll(e))}function b(e,t){e.forEach((function(e){e&&(e.style.transitionDuration=t+"ms")}))}function y(e,t){e.forEach((function(e){e&&e.setAttribute("data-state",t)}))}function w(e){var t,n=u(e)[0];return null!=n&&null!=(t=n.ownerDocument)&&t.body?n.ownerDocument:document}function E(e,t,n){var r=t+"EventListener";["transitionend","webkitTransitionEnd"].forEach((function(t){e[r](t,n)}))}function O(e,t){for(var n=t;n;){var r;if(e.contains(n))return!0;n=null==n.getRootNode||null==(r=n.getRootNode())?void 0:r.host}return!1}var x={isTouch:!1},C=0;function T(){x.isTouch||(x.isTouch=!0,window.performance&&document.addEventListener("mousemove",A))}function A(){var e=performance.now();e-C<20&&(x.isTouch=!1,document.removeEventListener("mousemove",A)),C=e}function L(){var e=document.activeElement;if(g(e)){var t=e._tippy;e.blur&&!t.state.isVisible&&e.blur()}}var D=!!("undefined"!=typeof window&&"undefined"!=typeof document)&&!!window.msCrypto,R=Object.assign({appendTo:n,aria:{content:"auto",expanded:"auto"},delay:0,duration:[300,250],getReferenceClientRect:null,hideOnClick:!0,ignoreAttributes:!1,interactive:!1,interactiveBorder:2,interactiveDebounce:0,moveTransition:"",offset:[0,10],onAfterUpdate:function(){},onBeforeUpdate:function(){},onCreate:function(){},onDestroy:function(){},onHidden:function(){},onHide:function(){},onMount:function(){},onShow:function(){},onShown:function(){},onTrigger:function(){},onUntrigger:function(){},onClickOutside:function(){},placement:"top",plugins:[],popperOptions:{},render:null,showOnCreate:!1,touch:!0,trigger:"mouseenter focus",triggerTarget:null},{animateFill:!1,followCursor:!1,inlinePositioning:!1,sticky:!1},{allowHTML:!1,animation:"fade",arrow:!0,content:"",inertia:!1,maxWidth:350,role:"tooltip",theme:"",zIndex:9999}),k=Object.keys(R);function P(e){var t=(e.plugins||[]).reduce((function(t,n){var r,o=n.name,i=n.defaultValue;o&&(t[o]=void 0!==e[o]?e[o]:null!=(r=R[o])?r:i);return t}),{});return Object.assign({},e,t)}function j(e,t){var n=Object.assign({},t,{content:i(t.content,[e])},t.ignoreAttributes?{}:function(e,t){return(t?Object.keys(P(Object.assign({},R,{plugins:t}))):k).reduce((function(t,n){var r=(e.getAttribute("data-tippy-"+n)||"").trim();if(!r)return t;if("content"===n)t[n]=r;else try{t[n]=JSON.parse(r)}catch(e){t[n]=r}return t}),{})}(e,t.plugins));return n.aria=Object.assign({},R.aria,n.aria),n.aria={expanded:"auto"===n.aria.expanded?t.interactive:n.aria.expanded,content:"auto"===n.aria.content?t.interactive?null:"describedby":n.aria.content},n}function M(e,t){e.innerHTML=t}function V(e){var t=d();return!0===e?t.className="tippy-arrow":(t.className="tippy-svg-arrow",v(e)?t.appendChild(e):M(t,e)),t}function I(e,t){v(t.content)?(M(e,""),e.appendChild(t.content)):"function"!=typeof t.content&&(t.allowHTML?M(e,t.content):e.textContent=t.content)}function S(e){var t=e.firstElementChild,n=f(t.children);return{box:t,content:n.find((function(e){return e.classList.contains("tippy-content")})),arrow:n.find((function(e){return e.classList.contains("tippy-arrow")||e.classList.contains("tippy-svg-arrow")})),backdrop:n.find((function(e){return e.classList.contains("tippy-backdrop")}))}}function N(e){var t=d(),n=d();n.className="tippy-box",n.setAttribute("data-state","hidden"),n.setAttribute("tabindex","-1");var r=d();function o(n,r){var o=S(t),i=o.box,a=o.content,s=o.arrow;r.theme?i.setAttribute("data-theme",r.theme):i.removeAttribute("data-theme"),"string"==typeof r.animation?i.setAttribute("data-animation",r.animation):i.removeAttribute("data-animation"),r.inertia?i.setAttribute("data-inertia",""):i.removeAttribute("data-inertia"),i.style.maxWidth="number"==typeof r.maxWidth?r.maxWidth+"px":r.maxWidth,r.role?i.setAttribute("role",r.role):i.removeAttribute("role"),n.content===r.content&&n.allowHTML===r.allowHTML||I(a,e.props),r.arrow?s?n.arrow!==r.arrow&&(i.removeChild(s),i.appendChild(V(r.arrow))):i.appendChild(V(r.arrow)):s&&i.removeChild(s)}return r.className="tippy-content",r.setAttribute("data-state","hidden"),I(r,e.props),t.appendChild(n),n.appendChild(r),o(e.props,e.props),{popper:t,onUpdate:o}}N.$$tippy=!0;var B=1,H=[],U=[];function _(o,s){var v,g,h,C,T,A,L,k,M=j(o,Object.assign({},R,P(l(s)))),V=!1,I=!1,N=!1,_=!1,F=[],W=a(we,M.interactiveDebounce),X=B++,Y=(k=M.plugins).filter((function(e,t){return k.indexOf(e)===t})),$={id:X,reference:o,popper:d(),popperInstance:null,props:M,state:{isEnabled:!0,isVisible:!1,isDestroyed:!1,isMounted:!1,isShown:!1},plugins:Y,clearDelayTimeouts:function(){clearTimeout(v),clearTimeout(g),cancelAnimationFrame(h)},setProps:function(e){if($.state.isDestroyed)return;ae("onBeforeUpdate",[$,e]),be();var t=$.props,n=j(o,Object.assign({},t,l(e),{ignoreAttributes:!0}));$.props=n,he(),t.interactiveDebounce!==n.interactiveDebounce&&(ce(),W=a(we,n.interactiveDebounce));t.triggerTarget&&!n.triggerTarget?u(t.triggerTarget).forEach((function(e){e.removeAttribute("aria-expanded")})):n.triggerTarget&&o.removeAttribute("aria-expanded");ue(),ie(),J&&J(t,n);$.popperInstance&&(Ce(),Ae().forEach((function(e){requestAnimationFrame(e._tippy.popperInstance.forceUpdate)})));ae("onAfterUpdate",[$,e])},setContent:function(e){$.setProps({content:e})},show:function(){var e=$.state.isVisible,t=$.state.isDestroyed,o=!$.state.isEnabled,a=x.isTouch&&!$.props.touch,s=r($.props.duration,0,R.duration);if(e||t||o||a)return;if(te().hasAttribute("disabled"))return;if(ae("onShow",[$],!1),!1===$.props.onShow($))return;$.state.isVisible=!0,ee()&&(z.style.visibility="visible");ie(),de(),$.state.isMounted||(z.style.transition="none");if(ee()){var u=re(),p=u.box,f=u.content;b([p,f],0)}A=function(){var e;if($.state.isVisible&&!_){if(_=!0,z.offsetHeight,z.style.transition=$.props.moveTransition,ee()&&$.props.animation){var t=re(),n=t.box,r=t.content;b([n,r],s),y([n,r],"visible")}se(),ue(),c(U,$),null==(e=$.popperInstance)||e.forceUpdate(),ae("onMount",[$]),$.props.animation&&ee()&&function(e,t){me(e,t)}(s,(function(){$.state.isShown=!0,ae("onShown",[$])}))}},function(){var e,t=$.props.appendTo,r=te();e=$.props.interactive&&t===n||"parent"===t?r.parentNode:i(t,[r]);e.contains(z)||e.appendChild(z);$.state.isMounted=!0,Ce()}()},hide:function(){var e=!$.state.isVisible,t=$.state.isDestroyed,n=!$.state.isEnabled,o=r($.props.duration,1,R.duration);if(e||t||n)return;if(ae("onHide",[$],!1),!1===$.props.onHide($))return;$.state.isVisible=!1,$.state.isShown=!1,_=!1,V=!1,ee()&&(z.style.visibility="hidden");if(ce(),ve(),ie(!0),ee()){var i=re(),a=i.box,s=i.content;$.props.animation&&(b([a,s],o),y([a,s],"hidden"))}se(),ue(),$.props.animation?ee()&&function(e,t){me(e,(function(){!$.state.isVisible&&z.parentNode&&z.parentNode.contains(z)&&t()}))}(o,$.unmount):$.unmount()},hideWithInteractivity:function(e){ne().addEventListener("mousemove",W),c(H,W),W(e)},enable:function(){$.state.isEnabled=!0},disable:function(){$.hide(),$.state.isEnabled=!1},unmount:function(){$.state.isVisible&&$.hide();if(!$.state.isMounted)return;Te(),Ae().forEach((function(e){e._tippy.unmount()})),z.parentNode&&z.parentNode.removeChild(z);U=U.filter((function(e){return e!==$})),$.state.isMounted=!1,ae("onHidden",[$])},destroy:function(){if($.state.isDestroyed)return;$.clearDelayTimeouts(),$.unmount(),be(),delete o._tippy,$.state.isDestroyed=!0,ae("onDestroy",[$])}};if(!M.render)return $;var q=M.render($),z=q.popper,J=q.onUpdate;z.setAttribute("data-tippy-root",""),z.id="tippy-"+$.id,$.popper=z,o._tippy=$,z._tippy=$;var G=Y.map((function(e){return e.fn($)})),K=o.hasAttribute("aria-expanded");return he(),ue(),ie(),ae("onCreate",[$]),M.showOnCreate&&Le(),z.addEventListener("mouseenter",(function(){$.props.interactive&&$.state.isVisible&&$.clearDelayTimeouts()})),z.addEventListener("mouseleave",(function(){$.props.interactive&&$.props.trigger.indexOf("mouseenter")>=0&&ne().addEventListener("mousemove",W)})),$;function Q(){var e=$.props.touch;return Array.isArray(e)?e:[e,0]}function Z(){return"hold"===Q()[0]}function ee(){var e;return!(null==(e=$.props.render)||!e.$$tippy)}function te(){return L||o}function ne(){var e=te().parentNode;return e?w(e):document}function re(){return S(z)}function oe(e){return $.state.isMounted&&!$.state.isVisible||x.isTouch||C&&"focus"===C.type?0:r($.props.delay,e?0:1,R.delay)}function ie(e){void 0===e&&(e=!1),z.style.pointerEvents=$.props.interactive&&!e?"":"none",z.style.zIndex=""+$.props.zIndex}function ae(e,t,n){var r;(void 0===n&&(n=!0),G.forEach((function(n){n[e]&&n[e].apply(n,t)})),n)&&(r=$.props)[e].apply(r,t)}function se(){var e=$.props.aria;if(e.content){var t="aria-"+e.content,n=z.id;u($.props.triggerTarget||o).forEach((function(e){var r=e.getAttribute(t);if($.state.isVisible)e.setAttribute(t,r?r+" "+n:n);else{var o=r&&r.replace(n,"").trim();o?e.setAttribute(t,o):e.removeAttribute(t)}}))}}function ue(){!K&&$.props.aria.expanded&&u($.props.triggerTarget||o).forEach((function(e){$.props.interactive?e.setAttribute("aria-expanded",$.state.isVisible&&e===te()?"true":"false"):e.removeAttribute("aria-expanded")}))}function ce(){ne().removeEventListener("mousemove",W),H=H.filter((function(e){return e!==W}))}function pe(e){if(!x.isTouch||!N&&"mousedown"!==e.type){var t=e.composedPath&&e.composedPath()[0]||e.target;if(!$.props.interactive||!O(z,t)){if(u($.props.triggerTarget||o).some((function(e){return O(e,t)}))){if(x.isTouch)return;if($.state.isVisible&&$.props.trigger.indexOf("click")>=0)return}else ae("onClickOutside",[$,e]);!0===$.props.hideOnClick&&($.clearDelayTimeouts(),$.hide(),I=!0,setTimeout((function(){I=!1})),$.state.isMounted||ve())}}}function fe(){N=!0}function le(){N=!1}function de(){var e=ne();e.addEventListener("mousedown",pe,!0),e.addEventListener("touchend",pe,t),e.addEventListener("touchstart",le,t),e.addEventListener("touchmove",fe,t)}function ve(){var e=ne();e.removeEventListener("mousedown",pe,!0),e.removeEventListener("touchend",pe,t),e.removeEventListener("touchstart",le,t),e.removeEventListener("touchmove",fe,t)}function me(e,t){var n=re().box;function r(e){e.target===n&&(E(n,"remove",r),t())}if(0===e)return t();E(n,"remove",T),E(n,"add",r),T=r}function ge(e,t,n){void 0===n&&(n=!1),u($.props.triggerTarget||o).forEach((function(r){r.addEventListener(e,t,n),F.push({node:r,eventType:e,handler:t,options:n})}))}function he(){var e;Z()&&(ge("touchstart",ye,{passive:!0}),ge("touchend",Ee,{passive:!0})),(e=$.props.trigger,e.split(/\s+/).filter(Boolean)).forEach((function(e){if("manual"!==e)switch(ge(e,ye),e){case"mouseenter":ge("mouseleave",Ee);break;case"focus":ge(D?"focusout":"blur",Oe);break;case"focusin":ge("focusout",Oe)}}))}function be(){F.forEach((function(e){var t=e.node,n=e.eventType,r=e.handler,o=e.options;t.removeEventListener(n,r,o)})),F=[]}function ye(e){var t,n=!1;if($.state.isEnabled&&!xe(e)&&!I){var r="focus"===(null==(t=C)?void 0:t.type);C=e,L=e.currentTarget,ue(),!$.state.isVisible&&m(e)&&H.forEach((function(t){return t(e)})),"click"===e.type&&($.props.trigger.indexOf("mouseenter")<0||V)&&!1!==$.props.hideOnClick&&$.state.isVisible?n=!0:Le(e),"click"===e.type&&(V=!n),n&&!r&&De(e)}}function we(e){var t=e.target,n=te().contains(t)||z.contains(t);"mousemove"===e.type&&n||function(e,t){var n=t.clientX,r=t.clientY;return e.every((function(e){var t=e.popperRect,o=e.popperState,i=e.props.interactiveBorder,a=p(o.placement),s=o.modifiersData.offset;if(!s)return!0;var u="bottom"===a?s.top.y:0,c="top"===a?s.bottom.y:0,f="right"===a?s.left.x:0,l="left"===a?s.right.x:0,d=t.top-r+u>i,v=r-t.bottom-c>i,m=t.left-n+f>i,g=n-t.right-l>i;return d||v||m||g}))}(Ae().concat(z).map((function(e){var t,n=null==(t=e._tippy.popperInstance)?void 0:t.state;return n?{popperRect:e.getBoundingClientRect(),popperState:n,props:M}:null})).filter(Boolean),e)&&(ce(),De(e))}function Ee(e){xe(e)||$.props.trigger.indexOf("click")>=0&&V||($.props.interactive?$.hideWithInteractivity(e):De(e))}function Oe(e){$.props.trigger.indexOf("focusin")<0&&e.target!==te()||$.props.interactive&&e.relatedTarget&&z.contains(e.relatedTarget)||De(e)}function xe(e){return!!x.isTouch&&Z()!==e.type.indexOf("touch")>=0}function Ce(){Te();var t=$.props,n=t.popperOptions,r=t.placement,i=t.offset,a=t.getReferenceClientRect,s=t.moveTransition,u=ee()?S(z).arrow:null,c=a?{getBoundingClientRect:a,contextElement:a.contextElement||te()}:o,p=[{name:"offset",options:{offset:i}},{name:"preventOverflow",options:{padding:{top:2,bottom:2,left:5,right:5}}},{name:"flip",options:{padding:5}},{name:"computeStyles",options:{adaptive:!s}},{name:"$$tippy",enabled:!0,phase:"beforeWrite",requires:["computeStyles"],fn:function(e){var t=e.state;if(ee()){var n=re().box;["placement","reference-hidden","escaped"].forEach((function(e){"placement"===e?n.setAttribute("data-placement",t.placement):t.attributes.popper["data-popper-"+e]?n.setAttribute("data-"+e,""):n.removeAttribute("data-"+e)})),t.attributes.popper={}}}}];ee()&&u&&p.push({name:"arrow",options:{element:u,padding:3}}),p.push.apply(p,(null==n?void 0:n.modifiers)||[]),$.popperInstance=e.createPopper(c,z,Object.assign({},n,{placement:r,onFirstUpdate:A,modifiers:p}))}function Te(){$.popperInstance&&($.popperInstance.destroy(),$.popperInstance=null)}function Ae(){return f(z.querySelectorAll("[data-tippy-root]"))}function Le(e){$.clearDelayTimeouts(),e&&ae("onTrigger",[$,e]),de();var t=oe(!0),n=Q(),r=n[0],o=n[1];x.isTouch&&"hold"===r&&o&&(t=o),t?v=setTimeout((function(){$.show()}),t):$.show()}function De(e){if($.clearDelayTimeouts(),ae("onUntrigger",[$,e]),$.state.isVisible){if(!($.props.trigger.indexOf("mouseenter")>=0&&$.props.trigger.indexOf("click")>=0&&["mouseleave","mousemove"].indexOf(e.type)>=0&&V)){var t=oe(!1);t?g=setTimeout((function(){$.state.isVisible&&$.hide()}),t):h=requestAnimationFrame((function(){$.hide()}))}}else ve()}}function F(e,n){void 0===n&&(n={});var r=R.plugins.concat(n.plugins||[]);document.addEventListener("touchstart",T,t),window.addEventListener("blur",L);var o=Object.assign({},n,{plugins:r}),i=h(e).reduce((function(e,t){var n=t&&_(t,o);return n&&e.push(n),e}),[]);return v(e)?i[0]:i}F.defaultProps=R,F.setDefaultProps=function(e){Object.keys(e).forEach((function(t){R[t]=e[t]}))},F.currentInput=x;var W=Object.assign({},e.applyStyles,{effect:function(e){var t=e.state,n={popper:{position:t.options.strategy,left:"0",top:"0",margin:"0"},arrow:{position:"absolute"},reference:{}};Object.assign(t.elements.popper.style,n.popper),t.styles=n,t.elements.arrow&&Object.assign(t.elements.arrow.style,n.arrow)}}),X={mouseover:"mouseenter",focusin:"focus",click:"click"};var Y={name:"animateFill",defaultValue:!1,fn:function(e){var t;if(null==(t=e.props.render)||!t.$$tippy)return{};var n=S(e.popper),r=n.box,o=n.content,i=e.props.animateFill?function(){var e=d();return e.className="tippy-backdrop",y([e],"hidden"),e}():null;return{onCreate:function(){i&&(r.insertBefore(i,r.firstElementChild),r.setAttribute("data-animatefill",""),r.style.overflow="hidden",e.setProps({arrow:!1,animation:"shift-away"}))},onMount:function(){if(i){var e=r.style.transitionDuration,t=Number(e.replace("ms",""));o.style.transitionDelay=Math.round(t/10)+"ms",i.style.transitionDuration=e,y([i],"visible")}},onShow:function(){i&&(i.style.transitionDuration="0ms")},onHide:function(){i&&y([i],"hidden")}}}};var $={clientX:0,clientY:0},q=[];function z(e){var t=e.clientX,n=e.clientY;$={clientX:t,clientY:n}}var J={name:"followCursor",defaultValue:!1,fn:function(e){var t=e.reference,n=w(e.props.triggerTarget||t),r=!1,o=!1,i=!0,a=e.props;function s(){return"initial"===e.props.followCursor&&e.state.isVisible}function u(){n.addEventListener("mousemove",f)}function c(){n.removeEventListener("mousemove",f)}function p(){r=!0,e.setProps({getReferenceClientRect:null}),r=!1}function f(n){var r=!n.target||t.contains(n.target),o=e.props.followCursor,i=n.clientX,a=n.clientY,s=t.getBoundingClientRect(),u=i-s.left,c=a-s.top;!r&&e.props.interactive||e.setProps({getReferenceClientRect:function(){var e=t.getBoundingClientRect(),n=i,r=a;"initial"===o&&(n=e.left+u,r=e.top+c);var s="horizontal"===o?e.top:r,p="vertical"===o?e.right:n,f="horizontal"===o?e.bottom:r,l="vertical"===o?e.left:n;return{width:p-l,height:f-s,top:s,right:p,bottom:f,left:l}}})}function l(){e.props.followCursor&&(q.push({instance:e,doc:n}),function(e){e.addEventListener("mousemove",z)}(n))}function d(){0===(q=q.filter((function(t){return t.instance!==e}))).filter((function(e){return e.doc===n})).length&&function(e){e.removeEventListener("mousemove",z)}(n)}return{onCreate:l,onDestroy:d,onBeforeUpdate:function(){a=e.props},onAfterUpdate:function(t,n){var i=n.followCursor;r||void 0!==i&&a.followCursor!==i&&(d(),i?(l(),!e.state.isMounted||o||s()||u()):(c(),p()))},onMount:function(){e.props.followCursor&&!o&&(i&&(f($),i=!1),s()||u())},onTrigger:function(e,t){m(t)&&($={clientX:t.clientX,clientY:t.clientY}),o="focus"===t.type},onHidden:function(){e.props.followCursor&&(p(),c(),i=!0)}}}};var G={name:"inlinePositioning",defaultValue:!1,fn:function(e){var t,n=e.reference;var r=-1,o=!1,i=[],a={name:"tippyInlinePositioning",enabled:!0,phase:"afterWrite",fn:function(o){var a=o.state;e.props.inlinePositioning&&(-1!==i.indexOf(a.placement)&&(i=[]),t!==a.placement&&-1===i.indexOf(a.placement)&&(i.push(a.placement),e.setProps({getReferenceClientRect:function(){return function(e){return function(e,t,n,r){if(n.length<2||null===e)return t;if(2===n.length&&r>=0&&n[0].left>n[1].right)return n[r]||t;switch(e){case"top":case"bottom":var o=n[0],i=n[n.length-1],a="top"===e,s=o.top,u=i.bottom,c=a?o.left:i.left,p=a?o.right:i.right;return{top:s,bottom:u,left:c,right:p,width:p-c,height:u-s};case"left":case"right":var f=Math.min.apply(Math,n.map((function(e){return e.left}))),l=Math.max.apply(Math,n.map((function(e){return e.right}))),d=n.filter((function(t){return"left"===e?t.left===f:t.right===l})),v=d[0].top,m=d[d.length-1].bottom;return{top:v,bottom:m,left:f,right:l,width:l-f,height:m-v};default:return t}}(p(e),n.getBoundingClientRect(),f(n.getClientRects()),r)}(a.placement)}})),t=a.placement)}};function s(){var t;o||(t=function(e,t){var n;return{popperOptions:Object.assign({},e.popperOptions,{modifiers:[].concat(((null==(n=e.popperOptions)?void 0:n.modifiers)||[]).filter((function(e){return e.name!==t.name})),[t])})}}(e.props,a),o=!0,e.setProps(t),o=!1)}return{onCreate:s,onAfterUpdate:s,onTrigger:function(t,n){if(m(n)){var o=f(e.reference.getClientRects()),i=o.find((function(e){return e.left-2<=n.clientX&&e.right+2>=n.clientX&&e.top-2<=n.clientY&&e.bottom+2>=n.clientY})),a=o.indexOf(i);r=a>-1?a:r}},onHidden:function(){r=-1}}}};var K={name:"sticky",defaultValue:!1,fn:function(e){var t=e.reference,n=e.popper;function r(t){return!0===e.props.sticky||e.props.sticky===t}var o=null,i=null;function a(){var s=r("reference")?(e.popperInstance?e.popperInstance.state.elements.reference:t).getBoundingClientRect():null,u=r("popper")?n.getBoundingClientRect():null;(s&&Q(o,s)||u&&Q(i,u))&&e.popperInstance&&e.popperInstance.update(),o=s,i=u,e.state.isMounted&&requestAnimationFrame(a)}return{onMount:function(){e.props.sticky&&a()}}}};function Q(e,t){return!e||!t||(e.top!==t.top||e.right!==t.right||e.bottom!==t.bottom||e.left!==t.left)}return F.setDefaultProps({plugins:[Y,J,G,K],render:N}),F.createSingleton=function(e,t){var n;void 0===t&&(t={});var r,o=e,i=[],a=[],c=t.overrides,p=[],f=!1;function l(){a=o.map((function(e){return u(e.props.triggerTarget||e.reference)})).reduce((function(e,t){return e.concat(t)}),[])}function v(){i=o.map((function(e){return e.reference}))}function m(e){o.forEach((function(t){e?t.enable():t.disable()}))}function g(e){return o.map((function(t){var n=t.setProps;return t.setProps=function(o){n(o),t.reference===r&&e.setProps(o)},function(){t.setProps=n}}))}function h(e,t){var n=a.indexOf(t);if(t!==r){r=t;var s=(c||[]).concat("content").reduce((function(e,t){return e[t]=o[n].props[t],e}),{});e.setProps(Object.assign({},s,{getReferenceClientRect:"function"==typeof s.getReferenceClientRect?s.getReferenceClientRect:function(){var e;return null==(e=i[n])?void 0:e.getBoundingClientRect()}}))}}m(!1),v(),l();var b={fn:function(){return{onDestroy:function(){m(!0)},onHidden:function(){r=null},onClickOutside:function(e){e.props.showOnCreate&&!f&&(f=!0,r=null)},onShow:function(e){e.props.showOnCreate&&!f&&(f=!0,h(e,i[0]))},onTrigger:function(e,t){h(e,t.currentTarget)}}}},y=F(d(),Object.assign({},s(t,["overrides"]),{plugins:[b].concat(t.plugins||[]),triggerTarget:a,popperOptions:Object.assign({},t.popperOptions,{modifiers:[].concat((null==(n=t.popperOptions)?void 0:n.modifiers)||[],[W])})})),w=y.show;y.show=function(e){if(w(),!r&&null==e)return h(y,i[0]);if(!r||null!=e){if("number"==typeof e)return i[e]&&h(y,i[e]);if(o.indexOf(e)>=0){var t=e.reference;return h(y,t)}return i.indexOf(e)>=0?h(y,e):void 0}},y.showNext=function(){var e=i[0];if(!r)return y.show(0);var t=i.indexOf(r);y.show(i[t+1]||e)},y.showPrevious=function(){var e=i[i.length-1];if(!r)return y.show(e);var t=i.indexOf(r),n=i[t-1]||e;y.show(n)};var E=y.setProps;return y.setProps=function(e){c=e.overrides||c,E(e)},y.setInstances=function(e){m(!0),p.forEach((function(e){return e()})),o=e,m(!1),v(),l(),p=g(y),y.setProps({triggerTarget:a})},p=g(y),y},F.delegate=function(e,n){var r=[],o=[],i=!1,a=n.target,c=s(n,["target"]),p=Object.assign({},c,{trigger:"manual",touch:!1}),f=Object.assign({touch:R.touch},c,{showOnCreate:!0}),l=F(e,p);function d(e){if(e.target&&!i){var t=e.target.closest(a);if(t){var r=t.getAttribute("data-tippy-trigger")||n.trigger||R.trigger;if(!t._tippy&&!("touchstart"===e.type&&"boolean"==typeof f.touch||"touchstart"!==e.type&&r.indexOf(X[e.type])<0)){var s=F(t,f);s&&(o=o.concat(s))}}}}function v(e,t,n,o){void 0===o&&(o=!1),e.addEventListener(t,n,o),r.push({node:e,eventType:t,handler:n,options:o})}return u(l).forEach((function(e){var n=e.destroy,a=e.enable,s=e.disable;e.destroy=function(e){void 0===e&&(e=!0),e&&o.forEach((function(e){e.destroy()})),o=[],r.forEach((function(e){var t=e.node,n=e.eventType,r=e.handler,o=e.options;t.removeEventListener(n,r,o)})),r=[],n()},e.enable=function(){a(),o.forEach((function(e){return e.enable()})),i=!1},e.disable=function(){s(),o.forEach((function(e){return e.disable()})),i=!0},function(e){var n=e.reference;v(n,"touchstart",d,t),v(n,"mouseover",d),v(n,"focusin",d),v(n,"click",d)}(e)})),l},F.hideAll=function(e){var t=void 0===e?{}:e,n=t.exclude,r=t.duration;U.forEach((function(e){var t=!1;if(n&&(t=g(n)?e.reference===n:e.popper===n.popper),!t){var o=e.props.duration;e.setProps({duration:r}),e.hide(),e.state.isDestroyed||e.setProps({duration:o})}}))},F.roundArrow='<svg width="16" height="6" xmlns="http://www.w3.org/2000/svg"><path d="M0 6s1.796-.013 4.67-3.615C5.851.9 6.93.006 8 0c1.07-.006 2.148.887 3.343 2.385C14.233 6.005 16 6 16 6H0z"></svg>',F}));
```
