## Appendix for "Challenges and lessons learned from preliminary modeling of with-in herd transmission of highly pathogenic avian influenza H_5_N_1_ in dairy cattle": AppendixCode2.R

```
library(tidyverse)
library(dde)
library(odin)
library(data.table)

model_sir <- odin({
  steps_per_day <- user(1)
  dt <- 1 / steps_per_day
  initial(time) <- 0
  update(time) <- (step + 1) * dt

  ## Compartments 
  update(S[]) <- S[i] - n_SI[i]
  update(I[]) <- I[i] + n_SI[i] - n_IR[i]
  update(R[]) <- R[i] + n_IR[i]

  ## Transition probs
  p_SI[] <- 1 - exp(-beta * I[i] / N[i]) # S to I
  p_IR <- 1 - exp(-gamma) # I to B1


  n_SI[] <- rbinom(S[i], p_SI[i] * dt)
  n_IR[] <- rbinom(I[i], p_IR * dt)
  

  ## Total population size
  N[] <- S[i] + I[i] + R[i]

  initial(S[]) <- S_ini
  initial(I[]) <- I_ini
  initial(R[]) <- 0
  
  ## Convert some ßvalues into incidence
  initial(S_inc[]) <- 0
  update(S_inc[]) <- if (step %% steps_per_day == 0) n_SI[i] else S_inc[i] + n_SI[i]
  
  ## User inputs
  S_ini <- user(499)
  I_ini <- user(1)
  beta <- user(0.83)
  gamma <- user(0.5)

  nsim <- user(500)
  dim(N) <- nsim
  dim(S) <- nsim
  dim(I) <- nsim
  dim(R) <- nsim
  dim(S_inc) <- nsim
  dim(p_SI) <- nsim
  dim(n_SI) <- nsim
  dim(n_IR) <- nsim


})


x <- model_sir$new(beta = 1.67/(1/2), gamma = 1/2)

run_scenarion <- function(r0_estimate = 1.67, fever_duration = 2){
  gamma_in <- 1 / fever_duration
  beta_in <- r0_estimate * gamma_in
  x <- model_sir$new(beta = beta_in, gamma = gamma_in)
  sim <- tibble::as_tibble(x$run(0:180))
  sim_long <- data.table::setDT(sim)
  sim_long <- melt(sim_long, id.vars = c("step", "time"))[
    , iter := stringr::str_extract(variable, "\\d+")][
    , compartment := gsub("[^a-zA-Z_]", "", variable)
    ][
    ,variable := NULL
    ]

  sim_narrow <- dcast(sim_long, step + time + iter ~ compartment, value.var = "value")[order(iter, time)]

  sim_narrow[ , prop_infected := I / 500]

  out_scenarios <- sim_narrow[, .SD[which.max(prop_infected)], by = iter, .SDcols = c("time", "prop_infected")] |>
    setNames(c("iter", "epi_peak_day", "prop_infected_peak"))

  epi_duration <- sim_narrow[, .SD[which.max(R)], by = iter, .SDcols = c("time", "R")][

  ] |>
  setNames(c("iter", "epi_duration", "R"))
  epi_duration[ , prop_max_infected := R / 500]
  out_scenarios[ , beta := beta_in]
  out_scenarios[ , gamma := gamma_in]
  out_scenarios[ , duration_infectious := fever_duration]
  out_scenarios[ , r0_est := r0_estimate]
  out_scenarios <- merge(out_scenarios, epi_duration, by = "iter")
  
  #sim$prop_infected <- sim$I / 500
  return(out_scenarios)
}

sim_grid <- tidyr::expand_grid(r = seq(1, 6, length.out = 100), gamma = seq(1, 7, length.out = 100))

z <- run_scenarion()

if (!file.exists(here::here("src", "simulation-stochastic-parameter-search.rds"))) {
  simulations <- map2(sim_grid$r, sim_grid$gamma, ~run_scenarion(.x, .y))
  saveRDS(simulations, here::here("src", "simulation-stochastic-parameter-search.rds"))
} else {
  simulations <- readRDS(here::here("src", "simulation-stochastic-parameter-search.rds"))
}


sim_fitting_params <- bind_rows(simulations) |>
  filter((epi_duration >= 20 & epi_duration <= 30) &
           prop_infected_peak >= .2 & prop_infected_peak <= 0.4 &
           epi_peak_day >= 7 & epi_peak_day <= 17) |>
  arrange(beta)

sim_fitting_params <- bind_rows(simulations) |>
  filter(prop_max_infected > 5/500) |>
  filter((epi_duration >= 20 & epi_duration <= 30) &
           #prop_infected_peak >= .2 & prop_infected_peak <= 0.4 &
           prop_max_infected >= .82 & prop_max_infected < .91,
           epi_peak_day >= 7 & epi_peak_day <= 17) |>
  arrange(beta)

out_fever <- round(quantile(sim_fitting_params$duration_infectious), 2)
out_r0 <- round(quantile(sim_fitting_params$r0_est), 2)

make_plot <- function(){
  par(mfrow = c(1,3))
hist(sim_fitting_params$r0_est, breaks = 30,
     xlab = expression("Estimated "~R[0]),
     main = ""
     #main = "Estimated R0 from stochastic simulations"
     )
#mtext(side = 3, sprintf("Estimated: %s [range: %s, %s]", out_r0[3], out_r0[1], out_r0[5]))
mtext(side = 3, adj = 0, "A", line = 1)

plot(cbind(sim_fitting_params$beta, sim_fitting_params$duration_infectious),
     xlab = expression(beta), ylab = "duration of infection (Days)",
     main = ""
     #main = "Exploration of parameters"
     )
mtext(side = 3, adj = 0, "B", line = 1)

hist(sim_fitting_params$duration_infectious, breaks = 30,
     xlab = "Estimated Duration of Infectious (days)",
     main = "",
     #main = "Estimated Infectious from stochastic simulations"
     )
#mtext(side = 3, sprintf("Estimated: %s [range: %s, %s]", out_fever[3], out_fever[1], out_fever[5]))
mtext(side = 3, adj = 0, "C", line = 1)
}

cairo_pdf(here::here("figures", "stochastic-sir-parameter-estimate.pdf"),
    height = 8.5, width = 11)
    make_plot()
dev.off()

png(here::here("figures", "stochastic-sir-parameter-estimate.png"),
    height = 8.5, width = 11, units = "in", res = 300)
    make_plot()
dev.off()


png(here::here("figures", "stochastic-sir-parameter-kde-estimates.png"),
    height = 8.5, width = 11, units = "in", res = 300)
par(mfrow=c(1,1))
filled.contour(MASS::kde2d(sim_fitting_params$duration_infectious, 
        sim_fitting_params$r0_est), ylab = expression(R[0]),
        xlab = "Duration of infectious (days)", plot.axes = {
          axis(1)
          axis(2)
contour(MASS::kde2d(sim_fitting_params$duration_infectious,
        sim_fitting_params$r0_est), add = TRUE)
        })
dev.off()
hist(z$prop_max_infected, breaks = 30)

x$run(0:30)


joint_param_space <- MASS::kde2d(sim_fitting_params$duration_infectious, 
        sim_fitting_params$r0_est)


o <- which(joint_param_space$z == max(joint_param_space$z), arr.ind = TRUE)
# Summarizing the values ------------------------------------------
cat("Fever duration: ", joint_param_space$x[o[1]])
cat("R0: ", joint_param_space$y[o[2]])

dat_to_bootstrap <- cbind(infectious = sim_fitting_params$duration_infectious,
                          r0 = sim_fitting_params$r0_est)
# Bootstrap helper functions ------------------------------------------
estimation_function <- function(x){
  joint_param_space <- MASS::kde2d(x[ ,1], x[ ,2])
  o <- which(joint_param_space$z == max(joint_param_space$z), arr.ind = TRUE)
  data.frame(infection = joint_param_space$x[o[1]], r0 = joint_param_space$y[o[2]])
}

sample_data <- function(x){
  x[sample(nrow(x), replace = TRUE),]
}
# Run the parametric boostrap ------------------------------------------
library(furrr)
# Do this over multiple cores because it is intensive
plan(multisession)
out_boot <- future_map_dfr(1:500, ~estimation_function(sample_data(dat_to_bootstrap)))
plan(sequential)
# Examine the results ------------------------------------------
lapply(out_boot, quantile, probs = c(.025, .975))
```
