## Appendix for "Challenges and lessons learned from preliminary modeling of with-in herd transmission of highly pathogenic avian influenza H_5_N_1_ in dairy cattle": Technical Appendix.docx

$\frac{d\mathbf{S}}{dt}=\mu\mathbf{N}-\beta\mathbf{IS}-\mu\mathbf{S}$	(1.1)

$\frac{d\mathbf{I}}{dt}=\beta\mathbf{IS-}\gamma\mathbf{I}-\mu\mathbf{I}$	(1.2)

$\frac{d\mathbf{B}}{dt}=\gamma\left( 1-\varphi\right)\mathbf{I}-\alpha\mathbf{B}-\mu\mathbf{B}$	(1.3)

$\frac{d\mathbf{R}}{dt}=\alpha\left( 1-\rho\right)\mathbf{B}-\mu\mathbf{R}$	(1.4)

$\frac{d\mathbf{D}}{dt}=\varphi\gamma\mathbf{I}$	(1.5)

$\frac{d\mathbf{C}}{dt}=\rho\alpha\mathbf{B}$	(1.6)
