## Supplementary figures and images for "Challenges and lessons learned from preliminary modeling of with-in herd transmission of highly pathogenic avian influenza H_5_N_1_ in dairy cattle"

### Appendix Figure 1.tiff

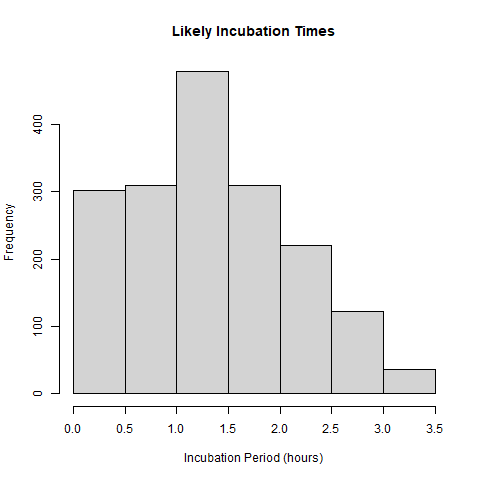

### Appendix Figure 2.tif

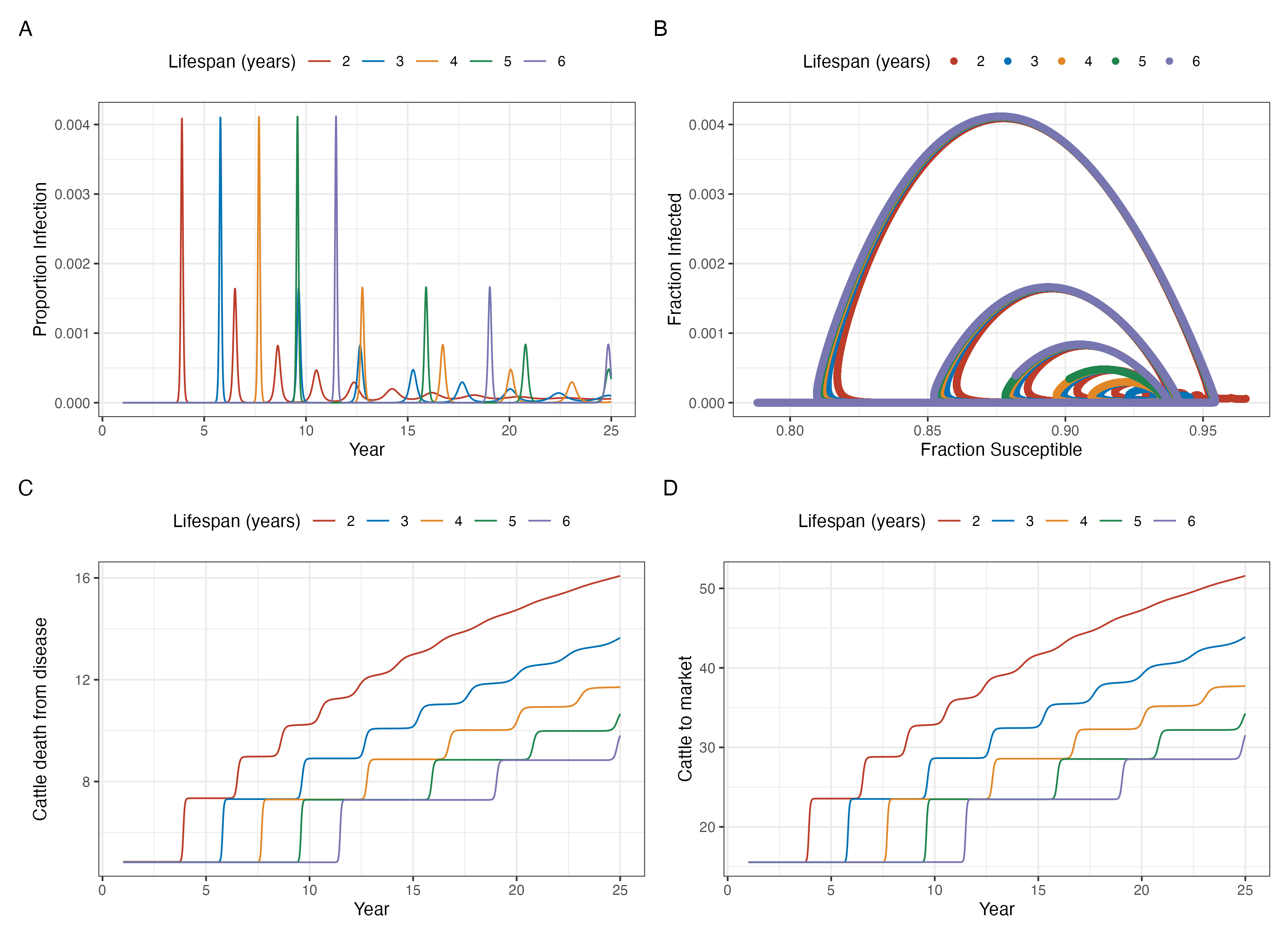

### fig_choropleth-1.png

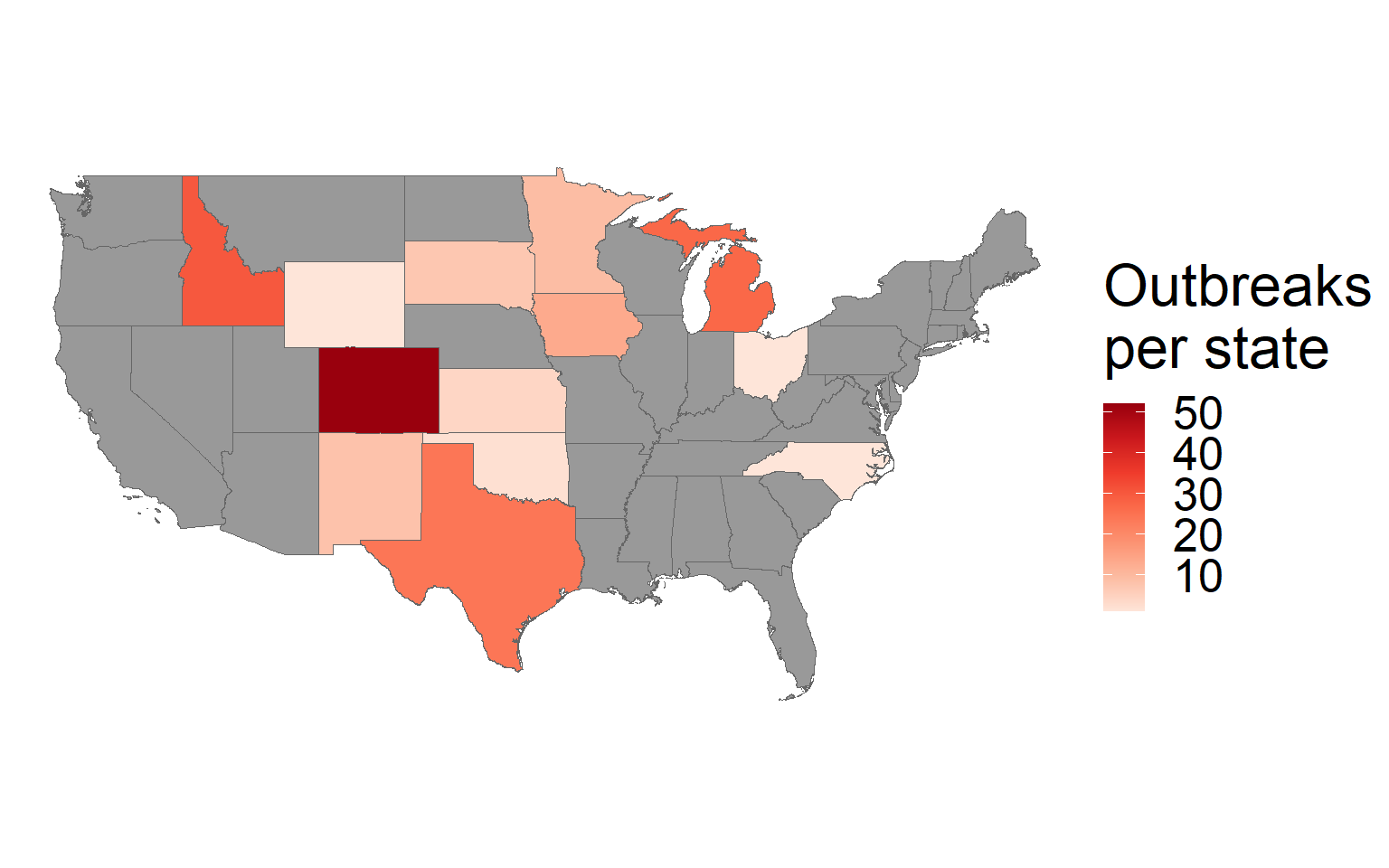

### fig_densitymap-1.png

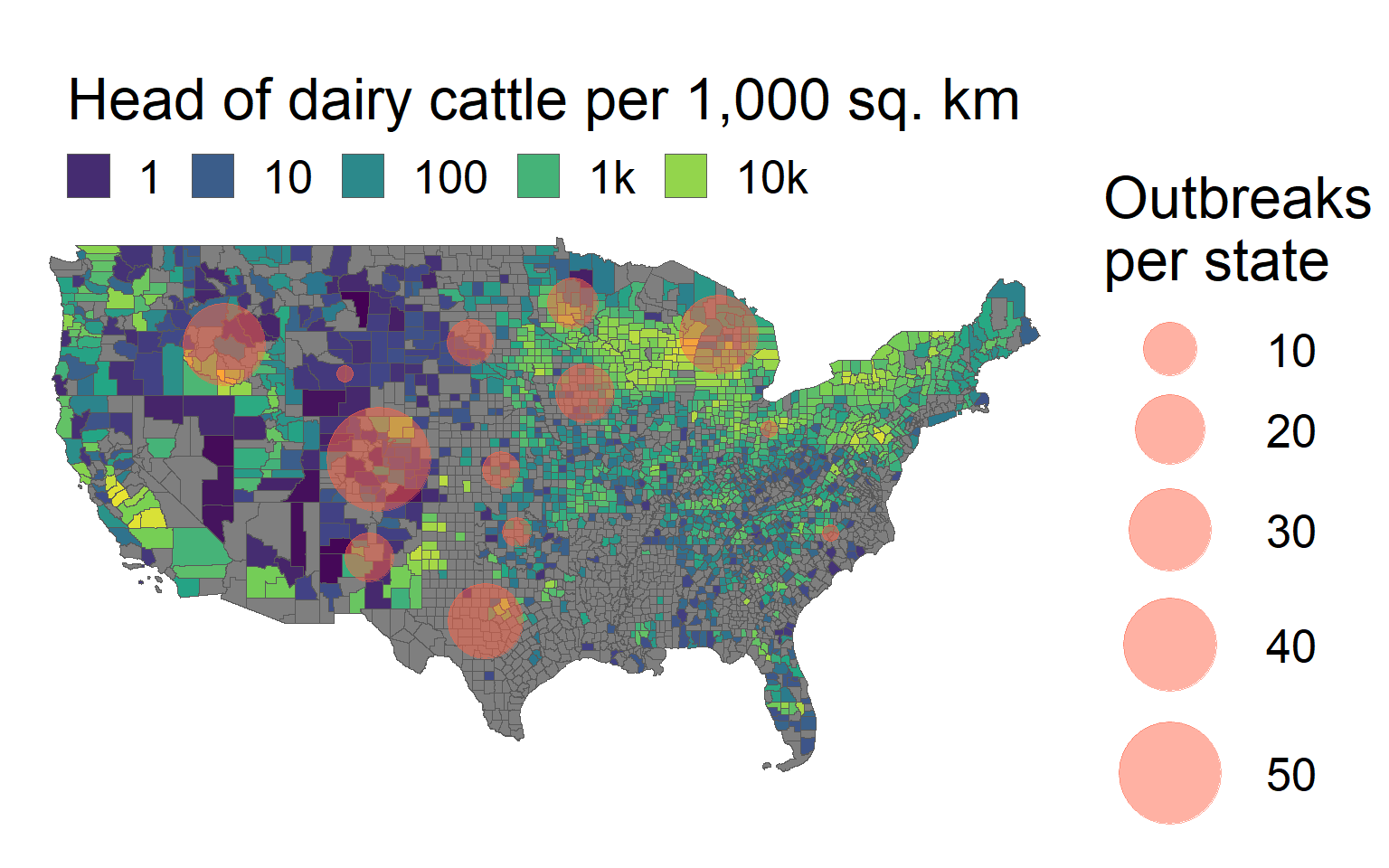

### fig_epicurves-1.png

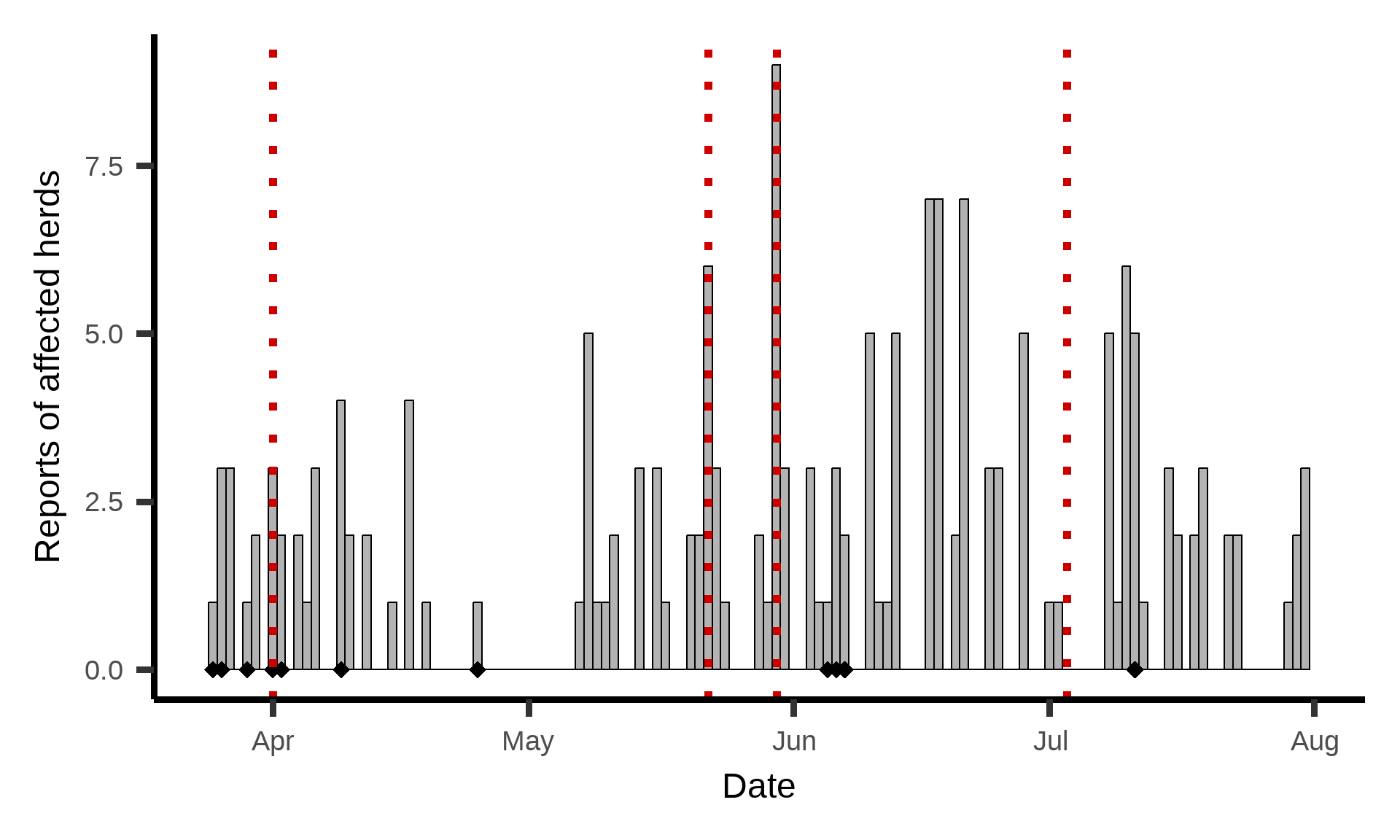

### fig_modeloutput-1.png

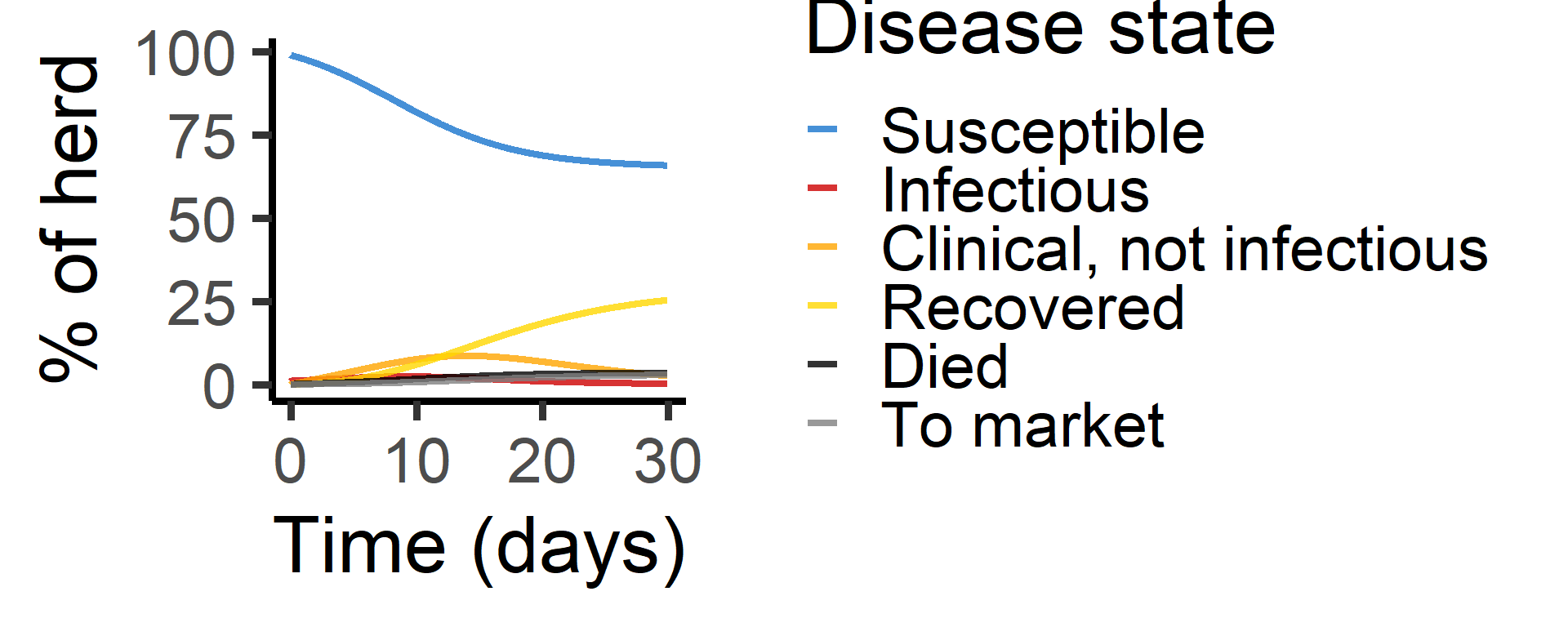
